# Individual differences in wellbeing are supported by separable sets of co-active self- and visual-attention-related brain networks

**DOI:** 10.1101/2023.08.29.552993

**Authors:** Yumeng Ma, Jeremy I Skipper

## Abstract

How does the brain support ‘wellbeing’? Because it is a multidimensional construct, it is likely the product of multiple co-active brain networks that vary across individuals. This is perhaps why prior neuroimaging studies have found inconsistent anatomical associations with wellbeing. Furthermore, these used ‘laboratory-style’ or ‘resting-state’ methods not amenable to finding manifold networks. To address these issues, we had participants watch a full-length romantic comedy-drama film during functional magnetic resonance imaging. We hypothesised that individual differences in wellbeing measured before scanning would be correlated with individual differences in brain networks associated with ‘embodied’ and ‘narrative’ self-related processing. Indeed, searchlight spatial inter-participant representational similarity and subsequent analyses revealed seven sets of co-activated networks associated with individual differences in wellbeing. Two were ‘embodied self’ related, including brain regions associated with autonomic and affective processing. Three sets were ‘narrative self’ related, involving speech, language, and autobiographical memory-related regions. Finally, two sets of visual-attention-related networks emerged. These results suggest that the neurobiology of wellbeing in the real world is supported by diverse but functionally definable and separable sets of networks. This has implications for psychotherapy where individualised interventions might target, e.g., neuroplasticity in language-related narrative over embodied self or visual-attentional related processes.

## Introduction

‘Wellbeing’ is a multidimensional construct, without a single definition^1,2^. It is often viewed from the hedonic or ‘subjective wellbeing’ perspective, characterised as ‘a person feeling and thinking’ their ‘life is desirable regardless of how others see it’^3,4^. This view contains both subjective emotional/affective (i.e., feeling) and evaluative/cognitive (i.e., thinking) components^3,5^. It is sometimes contrasted with a eudaimonic or ‘psychological wellbeing’ perspective that is more associated with the meaning, purpose, and self-realisation components of human activity^6–8^. Though these general characterizations grossly capture unique components of wellbeing, they are also highly interrelated^9,10^.

Reflecting the multidimensional nature of wellbeing, there are more than 100 self-report measures that are used to evaluate about 200 different dimensions^2,11^. These measures reflect different perspectives, like those above, sometimes with no overlap on any dimension measured^11–14^. Nonetheless, they capture something important as they have variously been used to show that wellbeing is associated with better physical health and decreased mortality^15–19^, higher chances of recovery from serious mental illnesses like schizophrenia or bipolar disorder^20,21^, increased productivity and higher job performance^22–24^, and improved social relationships^25^.

As might be expected from a multidimensional construct, these self-report measures correlate wellbeing with a large array of individual differences. To give a non-exhaustive overview, wellbeing is associated with individual differences in hearing^26^, interoception^27,28^, visual attention^29^, emotional intelligence^30,31^, up-regulating positive emotions^32^, following feelings^33^, cognitive emotional regulation strategies^34,35^, facets of mindfulness^36,37^, rumination^38^, finding meaning^39^, narratives about self-growth^40^, the abstractness versus concreteness of self-construal^41^, degree of past, present, and future thinking^42^, nostalgia^43^, authenticity^44^, ‘mental toughness’^45^, personality^46^, ethical orientation^47^, and social participation and outgoingness^48,49^.

### Neurobiology of Wellbeing

Despite its importance, our understanding of the neurobiology of wellbeing is limited. Results have been highly variable, with arguably ‘no consistent associations’ between specific brain regions and wellbeing across studies^50–52^. Reviews of the 20 to 60 neuroimaging studies related to wellbeing show that activation spans much of the brain, though not reproducibly^50–55^. We suggest that there are a few interrelated reasons for this. The first is implicit in the above, i.e., the multidimensional nature of wellbeing and the individual differences mirroring this dimensionality. This implies that there will be multiple underlying, co-active, and interacting brain networks supporting the manifestation of wellbeing distributed throughout the whole brain. Furthermore, these networks would be expected to be different or weighted differently depending on individual differences.

Yet, the paradigms used in neuroimaging studies of wellbeing have likely ensured variability of results across studies while not being able to find these different networks or weightings within any one study. Specifically, about half of existing studies can be characterised as ‘laboratory-style’, involving reductionist paradigms with relatively unnatural but well controlled stimuli and tasks^51^. Stimuli include affective pictures, music, playing cards, robot arms, sentences, and video games, among others. Tasks involve emotional reactivity, facial recognition, gender discrimination, imagination, monetary rewards, repetition detection, etc. The remaining studies use a ‘resting-state’ paradigm with no controlled stimuli or tasks^51^. Participants might be trying to stay awake, fixating cross-hairs, trying not to think, visualising, and/or silently talking to themselves^56,57^. This heterogeneity of stimuli and tasks in laboratory-style and behaviours in resting-state studies likely produces anatomically variable results across studies.

Furthermore, neither paradigm as employed is amenable to finding the many regions and networks likely supporting wellbeing. Laboratory-style studies usually examined only affective perception and/or experienced affect, in exclusion of many other possible dimensions like the evaluative aspects of wellbeing^51^. Thus, the subtractive logic often employed could only possibly reveal ‘blobs’ related to those few dimensions, while obscuring associated networks (due to the region-focused methods) and completely hiding networks associated with dimensions not under study. When network analyses were employed in either paradigm, only a few networks could possibly be discovered because only one or a small number of seed regions of interest were used in nearly all studies^51^.

### Wellbeing and Self

What organising principles might be used to understand the whole-brain distribution of networks we propose to support individual differences in wellbeing? We and others have suggested that wellbeing is self-related and can be mapped onto self-related brain processes^58–60^. Indeed, self-report measures of wellbeing include self-related dimensions either implicitly (because all require self-related judgements) or explicitly, e.g., self-acceptance, self-care, self-confidence, self-control, self-discovery, self-efficacy, self-esteem, self-realisation, self-regard, and self-satisfaction^11^. Self-related processing encompasses large portions of the brain but might be grossly divided into those supporting ‘embodied’ and ‘narrative’ self-related processes^61–65^, though this is not to say these are also not intimately interrelated^66,67^.

The ‘embodied self’ refers to aspects of the self that are not or at least less obviously based on semantic representations and focus on the self in the present moment, like interoceptive awareness, the experience of body ownership, and emotions^68,69^. We suggest that this primarily links embodied self-related processes and supporting brain networks to individual differences in the emotional/affective (i.e., feeling) component of ‘subjective wellbeing’. For example, individuals better at interoception are better at emotional regulation and this is particularly reliant on the insula^70–75^. Therefore it can be expected that the insula predicts higher wellbeing in those people. Indeed, many of the regions involved in the embodied self^68^, nested in emotional processing more generally^76,77^, are also implicated in neuroimaging studies of wellbeing. In addition to the insula, these include the amygdala, anterior cingulate cortex, basal ganglia, dorso- and medial prefrontal cortices, and thalamus^50–55^.

In contrast to the embodied self, the ‘narrative self’ has more obvious semantic content that extends the present to past autobiographical memories and imagined futures^63^. Though it is almost never acknowledged, these narratives are typically constructed and manipulated with language^60^. We suggest that this links narrative self-related processes and associated brain networks to individual differences in the evaluative/cognitive (i.e., thinking) component of ‘subjective wellbeing’ and the meaning and purpose components of ‘psychological wellbeing’. Specifically, the words we use to describe our world, ourselves, and the meaning of our lives are contingent on our personal contexts and can become deeply entrenched through a lifetime of learning and repetition^60^. While this is often adaptive and beneficial, it can also have negative consequences, as in rumination via increased verbal thinking^78–80^. Therefore, it can be expected that language and autobiographical memory-related regions will predict individual differences in wellbeing more in some people. Indeed, of all regions, the superior temporal gyrus (associated with language comprehension) and precuneus (associated with autobiographical memory) are often the most predictive of mental health and wellbeing (for a review, see^60^). These regions also regularly appear in neuroimaging studies of wellbeing^50–55^.

### Study Overview

In summary, existing work on the neurobiology of wellbeing has yielded variable results, likely because ‘wellbeing’ is multidimensional and subject to individual differences. Furthermore, the neuroimaging methods used have likely contributed to this variability and are not conducive to discovering the multiple co-activated real-world networks that likely underlie these multiple dimensions. We proposed a general self-related framework for organising expectations about which of these co-activated networks might be associated with individual differences in wellbeing. Specifically, embodied self-related networks will be larger determinants of wellbeing in some while narrative self-related networks will be more important in other individuals.

To be able to test this framework, we developed methods capable of capturing individual differences in a construct that relies on multiple co-active real-world brain networks. This required a stimulus that engages a full range of networks in a manner that is similar to everyday life. We accomplished this by asking 20 participants to complete a wellbeing questionnaire and then watch the romantic comedy-drama film ‘*500 Days of Summer*’ during fMRI^81^. Next, we developed an analysis pipeline that could find corresponding individual differences in wellbeing and brain activity, while grouping this activity into networks and sets of co-active networks (Fig. 1).

**Figure 1.**
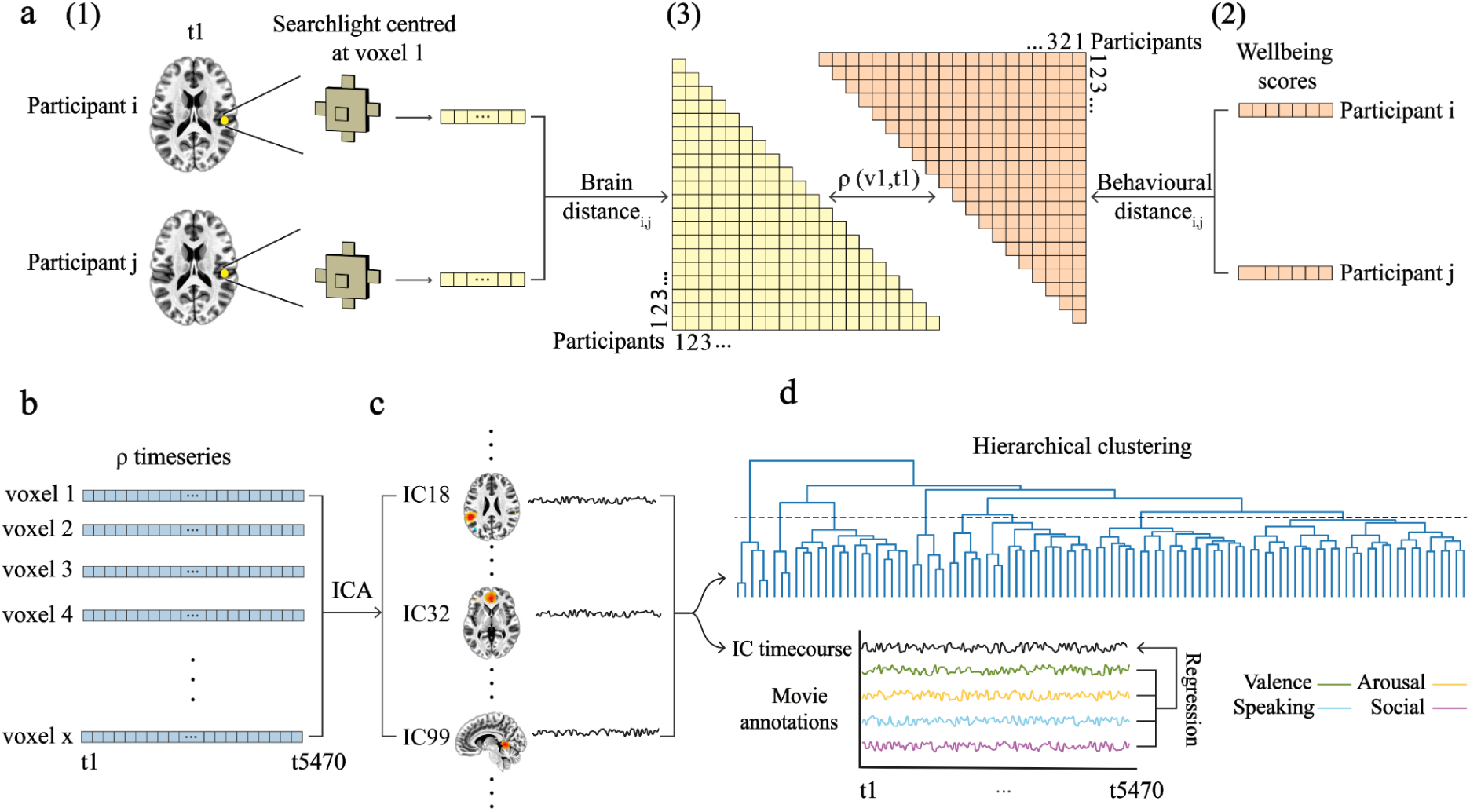
Analysis pipeline. a, Searchlight spatial inter-participant representational similarity analysis (SSIP-RSA). (1) For a specific time point (t1), the correlation distance of brain activity patterns inside a searchlight centred at a voxel (v1) with a radius of six mm (33 voxels) was calculated between each pair of participants (i, j). (2) The correlation distance of wellbeing score profiles was computed between each pair of participants (i, j). (3) Partial Spearman’s rank correlation (ρ) was calculated between participant-by-participant brain and behavioural distance matrices for v1 at t1, controlling for age and gender (not shown). b, The ρ value was calculated for each voxel at each of 5470 time points. c, Independent component (IC) analysis (ICA) was used to decompose the whole-brain timeseries of ρ values into 100 IC brain maps and corresponding time courses. d top, These IC time courses were hierarchically clustered and the resulting dendrogram was cut at the level of eight clades (dotted line). IC maps in each clade were thresholded with an activation probability larger than 0.5 calculated using a Gaussian Mixture Model and cluster sizes >50 voxels. Resulting IC maps from the same clade were merged, leaving seven co-activated networks (shown in Figs. 3-5 and Fig. 7). d bottom, Multiple linear regression was also performed on the IC time courses. The latter were treated as the dependent variables to regress against film annotations. Two low-level annotations were also included as controls in the regression model (not shown).

First, ‘searchlight spatial inter-participant representational similarity analysis’ (SSIP-RSA)^82–84^ determines how variations in brain activity patterns across participants in a searchlight at each time point in the film relates to variations in wellbeing as measured by the NIH Toolbox^85^ (Fig. 1a and 1b). Second, independent component analysis (ICA) decomposes the resulting timeseries into 100 spatial independent components (ICs), i.e., networks, and associated timecourses, while reducing dimensionality (Fig. 1c). Third, hierarchical clustering of those timecourses is used to find sets of co-active networks and further reduce dimensionality (Fig. 1d, top). Finally, we quantify the functional properties of these sets of networks by spatially correlating them with 10 pre-chosen term-based neuroimaging meta-analyses from a database of thousands of meta-analyses (i.e., Neurosynth)^86^.

We hypothesised that some sets of co-active networks representing individual differences in wellbeing would be more associated with terms we selected to correspond to embodied self-related processing. These are the ‘autonomic’, ‘affective’, ‘emotion’, ‘interoceptive’, and ‘self referential’ meta-analyses and associated regions (e.g., the insula and anterior cingulate cortex). Another set of networks was hypothesised to be associated with narrative self-related terms, including ‘autobiographical’, ‘language’, ‘memory retrieval’, ‘speech perception’, and ‘speech production’ meta-analyses and associated regions (e.g., the precuneus and superior temporal gyrus; see Decoding, Materials and Methods for justification of terms). To corroborate results, we also provide the top 10 correlations for resulting networks and sets of co-active networks with all neuroimaging meta-analyses in the database. Finally, we conducted an exploratory analysis to determine if individual differences in embodied and narrative self-related networks dynamically scale with socio-affective film context by regressing annotations against IC timecourses (Fig. 1d, bottom).

## Results

We hypothesised that individual differences in wellbeing would be correlated with individual differences in whole brain sets of networks associated with embodied and narrative self-related processing. Wellbeing scores from the wellbeing subdomain of the emotional battery included in the NIH Toolbox were well distributed (Fig. 2)^87^. SSIP-RSA using these scores followed by ICA and hierarchical clustering (see Fig. 1a-d) resulted in eight sets of co-activated networks or clades (where a clade is a distinct group of ICs that share a common point of origin on the dendrogram; Fig. 1d and 3a). One of these clades was excluded from further analysis because it did not survive cluster size thresholding. Only ICs corresponding to statistically significant individual differences as determined by a Mantel test and corrected for multiple comparisons are included in clades (all FDR-corrected ps < 0.01; cluster sizes > 50 voxels). In what follows, we present clades ordered thematically by the functional properties associated with each clade (Figs. 3-5; Fig. 7a; Supplementary Table S1).

**Figure 2.**
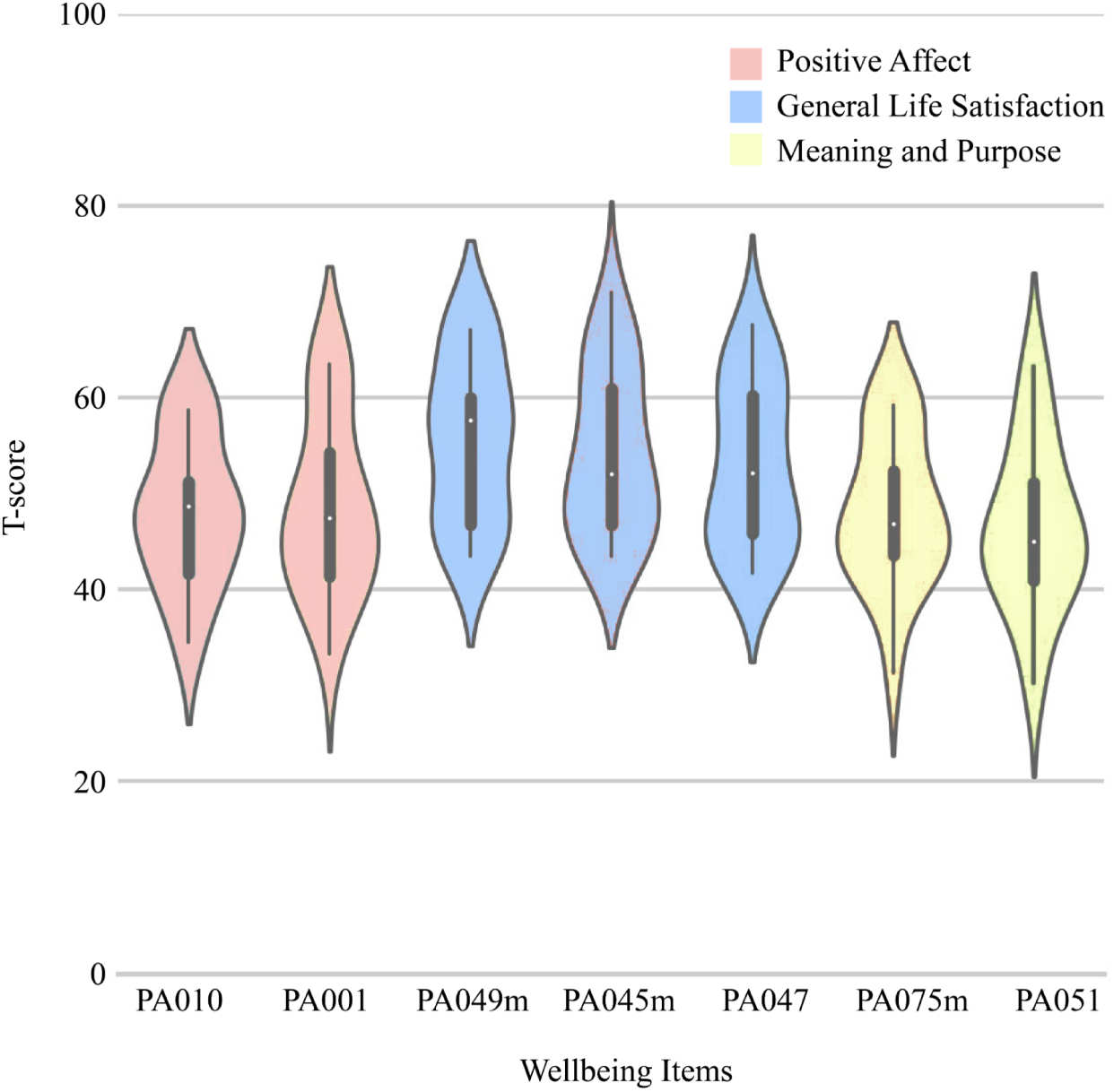
Wellbeing scores. Distributions of scores across 20 participants on seven items from the wellbeing subdomain of the NIH toolbox. Items comprising this subdomain are from the Positive Affect, General Life Satisfaction, and Meaning and Purpose questionnaires. The x-axis labels are specific item names from these questionnaires. The y-axis shows T-scores, where 50 is the mean of the US general population and 10 units is the standard deviation. The boxplot in the middle of each violin plot is the maximum, third quartile, median (white dot), first quartile, and minimum T-score. Width of the density plot represents frequency.

### Networks Sets 1/2

The co-activated networks in the first two clades are likely functionally related in that they are clustered next to each other (Fig. 3a, right). That is, they have similar timeseries suggesting that the information being processed in associated brain regions is thematically related. Supporting this, both clades loaded on different networks that were functionally associated with embodied self-related processing terms. For clade one, these most prominently involved medial frontal cortices anterior to and avoiding the precuneus (Fig. 3b, left). It included the anterior and midcingulate and dorsal and ventral medial frontal cortices. Other regions were the right insula and right cerebellum. Collectively, this clade was most correlated with the ‘affective’, ‘autonomic’, and ‘self referential’ neuroimaging meta-analyses (all rs ≥ 0.10, Fig. 3b, right). The second clade consisted predominantly of the bilateral insula and subcortical regions including the pallidum, putamen, thalamus, and ventral tegmental area (Fig. 3c, left). These regions were most correlated with the ‘affective’, ‘autonomic’, ‘interoceptive’, and ‘speech production’ meta-analyses (all rs ≥ 0.11, Fig. 3c, right). For both clades, results were supported by a more data-driven approach, examining the top 10 neuroimaging meta-analytic associations. That is, both clades had functionally overlapping correlations with the ‘mood’, ‘pain(ful)’, and ‘reward’ term-based meta-analyses (Supplementary Table S1). Clade one was more associated with, e.g., ‘emotional’ and ‘valence’ whereas clade two was more associated with ‘reinforcement’ and ‘reward’ overall (Supplementary Table S1).

**Figure 3.**
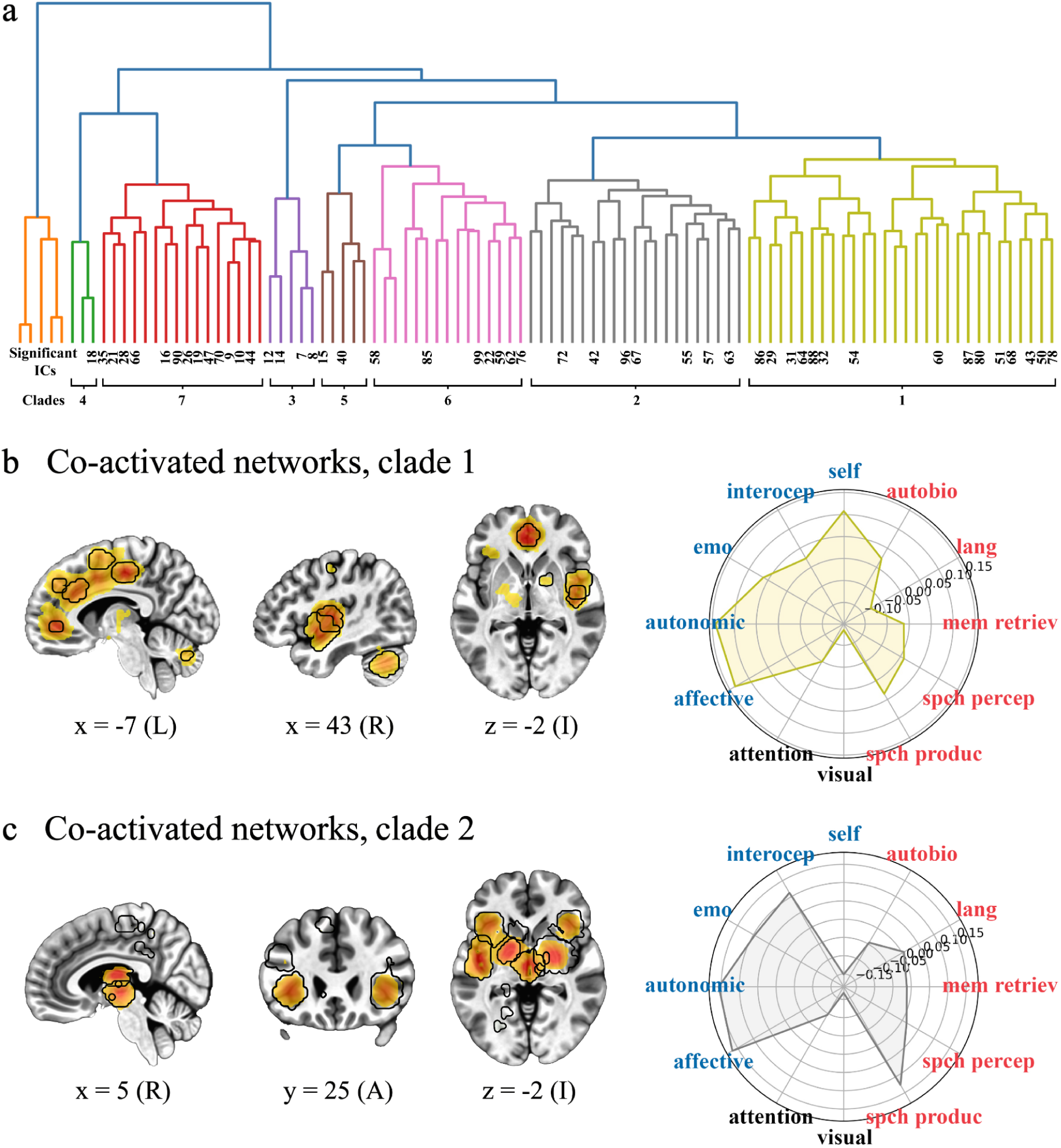
Network sets 1/2. Co-activated networks correlated with individual differences in wellbeing and associated with embodied self-related processing. a, The dendrogram from Fig. 1 is coloured by clade. b, Representative brain slices showing all independent components (ICs) from clade 1. c, Representative brain slices showing all ICs from clade 2. Activation on slices in a and b is coloured by z-scores, where yellows represent lower values and red represents higher values (≤ 15.00). Regions outlined in black show statistically significant individual differences as determined by a Mantel test (false discovery rate corrected p < 0.01, one-tailed). Numbers below slices are MNI coordinates, where x is L/R (left/right), y is A/P (anterior/posterior), and z is I/S (inferior/superior). Polar plots are the results of hypothesis-based correlations between co-activated network maps and term-based meta-analyses. Axis are correlation values with the internal area coloured by clade. Terms in red are related to narrative self-related processing, i.e., autobio (autobiographical), lang (language), mem retriev (memory retrieval), spch percep (speech perception), and spch produc (speech production). Terms in blue are related to the embodied self-related processing, i.e., affective, autonomic, emo (emotion), interocep (interoceptive), and self (self referential). The ‘visual’ and ‘attention’ terms in black are included for comparison. See Fig. 7a, bottom row for a three-dimensional whole-brain representation of the activation patterns in b and c.

### Network Sets 3/4/5

The co-activated networks in clades three through five are likely functionally related in that clades three and five are next to each other in the clustering dendrogram and all three clades are near each other on the left side of the dendrogram (Fig. 3a). Supporting this, clades, three, four, and five all loaded on regions functionally related to our definition of the narrative self-related processing. Specifically, the third clade consisted of the right Heschl’s gyrus, right superior temporal gyrus and left supramarginal gyrus (Fig. 4a, left). The clade as a whole was correlated with the ‘language’, ‘speech perception’, and ‘speech production’ meta-analyses (all rs ≥ 0.17, Fig. 4a, right). The fourth clade comprised the right ventral precentral gyrus (Fig. 4b, left) and correlated with the ‘speech production’ meta-analysis (r = 0.19; Fig. 4b, right). The fifth clade contained left posterior medial precuneus regions (Fig. 4c, left). These regions were most correlated with ‘attention’ and ‘memory retrieval’ meta-analyses (all rs ≥ 0.10, Fig. 4c, right). Again, results are supported by terms derived from the more data-driven approach. In addition to those mentioned, these included language-related terms like ‘articulatory’, ‘linguistic’, ‘phonological’, ‘sentence(s)’, ‘speech perception’, ‘production’, and ‘vocal’ and memory-related terms like ‘autobiographical’, ‘episodic memory’, and ‘memory retrieval’ (Supplementary Table S1). Collectively, these clades contained the IC that was the most correlated with individual differences across this entire study, located in the left supramarginal gyrus (ρ = 0.53). Furthermore, clades three, four, and five were the most predictive of individual differences on average (mean ρ = 0.38 versus 0.29 for clades 1/2 and 0.30 for clades 6/7; Supplementary Table S1).

**Figure 4.**
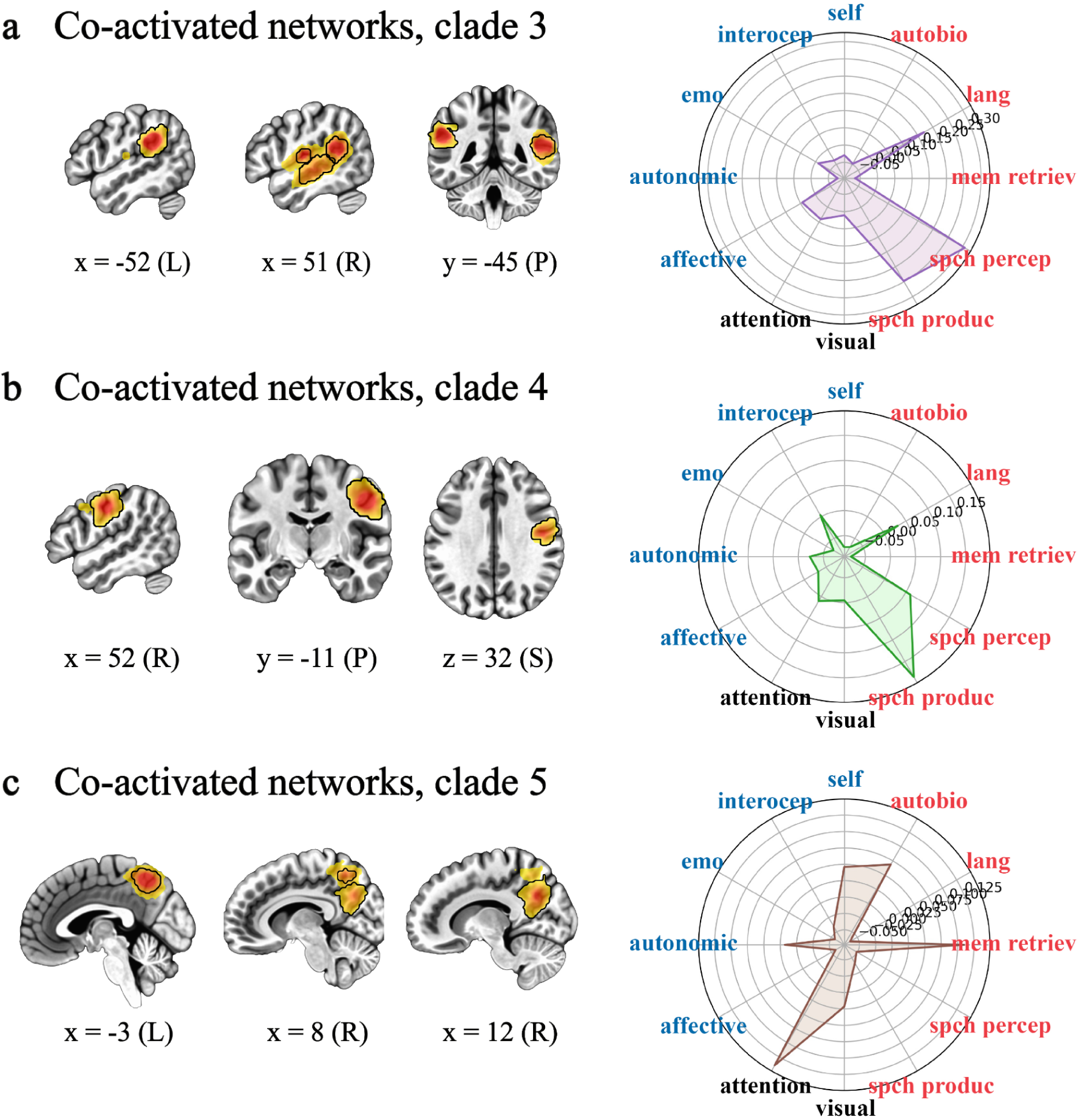
Network sets 3/4/5. Co-activated networks correlated with individual differences in wellbeing and associated with narrative self-related processing. a, Representative brain slices and polar plot from clade three (a), clade four (b), and clade five (c). See Fig. 3 for details and abbreviations and Fig. 7a, top row for a whole-brain representation of the activation patterns in a-c.

### Network Sets 6/7

The final two clades were functionally related but do not necessarily conform to any a priori hypotheses pertaining to self-related processing. Clade six showed sets of networks that included regions in the occipital lobe like the calcarine sulcus and lingual gyrus (Fig. 5a, left) whereas clade seven mainly consists of regions in the dorsolateral prefrontal cortex, bilateral intraparietal sulci, and middle temporal cortices (Fig. 5b, left). Supplementary Table S1 shows that both sets of networks are associated with an array of visual and visual attention-related meta-analyses. For these reasons, we included ‘visual’ and ‘attention’ in the polar plots in Figs. 3-5 for purposes of comparison. Clade six is correlated with the ‘visual’ (r = 0.20) and ‘autobiographical’ (r = 0.11) and clade seven with the ‘attention’ (r = 0.42) and ‘visual’ (r = 0.29) meta-analyses. Other meta-analytic terms included ‘eye fields’, ‘object(s)’, and ‘visual stream’ for clade six and ‘attentional’, ‘visual motion’, and ‘visuospatial’ for clade seven.

**Figure 5.**
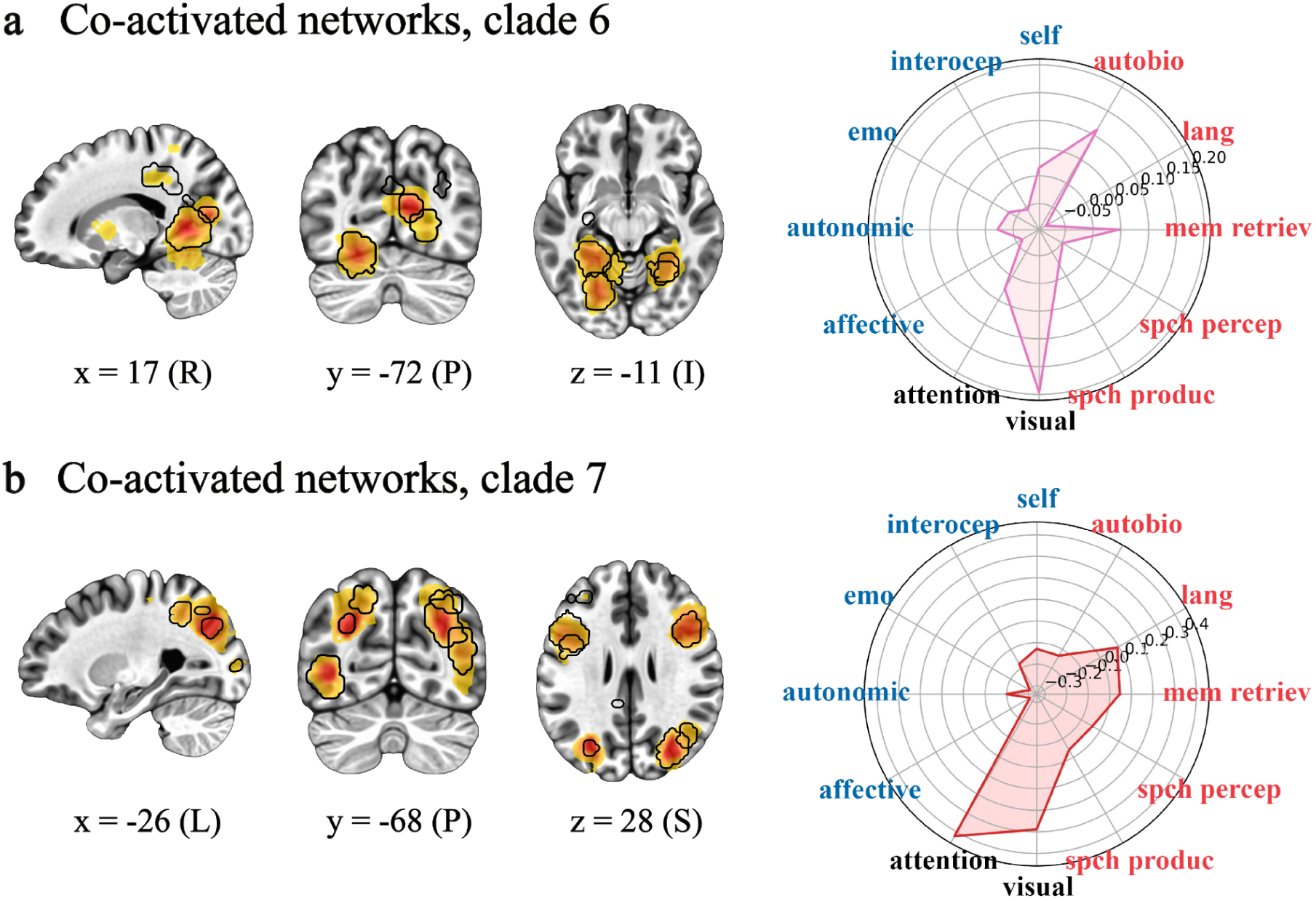
Network sets 6/7. Co-activated networks correlated with individual differences in wellbeing and associated with visual-attention-related processing. Representative brain slices and polar plots from clade six (a) and clade seven (b). See Fig. 3 for details and abbreviations and Fig. 7a, middle row for a whole-brain representation of the activation patterns in a and b.

### Socio-Affective Contexts

Finally, we conducted an exploratory regression analysis to determine whether fluctuations in brain networks that correlated with individual differences are related to socio-affective contexts in ‘*500 Days of Summer*’. Resulting positive and negative associations would suggest the degree to which networks are influenced by context in the film whereas a lack of association might suggest the relative impermeability to context. To examine this, we regressed valence, arousal, social interaction, and people speaking film annotations and two nuisance regressors generated by Masson and Isik^88^ against all IC timecourses (see Fig. 1d). There were no statistically significant results after correcting for multiple comparisons at p < 0.01 (the threshold we applied throughout). At a less conservative though still corrected threshold, there were 11 components that scaled with socio-affective contexts (all FDR-corrected ps < 0.05; Fig. 6).

**Figure 6.**
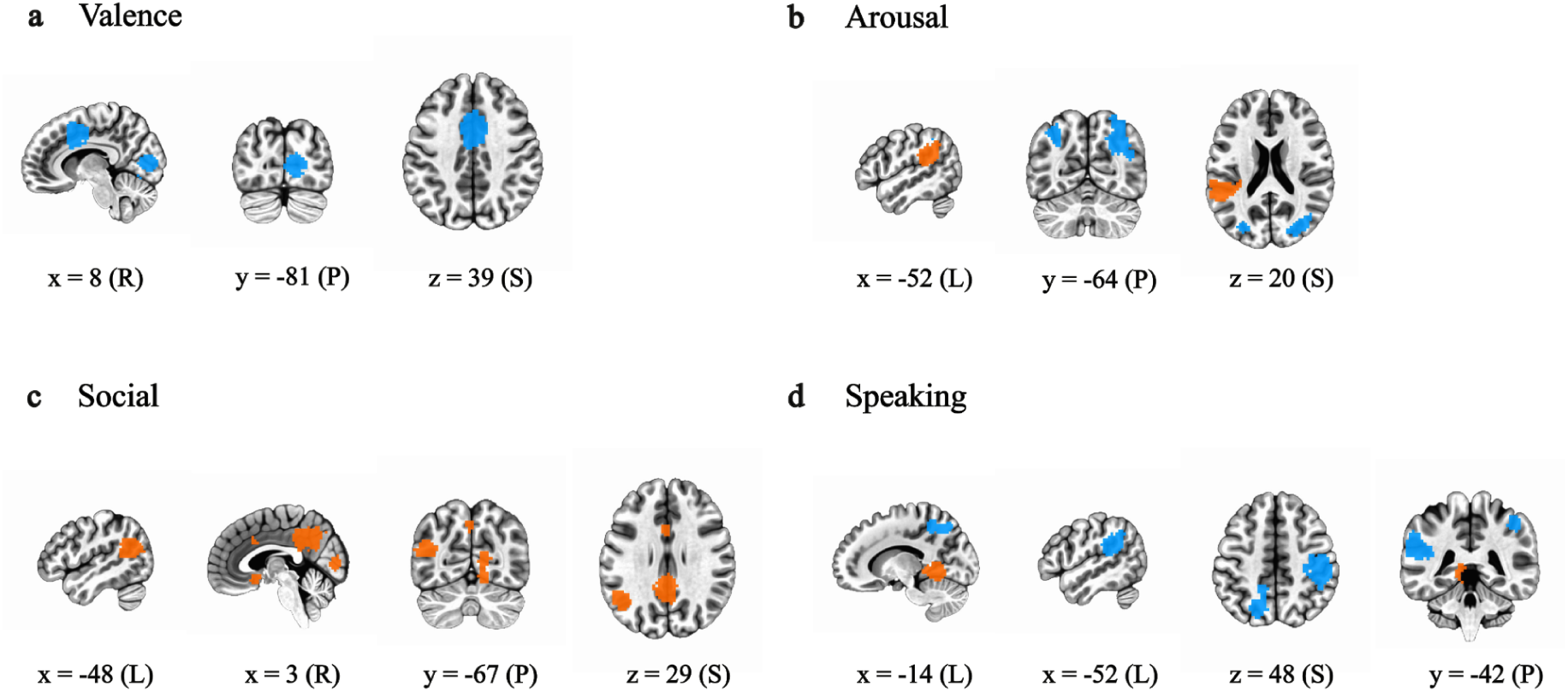
Socio-affective contexts. Multiple linear regression of independent component (IC) time courses against the film annotations for valence (a), arousal (b), presence of social interaction (c), and presence of people speaking (d). Shown are thresholded IC maps with time courses that have a statistically significant relationship with film annotation timeseries (false discovery rate corrected p < 0.05, two-tailed permutation, all clusters > 50 voxels). Positive and negative βs are shown in orange and blue, respectively. Numbers are MNI coordinates of slices, where x is L/R (left/right), y is A/P (anterior/posterior), and z is I/S (inferior/superior).

Specifically, networks with individual differences that varied with valence in the film are the right calcarine sulcus (IC 5; β = −0.085, p < 0.0001, uncorrected) and anterior midcingulate cortex (IC 31; β = −0.050, p = 0.001, uncorrected; Fig. 6a). Networks varying with arousal are the left (IC 21, β = −0.065, p < 0.0001, uncorrected) and right (IC 9; β = −0.062, p < 0.0001, uncorrected) intraparietal sulci and left supramarginal gyrus (IC 8; β = 0.061, p = 0.0002, uncorrected; Fig. 6b). Networks varying with the presence of social interactions are the right calcarine sulcus (IC 5; β = 0.063, p = 0.0004, uncorrected), posterior cingulate cortex (IC 52; β = 0.049, p = 0.0006, uncorrected), and midcingulate cortex, subgenual anterior cingulate cortex, and temporoparietal junction (IC 81; β = 0.049, p = 0.0012, uncorrected; Fig. 6c). Finally individual difference networks varying with the presence of people speaking are the bilateral intraparietal sulci (IC 35; β = −0.050, p = 0.0007, uncorrected), left lingual gyrus (IC 99; β = 0.046, p = 0.0012, uncorrected), and left supramarginal gyrus (IC8; β = −0.056, p = 0.0004, uncorrected; Fig. 6d). Thus, a full 55% of ICs (i.e., 6/11) are from clades six and seven associated with visual attention related processing (see Supplementary Table S1). Furthermore, 100% of regressors (4/4) were associated with an IC from these visual-attention-related clades.

## Discussion

Because it is a multidimensional construct and due to the variability of past neuroimaging results, we proposed that wellbeing is supported by a whole-brain distribution of co-active networks, subject to individual differences in weighting. We suggested that these might be broadly understood as being organised around two aspects of self-related processing. Sometimes wellbeing hinges on judgements about physical, interoceptive, and emotional states, i.e., embodied self-related processing. At others, wellbeing is associated with judgments around the effectiveness of the typically language-based stories we tell ourselves, i.e., narrative self-related processing. We evaluated this framework by assessing wellbeing and then asking participants to watch a romantic comedy-drama film during fMRI. Using SSIP-RSA combined with ICA and hierarchical clustering (Fig. 1), we confirmed hypotheses that individual differences in wellbeing (Fig. 2) were correlated with individual differences in sets of co-activated networks, divisible into embodied and narrative self-related processing (Figs. 3-4; Fig. 7a, top and bottom rows; Supplementary Table S1). We discovered a third set of visual-attention-related co-activated networks that also correlated with individual differences in wellbeing (Fig. 5; Fig. 7a, middle row; Supplementary Table S1). Dynamically varying socio-affective context from the film scaled primarily with individual regions from these visual-attention-related networks (Fig. 6).

**Figure 7.**
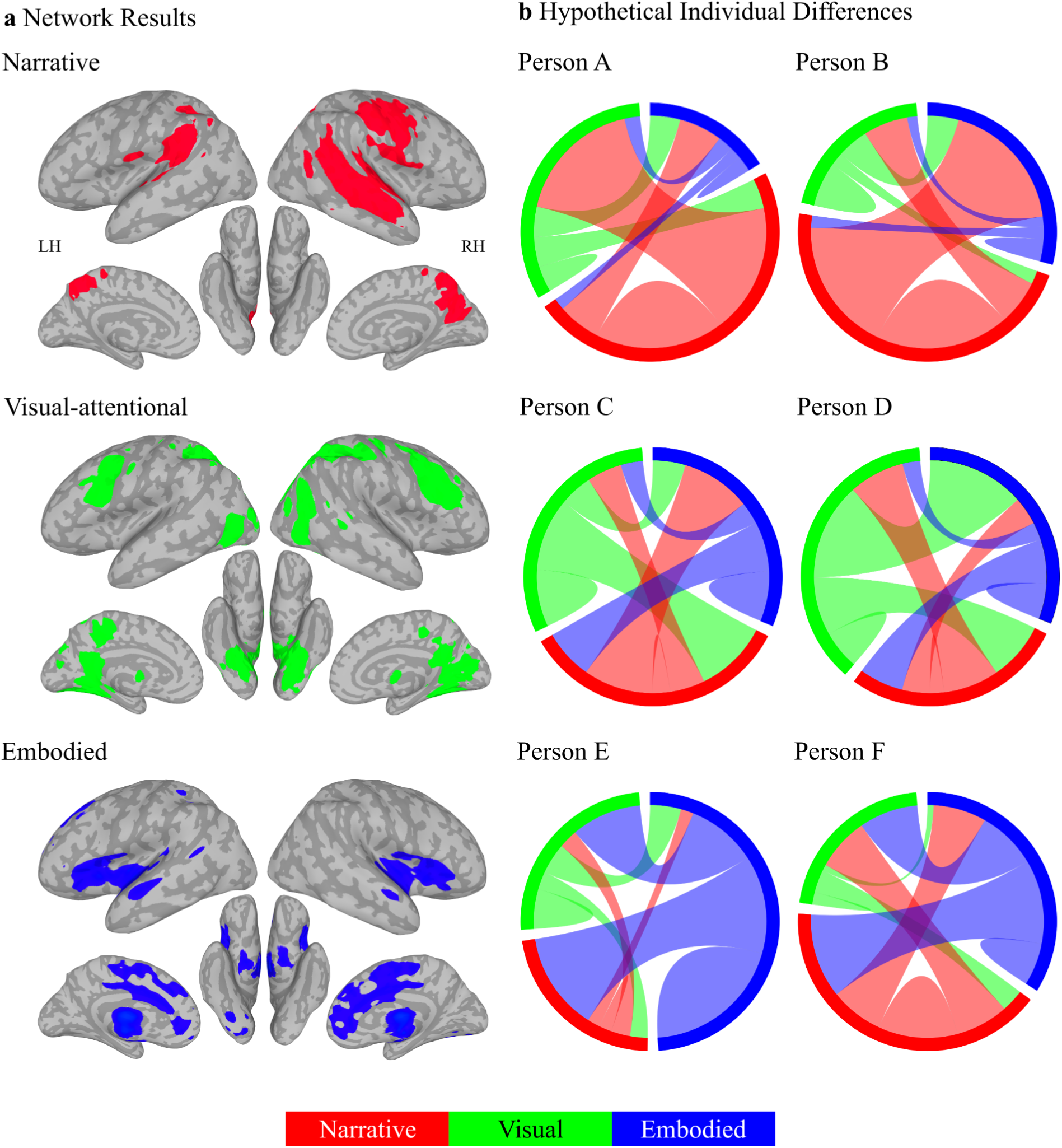
Multiple selves model of individual differences in wellbeing. (a) We propose the ‘self’ is a whole-brain process, broadly organised into three dynamically interacting self-related networks, each with multiple subnetworks (left column). These networks, from Figs. 3-5, are projected onto inflated MNI surfaces, with lighter and darker greys indicating sulci and gyri, respectively. Colours indicate activity in co-activated narrative (a, top, red; see Fig. 4), visual-attentional (a, middle, green; see Fig. 5), and embodied (c, bottom, blue; see Fig. 3) self-related networks linked to individual differences in wellbeing. (b) Each cord plot illustrates an individual’s hypothetical connectivity for these (sub)networks (Persons A-F), with cord width indicating strength and possible entrenchment of connections within and between network configurations, determining wellbeing. The wellbeing of Persons A and B is determined more strongly by narrative self-related networks (b, top, red) compared to Persons C/D (b, middle, green) and Persons E/F (b, bottom, blue), whose wellbeing is more determined by visual-attentional and embodied self-related networks, respectively. People may have different connectivity to different self-related networks; for example, although narrative self-related networks drive the wellbeing of Persons A and B, they have different connectivity to visual-attentional (b, top, red-to-green, Person A) and embodied (b, top, red-to-blue, Person B) self-related processes.

### Embodied Self

We defined embodied self-related processing as encompassing interoceptive awareness, body ownership, and emotional processing. Extensive behavioural and neurobiological research links these processes with mental health and wellbeing^68,71,89–96^. Consistent with this, we found that some individual differences in wellbeing were correlated with individual differences in brain activity in two closely related sets of co-activated networks that were associated with interception (pain and autonomic processes) and affective/emotional processing (Fig. 3; Fig. 7a, bottom row; Supplementary Table S1). Both sets included most of the insula bilaterally, regions consistently associated with interoceptive and emotional awareness^70,72–75,97,98^. The first set of co-activated networks additionally contained an array of medial frontal regions anterior to the precuneus, like the anterior cingulate cortex (Fig. 3b). These regions are collectively involved in a number of specific functions closely associated with emotional processing, particularly in more anterior portions^99,100^. They also play an important role in self-related processing in general and self-related emotional processing in particular^76,77,101,102^. In addition to the insula, the second set of co-activated networks included the pallidum, putamen, thalamus, and ventral tegmental area (Fig. 3c). These subcortical regions are primary components of reward processing in the brain^103,104^ and have also been linked to self-related processing^105^. The close functional relationship between these two sets of co-activated networks is consistent with recent discussions regarding the interrelationship between emotional and reward processing^106,107^.

### Narrative Self

We defined narrative self-related processing as being primarily constructed from language, inner speech, and autobiographical memory and associated brain regions^60^. Though research into these relationships is limited, what exists links these processes with mental health and wellbeing^60,78,108–117^. Consistent with this, we found that individual differences in wellbeing were correlated with three sets of co-activated networks that are associated with language, speech perception and production, and autobiographical memory-related processing (Fig. 4; Fig. 7a, top row; Supplementary Table S1). These narrative self-related networks contained the most predictive single brain region and were collectively the most predictive of individual differences in wellbeing in this study.

Regions in these sets of co-activated networks included the superior temporal, supramarginal, and ventral precentral gyrus that are variously involved in language comprehension, speech perception, and speech production, including inner speech^60,118–120^(Fig. 4a and 4b). They also included medial brain regions. Unlike the anterior distribution in embodied self-related networks, these were posterior, including aspects of the posterior cingulate and precuneus (Fig. 4c). Research pertaining to these regions suggests that they are associated with autobiographical memory, self-referential processing, and consciousness^121–123^. More generally, they are part of the larger ‘default mode network’ that has been argued to combine ‘episodic memory, language, and semantic memory processes to generate an ongoing internal narrative’^124^. Consistent with the various suggestions for anterior to posterior medial divisions, perhaps the more anterior medial regions are more associated with self-related emotional processing^100,121,125–130^. In contrast, we suggest that these more posterior regions are more associated with language/semantic aspects of self-related narrative processing^60,131,132^. This is supported by the close functional relationship between all three sets of co-activated networks, suggesting that the superior temporal, supramarginal, ventral precentral, and posterior medial regions all work together (see Fig. 3a).

We previously proposed a ‘HOLISTIC’ model to help account for these links, i.e., how the neurobiology of language and the default mode networks support both consciousness and wellbeing (aspects of which are depicted in Fig. 7)^60^. Roughly, the model suggests that our language and inner speech based narratives can become entrenched through learning and use within these language-related and default mode networks, making them more automatic and less accessible to conscious awareness. Though these entrenched narratives are typically adaptive, they might not always be appropriate or positive and may become less so over time, leading to decreased wellbeing. Therefore, the induction of neuroplasticity in these networks could lead to improved mental health and wellbeing. Indeed, changes to language-based narratives are arguably the primary target and mechanism of psychotherapy and perhaps even meditation^133^, leading to plasticity in language and default mode-related regions like the precuneus that often best predicts mental health and wellbeing outcomes^60^.

### Visual-Attention

We framed our hypotheses in terms of embodied and narrative self-related processing but unexpectedly found that individual differences in wellbeing are also correlated with individual differences in co-active networks associated with visual-attention-related processing (Fig. 5; Fig. 7a, middle row; Supplementary Table S1). One set of these networks included regions associated more with ‘lower level’ visual processing like primary visual cortex (Fig. 5a). The other set of co-active networks included more ‘higher level’ visual processing regions like bilateral middle temporal visual motion processing regions^134,135^. It also included much of the ‘dorsal attention network’ associated with top-down controlled attentional selection (e.g., the bilateral intraparietal sulcus; Fig. 5b)^136^. We also found that it was predominantly though not exclusively regions from these two sets of visual-attention-related networks that scaled with socio-affective context in the film (Fig. 6). However, the latter results should be interpreted with caution as no regions survived our initial p < .01 correction for multiple comparisons (though 11 survived at FDR-corrected p < .05).

These results suggest that individual differences in wellbeing might arise when people develop unique but stable attentional strategies for sampling that external world. This is consistent with research showing that mood impacts visual attention^29,137^ and sensory attentional biases are associated with mental health and contribute to wellbeing^138,139^. This warrants expanding the initial framework presented here to include three general neurobiological contributions to individual differences in wellbeing. These are the more internally focused embodied and narrative self-related networks and more externally focused and dynamically determined visual-attention-related networks. Perhaps these individual difference networks are more or less driven by the dominant formats that people experience their thoughts in. For example, experience sampling has been used to identify the ‘five frequent phenomena’ of inner experience^140^ and questionnaire measures have identified ‘modes’ of thinking^141^. From these, individuals more prone to ‘sensory awareness’/‘feeling’, ‘inner speaking’/‘internal verbalization’, or ‘inner seeing’/‘visual imagers’ might weight embodied self-related, narrative self-related, or visual-attention-related networks more on average, respectively^140,141^.

### Separability

Overall, results associate individual differences in wellbeing with separable embodied and narrative self- and visual-attentional related brain networks. Yet, functional descriptions of these networks do overlap somewhat. Most prominently, excluding the narrative self-related clades, 8.49% of the terms associated with the other clades were auditory and language-related (i.e., 27 of 318 total terms, with at least three terms in each clade, Supplementary Table S1). For example, both the embodied self-related processing clades resulted in anterior insula activity associated with, e.g., the terms ‘articulatory’ (clade one) and ‘phonological’ (clade two). The anterior insula is associated with both interoceptive processing and speech production, likely because autonomic/interoceptive processes like respiratory control are necessary for speech production and inner speech^60,142^. Thus, though these descriptions still fit within the general framework outlined here, they nonetheless suggest that clades found in this study are likely less functionally separable than they might appear.

Indeed, it would be nonsensical to argue that the underlying embodied and narrative self-related and visual-attention-related networks operate independently. Rather, we suggest they are generally interactive and integrative^60,143^. More specifically, the HOLISTIC model introduced above formalises this interactivity and suggests a rationale for the 8.49% prevalence of language-related terms in putatively non-language-related clades (Fig. 7)^60^. Namely, humans encounter and produce about 150,000 words a day. These constitute a primary tool for categorising our internal and external worlds, allowing us to extend our cognitive abilities by virtually manipulating these experiences with language. Neurobiologically, this can be accomplished because words are linked to interoceptive, emotional, and perceptual experiences in memory. Thus, when we experience emotions or colours, the associated labels are typically activated and vice versa. Because of this interdependence, words can impact wellbeing through entrenchment (as described above) but also through conscious manipulation. For example, labelling and reappraisal are primary tools for emotional regulation and are associated with the embodied, language/narrative, and visual-attention-related brain regions described above^144–146^. It is perhaps because of such interactivity that the narrative self-related co-active networks are the most correlated with individual differences in wellbeing.

### Limitations

A limitation of our initial embodied and narrative self-related framework is our assumption that participants are actually engaging in self-related processing, associating the film with their own experiences both implicitly and possibly explicitly (e.g., through inner speech). Indeed, it is inconceivable how the film could be understood if they did not and this contention is supported by both resting-state and naturalistic viewing research^147–152^. Nonetheless, because we did not explicitly test this assumption, alternative frameworks are possible. For example, results could be simply described in the more general terms associated with the resulting seven sets of co-active networks, i.e., as ‘emotion’, ‘reward’, ‘speech perception’, ‘speech production’, ‘memory’, ‘vision’, and ‘attention’. Alternatively, even more specific networks could be functionally described, down to the 100 sampled. Furthermore, the methodological decisions resulting in seven sets of co-active networks, derived from these 100 networks, were relatively arbitrary and perhaps even hundreds more networks might be described. Though this more agnostic and expansive approach is not wrong, it lacks overarching explanatory power for understanding the neurobiology of wellbeing as a whole. However, both perspectives are compatible with the HOLISTIC framework, which suggests that ‘the self’ is not confined to a single brain region or network. Instead, there are multiple dynamically organising and interacting self-related networks (as illustrated in Fig. 7). Though these are broadly described as narrative, visual, and embodied, our pluralistic view of the self can accommodate numerous additional self-related subnetworks (like those associated with different social roles). Strength and entrenchment within these networks, collectively determine individual differences in wellbeing.

## Conclusions

In this study, individual differences in wellbeing are shown to be supported by individual differences in three separable sets of co-activated networks associated with the embodied self, narrative self, and sensory-attentional related processing. These co-activated networks encompass much of the human brain, as might be expected from a multidimensional construct like wellbeing. They can be further subdivided into finer networks with more specific functions that predict individual differences in wellbeing. This holds promise for individualised approaches to medicine. For example, networks particularly associated with negative (and positive) wellbeing might be identified within an individual. These can then become the focus of individual or therapeutic interventions to increase wellbeing (e.g., through psychotherapy). This might take the form of inducing neuroplasticity through pharmacological means and/or manipulating interoception or visual attention more in some individuals or language-related narratives in others.

## Methods

This research made use of our ‘Naturalistic Neuroimaging Database’^81^. The NNDb includes data from 86 participants who completed the NIH Toolbox a few weeks before watching one of 10 previously unseen full-length films during fMRI. Here we analysed data from 20 participants who watched the 2009 romantic comedy-drama film ‘*500 Days of Summer’* (1 hour and 31 minutes).

### Participants

Of the 20 participants, 10 identified as female, with an age range of 19-53 years (M = 27.70, SD = 9.9). All were right-handed, native English speakers, with no history of neurological or psychiatric disorders, taking no medication, with no hearing impairments, and with unimpaired or corrected vision. The study was approved by the University College London Ethics Committee and all participants gave written informed consent. All procedures were performed in accordance with the Declaration of Helsinki.

### Wellbeing Measure

Participants completed the majority of the NIH Toolbox on an iPad about 2–4 weeks before fMRI scanning. The tests were administered in a sound shielded testing room and participants were provided with headphones. These included an emotional battery with a wellbeing subdomain that is composed of three components: positive affect, life satisfaction, and meaning and purpose^87^. Because not all participants answered the same questions, we included the seven items in the wellbeing subdomain with data from all 20 participants. These were two items from the ‘Positive Affect CAT Age 18+ v2.0’ (PA010 & PA001), three items from the ‘General Life Satisfaction CAT Age 18+ v2.0’ (PA049m, PA045m, & PA047), and two items from the ‘Meaning and Purpose CAT Age 18+ v2.0’ (PA075m & PA051). Raw scores were converted to standard T-scores where 50 represents the mean of the US general population (based on the 2010 Census) and 10 units represent one standard deviation. Scores from the seven items were well distributed, permitting further analysis (Fig. 2).

One limitation of this approach is that the low number of items in each wellbeing subdomain prohibited us from distinguishing between these subdomains neurobiologically. For example, we were unable to test whether the positive affect or meaning/purpose subdomains were more strongly associated with hypothesised embodied or narrative self-related brain networks, respectively. Another potential limitation is that the seven items from all 20 participants may not have been representative of the broader wellbeing measure. However, this is unlikely because almost all individual questions within the questionnaire are moderately to highly correlated^153^. To provide further validation, we correlated individual scores on the seven items with their corresponding full scores across participants. The seven-item average showed a high correlation with the full score (r = 0.95, p = 6.8×10^-11^). Individual items also exhibited high correlations, with r-values for PA010 = 0.88 (p = 2.8×10^-7^), PA001 = 0.88 (p = 4.0×10^-7^), PA049m = 0.69 (p = 0.00075), PA045m = 0.69 (p = 0.00081), PA047 = 0.79 (p = 3.9×10^-5^), PA075m = 0.75 (p = 0.00016), and PA051 = 0.82 (p = 1.0×10^-5^). These considerations suggest that the abbreviated wellbeing scores used here are an accurate reflection of the broader wellbeing construct.

### f/MRI Preprocessing

MRI data were obtained on a 1.5T Siemens MAGNETOM Avanto with a 32-channel head coil. Anatomical images were acquired using an MPRAGE sequence (TR = 2730 ms, TE = 3.57 ms, 176 sagittal slices, voxel size = 1 × 1 × 1 mm^3^). Functional images were acquired using a multiband echo planar imaging (EPI) sequence (TR = 1000 ms, TE = 54.8 ms, flip angle 75°, 40 interleaved slices, voxel size = 3.2 × 3.2 × 3.2 mm^3^, multiband factor = 4, no in-plane acceleration). The film ‘*500 Days of Summer*’ had 5470 whole brain volumes/TRs.

Unless otherwise noted, all preprocessing and data visualisation was done with the AFNI software package and specific programs are indicated where appropriate^154^ (https://afni.nimh.nih.gov/). First, anatomical images were corrected for intensity non-uniformity, deskulled, and nonlinearly aligned to an MNI template (i.e., MNI152_2009_template_SSW.nii.gz; using ‘@SSwarper’). Next, Freesurfer’s ‘recon-all’ was run with default parameters (version 7.0; http://www.freesurfer.net/)^155,156^. This allowed us to segment anatomical images into ventricle and white matter regions of interest to be used to make nuisance regressors (see next paragraph) and a group grey matter mask. The latter was created by resampling the anatomical images to functional resolution (‘3dresample’). From these, we removed the Freesurfer-derived ventricle and white matter regions of interest. We then took the union of these from all participants to create a group grey matter mask (using ‘3dMean’).

Preprocessing of the functional data was done using ‘afni_proc.py’ (https://afni.nimh.nih.gov/pub/dist/doc/program_help/afni_proc.py.html) and timeseries are available as derivatives on OpenNeuro (https://openneuro.org/datasets/ds002837/versions/2.0.0). Blocks were included for despiking (‘despike’), slice-timing correction (‘tshift’), volume registration to the run with the least number of outliers and warping to MNI space (in one step, ‘align’, ‘tlrc’, and ‘volreg’), masking (‘mask’), scaling to a mean of 100 (‘scale’), and regression (‘regress’). The latter used ‘3dDeconvolve’ to remove demeaned motion and motion derivatives, a polynomial degree of two, bandpass filtering (0.01 to 1), ventricular activity (first three principle components), and white matter activity from the timeseries on a per run basis in one step. Finally, as additional protection, hand-labelled ICA-based artefacts were also removed from the resulting timeseries (for details, see)^81^. Note that we did not smooth this data because the searchlight analysis uses a voxel radius of six mm.

### SSIP-RSA

We examined individual differences using searchlight spatial inter-participant representational similarity analysis (SSIP-RSA)^82–84^. This approach measures how matched inter-participant variability in fMRI activity patterns and wellbeing score profiles are in searchlights at each time point in the film (Fig. 1a). Specifically, we quantified inter-participant variability by computing a participant-by-participant distance matrix for both brain patterns (Fig. 1a, 1) and wellbeing scores (Fig. 1a, 2). Each square of the matrix is the correlation distance of the brain pattern vector or item-level wellbeing score vector between a pair of participants (i, j). The equation for the correlation distance is:

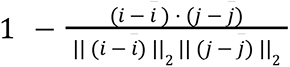

Here the brain distance matrix was computed time point by time point, using spatial patterns in searchlight spheres centred at each voxel with a radius of six mm (or 33 voxels). Unlike typical IP-RSA which uses temporal inter-participant correlation to measure brain synchrony of the whole timeseries, spatial inter-participant analysis allows us to calculate inter-participant variabilities dynamically at all 5470 time points in the film. This provides a foundation for the subsequent discovery of networks and co-active networks as well as the possibility to understand how film annotations are related to individual differences.

As controls, age and gender differences between participants were also considered in the SSIP-RSA. The absolute difference between the ages of two participants was calculated to compute the participant-by-participant age distance matrix. In the participant-by-participant gender distance matrix, same gender is coded as one and different genders as 0. We then calculated the partial Spearman’s rank correlation (ρ; controlling for age and gender) between the upper triangles of the brain and wellbeing distance matrices (Fig. 1a, 3), and wrote the ρ value back to the voxel in the centre of the searchlight. A significantly positive ρ value means that participants with more similar brain activity patterns also had more similar wellbeing score profiles. Therefore, the ρ value can be a proxy to show the extent a voxel is related to individual differences in wellbeing at a specific time point. The code used to conduct the SSIP-RSA and all subsequent analyses is available for inspection on GitHub (https://github.com/YumengMa01/WellbeingNetworks).

### ICA

To identify networks and reduce dimensionality, we performed independent component (IC) analysis on the SSIP-RSA results using ‘melodic’ from the FSL software package (version 5.0; https://fsl.fmrib.ox.ac.uk/fsl/fslwiki/MELODIC)^157^. By enforcing spatial independence between components, the whole-brian ρ timeseries (Fig. 1b) was decomposed into ICs with a spatial brain map and its corresponding timecourse (Fig. 1c). The spatial map contains loadings of each voxel for the IC and the time course indicates contributions of that IC at each time point. The original whole-brain ρ timeseries is a linear combination of all IC maps and their associated time courses. We arbitrarily chose 100 components (i.e., dimensions) because the automatic estimation method in melodic did not converge. This number reflects a dimensionality higher than used in most resting state studies and is more typical of naturalistic fMRI, perhaps reflecting the increased richness of such data^81,158,159^. All 100 IC spatial maps can be inspected in the Supplementary Information (Fig. S2) and are available for download from NeuroVault^160^ (https://neurovault.org/collections/15197/).

To address the potential limitation that results might be contingent on the number of chosen dimensions, we also conducted the IC analysis with 50 and 150 dimensions (see Supplementary Figs. S1 and S3 for spatial maps). Specifically, we collapsed the thresholded IC maps for the 50-, 100-, and 150-dimensional IC analyses separately, using the nonzero mean and a minimum cluster size of 50 voxels. We then compared the resulting maps using demeaned spatial correlation (with ‘3ddot’). The mean 100-dimensional map correlated with the 50- and 150-dimensional maps with correlation coefficients of 0.75 and 0.87, respectively. This suggests that the spatial information in our results is preserved across different dimensionality choices.

### Hierarchical Clustering

To identify co-active spatially independent networks and further reduce dimensionality, we used hierarchical clustering with the Ward method to cluster ICs with similar time courses together (using the SciPy clustering package with default parameters; Fig. 1d)^161^. We made the relatively arbitrary decision to cut the dendrogram at the level of eight clades as this visually appears to parcellate the dendrogram by height into more differentiated clades (see Fig. 1d and 3a; also see Supplementary Figs. S4-S6 for dendrograms generated with 50, 100, and 150 dimensions). Furthermore, cutting higher up would have merged a large number of ICs in some instances, e.g., clades one and two would be merged, involving 51 networks (see Fig. 3a). This might have been difficult to interpret as it would involve much of the brain. Cutting lower down would require potentially interpreting a much larger number of clades, reducing explainability in any one theoretical framework like the self-related framework we propose. Thus, cutting at eight allowed a middle ground that permitted interpretability while permitting us to discover alternative hypotheses. All clades are available for inspection in NeuroVault^160^ (https://neurovault.org/collections/15197/).

A possible limitation of our methodological choices pertaining to the hierarchical clustering is that significantly different clustering might result from different IC analyses. To demonstrate the stability of the clusters presented, we correlated the 49 significant thresholded IC maps from the 100-dimensional IC analysis (see ‘Mantel Test’, next) with each IC map in the 50- and 150-dimensional IC analyses (using ‘3ddot’ again). The highest correlation was used as a proxy for best ‘matching’ components across analyses. These matches are identified in red for the spatial maps in Supplementary Figs. S1 and S3, and for the dendrograms in Supplementary Figs. S4 and S6. These results together suggest that there are similar components across IC analyses and that these cluster in the dendrograms in similar manners across IC analyses.

### Mantel Test

To test if brain regions shown in each of the ICs are statistically significantly associated with individual differences in wellbeing (i.e., the ρ values are significantly high), we conducted a Mantel test^162^. This is a permutation test used to measure the significance of the correlation between two distance matrices. With the wellbeing, age, and gender distance matrices remaining unchanged, rows and columns of the brain distance matrix were shuffled in the same random order. The partial Spearman’s rank correlation (ρ) between the shuffled brain and wellbeing distance matrices was recalculated. After repeating this process 10,000 times, a null distribution of the ρ values was generated and a p-value was calculated using a one-tailed test.

Instead of a searchlight analysis, the brain distance matrix was computed using the BOLD signal pattern extracted from voxels with significant contributions to each IC at the peak time point in the corresponding IC time course. In order to select voxels significantly contributing to each IC, an appropriate threshold needed to be chosen. The output IC brain maps from FSL ‘melodic’ contain z-scores. These are calculated by subtracting the mean of loadings in all voxels from each voxel’s loading, and then dividing the result by the standard deviation of loadings in all voxels, so that the brain maps have zero mean and unit variance. However, the mean and standard deviation from whole-brain voxels are not the same as those for an underlying null distribution, so the z-scores have no statistical meaning. Therefore, traditional p-value-based thresholding cannot be used. Because different z-score thresholds can generate brain regions with different cluster sizes and we do not know which captures individual differences in wellbeing the best, here we used multiple thresholds following the method used by Poppe et al.^163^, and did Mantel tests on the threshold that generated the highest SSIP-RSA ρ value for activity at the peak time point in the IC’s time course under evaluation.

Specifically, after dividing the loading of each voxel by the maximum loading value in the map, we created brain masks including voxels surviving nine different thresholds separately, based on the normalised z-scores (normalised z larger than 0.1, 0.2, 0.3, 0.4, 0.5, 0.6, 0.7, 0.8, and 0.9) and cluster sizes larger than 50 voxels to remove ICs containing small clusters of voxels. This resulted in nine masks for each IC, each containing a different number of voxels. We then re-did the SSIP-RSA on brain patterns inside each of the nine masks at the peak time point and selected the threshold that generated the highest ρ value. We then did the Mantel test, generating a p-value for each IC, correcting for the number of ICs tested using a false discovery rate with a threshold of p < 0.01. All Mantel test results are available for inspection in NeuroVault^160^ (https://neurovault.org/collections/15197/).

Note that we did hierarchical clustering on all ICs from the ICA. However, we could have done this step after the Mantel tests were performed. To determine if this affects the results, we examined the clades after hierarchical clustering on just the IC time courses associated with the Mantel results. In terms of interpretation (see Decoding), the results were not substantially different. The primary difference is that clades four and five merged (see Network Sets 3/4/5). If anything, this bolsters the anatomical assertions of the language-based narrative framework, suggesting a strong relationship between language-related and precuneus regions made throughout.

### Artefact Identification

Following the ICA, hierarchical clustering, and the Mantel test, we manually examined all results to identify artefacts. ICs 1-4, six, and 83 had no activity after limiting to positive values and applying a minimum cluster size threshold of 50 voxels. ICs 1-4, and six were all in a single clade and that clade was therefore dropped from further analyses. ICA readily picks up artefacts and we hand labelled all components that might be artifactual^164^. This identified nine likely artifactual components. These were IC 27 (with activity appearing as confetti throughout the brain, activity crossing multiple tissue boundaries and types, and activity in the white matter), 33 (crossing multiple tissue boundaries, white matter), 45 (crossing multiple tissue boundaries, peaks not located in brain tissue, white matter), 48 (peaks not located in brain tissue), 73 (peaks not located in brain tissue), 82 (peaks not located in brain tissue), 84 (confetti, crossing multiple tissue boundaries, peaks not located in brain tissue, white matter), 94 (peaks not located in brain tissue, white matter), and 95 (peaks not located in brain tissue). All nine of these were excluded from further analyses.

We also examined all Mantel test results and found one outcome in which activity appeared as whole brain confetti (encompassing 9222 voxels), with activity crossing multiple tissue boundaries, and with activity in white matter. Reexamining the original IC suggests it might have been an artefact though it was not labelled as such initially because it was a borderline case. As such, IC 92 was also excluded from further analyses. Note that the inclusion of all hand-labelled artefacts would not significantly change the interpretation of the results as determined by reverse inference (see Decoding, next). All artefact ICs and Mantel test results are available for inspection in NeuroVault^160^ (https://neurovault.org/collections/15197/).

### Decoding

To help specify the functions of individual networks and collections of co-active networks associated with individual differences in wellbeing obtained in the prior steps, we performed reverse inference using both NiMARE (https://nimare.readthedocs.io/en/stable/index.html) and Neurosynth (https://neurosynth.org/) ‘decoding’^86,165^. Specifically, 10 a priori term-based meta-analyses relevant to our embodied and narrative self-related processing hypotheses were collected using the NiMARE Python package (version 0.0.12)^165^. These derive from the Neurosynth dataset (version 0.7, released July 2018)^86^, constructed from 507,891 voxel activation coordinates reported in 14,371 neuroimaging studies where terms appear at high frequency in the titles and abstracts (compared to studies where those terms do not appear). Next, the Pearson correlation was calculated between these 10 meta-analyses and the vectorised co-active networks or clade maps obtained from the hierarchical clustering analysis, using only ICs significant by the Mantel test not deemed artifactual. For the embodied self-related processing, term-based meta-analyses were ‘affective’, ‘autonomic’, ‘emotion’, ‘interoceptive’, and ‘self referential’. For narrative self-related processing, correlated term-based meta-analyses were ‘autobiographical’, ‘language’, ‘memory retrieval’, ‘speech perception’, and ‘speech production’. These are presented in polar plots where the axes are r-values (Figs. 3-5) and only correlations ≥ 0.10 are interpreted in the text. In addition to these a priori chosen sets, we also post hoc included the correlations with the ‘visual’ and ‘attention’ term-based meta-analyses in the Fig. 2 polar plots.

The choice of some of these terms is obvious while others need further justification. The term ‘self referential’ might be considered to be relevant to both embodied and narrative self-related processing. However, none of the 166 studies associated with the ‘self referential’ meta-analysis uses the term ‘narrative’. More generally, the term ‘narrative’ was not contained in the titles or abstracts of any papers with a high enough frequency to be included in the Neurosynth database. For this reason, we included ‘self referential’ as an embodied term. Next, ‘autobiographical’ and ‘memory retrieval’ might also be considered relevant to both the embodied and narrative self-related processing. However, we defined the embodied self as focused in the present moment whereas the narrative self supports thinking about the past and future (via mental time travel; see Introduction). Furthermore, we have proposed a neurobiological model relating networks associated with language and autobiographical memory to narrative processing^60^. For these reasons, we included ‘autobiographical’ and ‘memory retrieval’ as narrative self-related terms.

To lend support to our a priori choice of terms, we also conducted data-driven reverse inference, making use of all 1,334 term-based meta-analyses available on the web-based version of Neurosynth (which also uses dataset version 0.7)^86^ Specifically, we provide the top 10 functional associations for the peak voxel in each IC network for any term with a z-value greater than zero. We also used the web-based version of Neurosynth (https://neurosynth.org/decode/) and Neurovault (https://neurovault.org/images/add_for_neurosynth) to ‘decode’ the top 10 meta-analyses associated with each of the seven sets of co-active networks or clades. In both cases, reported terms are only those related to cognitive and behavioural function, excluding anatomical and clearly methodological terms. Terms like the ‘central sulcus’, ‘precentral gyrus/sulcus’ and ‘postcentral gyrus’ were considered anatomical as were their functional equivalents like ‘motor’, ’premotor’, and ’somatosensory’ as they are often used interchangeably in literature. Note that discrepancies between results from the a priori set of terms and these more data-driven analyses are accounted for by differences in using the NiMARE and the web-based Neurosynth packages.

### Regression

We conducted an exploratory analysis to determine if socio-affective contexts in the film participants viewed scaled with co-activated networks determined to be correlated with individual differences in wellbeing (Fig. 1d). This was accomplished by regressing the IC time courses against six film annotations from Masson and Isik^88^. These were valence, arousal, presence of social interactions (social), presence of a person speaking (speaking), pixel brightness, and auditory amplitude. They were generated for each three-second film segment after excluding the opening and ending credits. Specifically, valence and arousal annotations were derived by four raters based on a 1-5 Likert scale. As the ratings were moderately consistent between raters (valence Spearman ρ = 0.62, arousal ρ = 0.35), the mean ratings are used as objective features of the film. The social annotation was generated by two raters based on whether there was actions or communication happening between two or more characters (presence of social interaction = 1, absence = 0). The inter-rater consistency was high with a correlation value r = 0.86. The speaking annotation was rated by the same two raters (presence of person speaking = 1, absence = 0). Finally, pixel brightness and auditory amplitude were generated and averaged over each three-second segment and are considered ‘low-level’ controls^88^.

Before regression, the film annotation timeseries were normalised and upsampled from three to one second to match the temporal resolution of IC time courses. The IC time courses were shifted by four seconds to account for haemodynamic delay and rise to peak. They were then treated as dependent variables in the multiple linear regression model, while independent variables were the four annotations and two controls (i.e., pixel brightness and auditory amplitude). Considering the autocorrelations in both the IC time courses and the timeseries of film annotations, we used the non-parametric permutation to test significance. The normalised timeseries of the annotation to be tested was randomly shuffled, upsampled, and then put into the multiple linear regression model while the timeseries of other annotations remained unchanged. A null distribution of the β values for the annotation tested can then be obtained after shuffling its timeseries 10,000 times. A two-tailed permutation test was performed to obtain p-values. Because 0.01 was too conservative to yield significant results, we used an alpha value of 0.05 after correcting for the number of ICs and annotations tested using a false discovery rate algorithm.

## Data Availability

The preprocessed data used in analyses is publically available on OpenNeuro (https://openneuro.org/datasets/ds002837/versions/2.0.0). The code used to generate results is available on GitHub (https://github.com/YumengMa01/WellbeingNetworks). Results are available in Neurovault (https://neurovault.org/collections/15197/).

## Acknowledgements

We would like to thank the Birkbeck-UCL Centre for Neuroimaging (BUCNI) for supporting the NNDb. We also thank Annette Glotfelty and Daniel Lametti for helpful feedback on the manuscript and The LAB Lab and UNITy Project for discussion. JIS is supported by a Wellcome Leap award (‘Understanding Neuroplasticity Induced by TrYptamines’). JIS would also like to thank BB for my wellbeing and the Barone-Shaw family and the sanctuary of ‘Two Pines’.

## Author Contributions

YM and JIS conceived of the study and wrote the manuscript. JIS did the fMRI preprocessing. YM did all of the analyses and made Figs. 1-6, S1-S6 and Table S1. JIS made Fig. 7.

## Additional Information

