## Supplementary Information for "Individual differences in wellbeing are supported by separable sets of co-active self- and visual-attention-related brain networks"

**Figure S1. 50-dimensional spatial maps.** *Spatial maps generated from independent component analysis with 50 components. Maps are thresholded to only include positive values and cluster sizes larger than 50 voxels, and any maps with no surviving clusters after thresholding are excluded from display. A red border highlights ICs with spatial maps that are most correlated with those of the 49 significant ICs identified from the 100-dimensional analysis in the Mantel test. The corresponding IC number and correlation value are shown.*

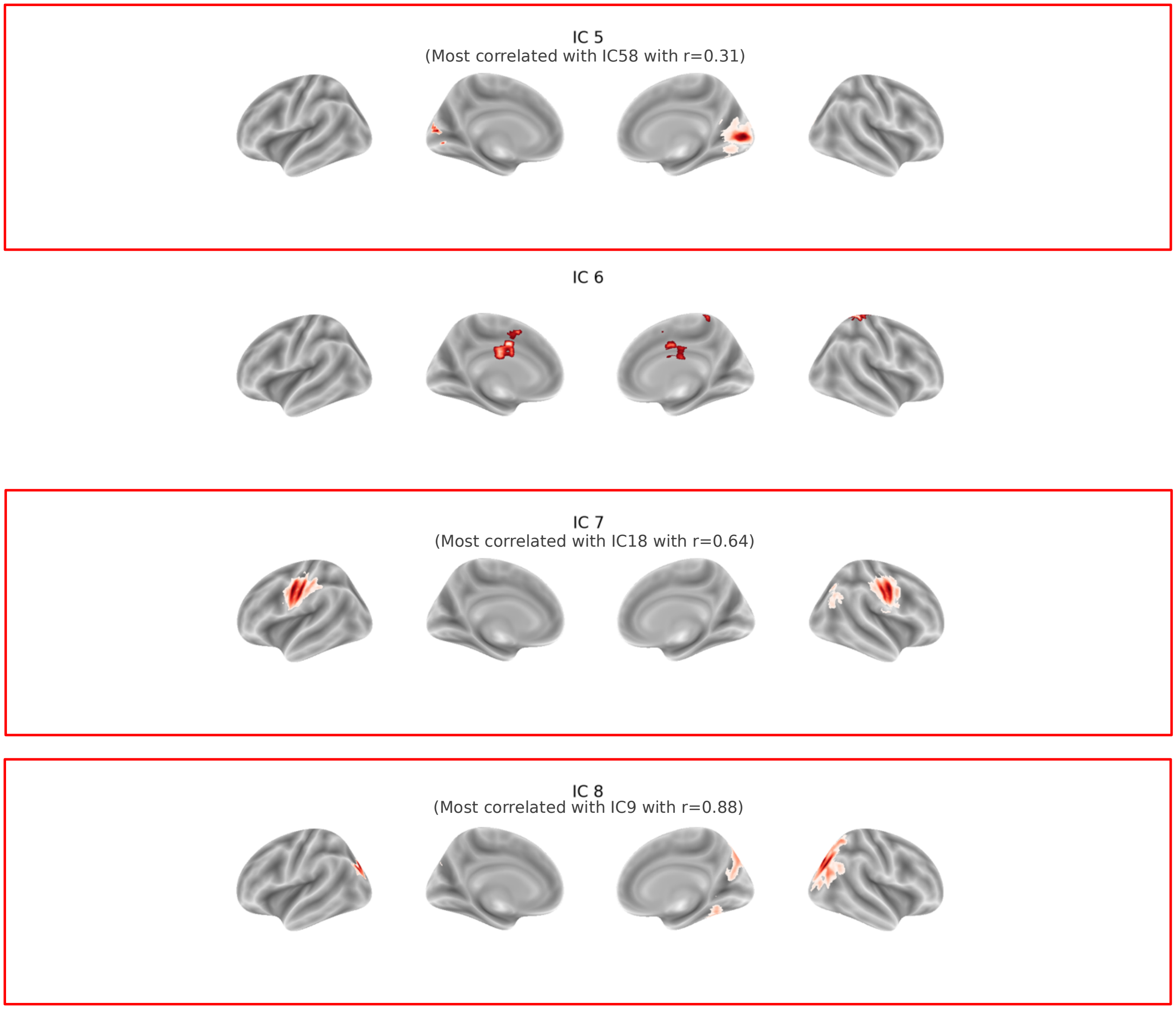

IC 9
(Most correlated with IC7 with r=0.93)

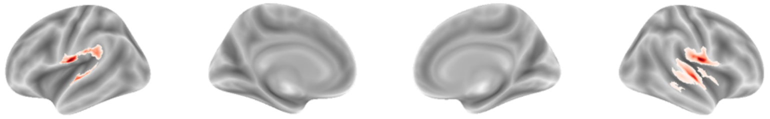

IC 10
(Most correlated with IC10 with r=0.92)

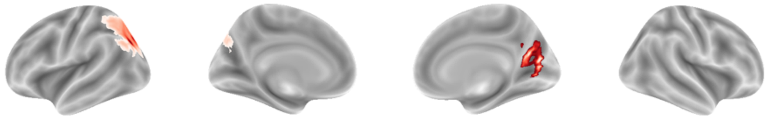

IC 11
(Most correlated with IC8 with r=0.94)

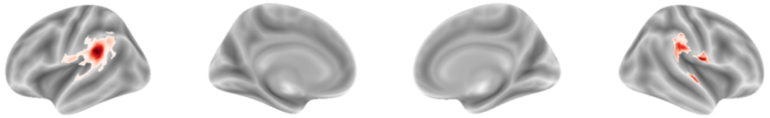

IC 12
(Most correlated with IC12 with r=0.83 and IC70 with r = 0.22)

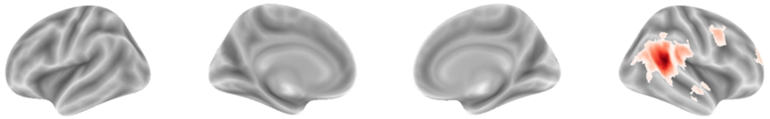

IC 13
(Most correlated with IC40 with r=0.39)

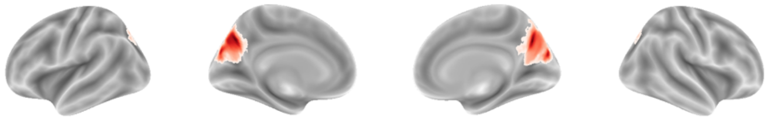

IC 14
(Most correlated with IC15 with r=0.87)

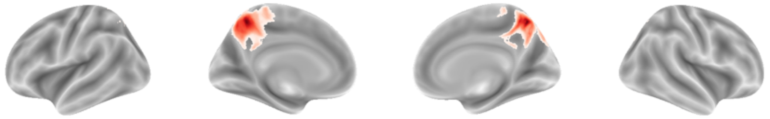

IC 15

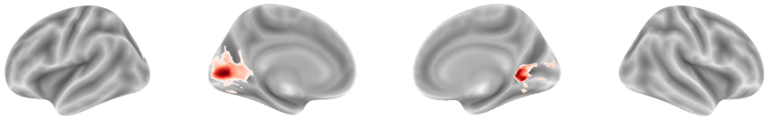

IC 16
(Most correlated with IC16 with r=0.89 and IC66 with r = 0.29)

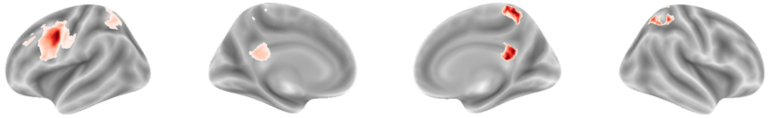

IC 17

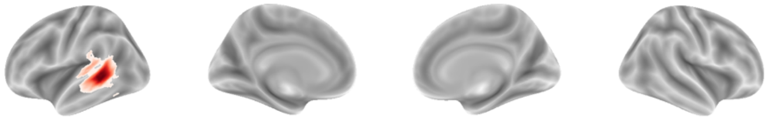

IC 18
(Most correlated with IC14 with r=0.80)

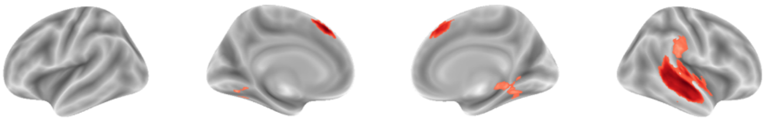

IC 19
(Most correlated with IC19 with r=0.88 and IC47 with r = 0.37)

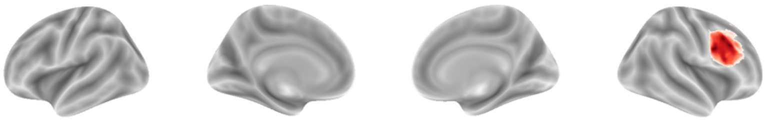

IC 20
(Most correlated with IC28 with r=0.80 and IC44 with r = 0.44)

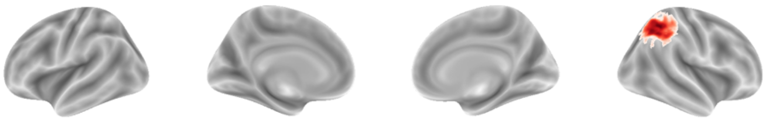

IC 21

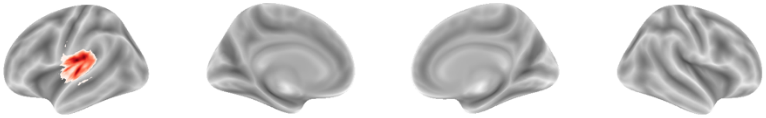

IC 22
(Most correlated with IC42 with r=0.73 and IC55 with r = 0.34)

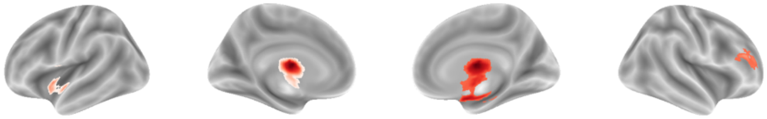

IC 23
(Most correlated with IC32 with r=0.79 and IC88 with r = 0.33)

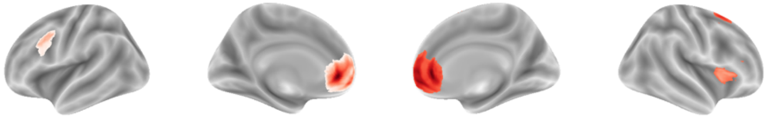

IC 24

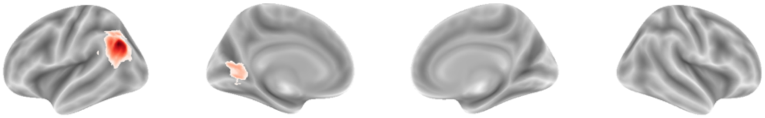

IC 25
(Most correlated with IC31 with r=0.57)

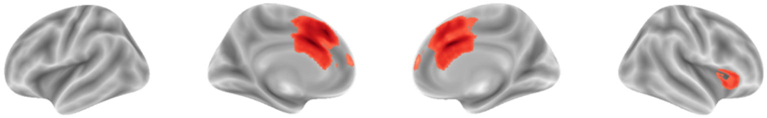

IC 26
(Most correlated with IC21 with r=0.66, IC90 with r = 0.21 and IC99 with r = 0.13)

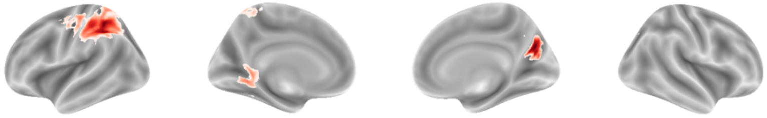

IC 27
(Most correlated with IC57 with r=0.25)

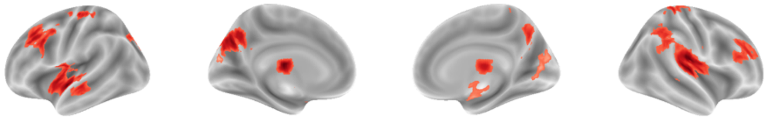

IC 28
(Most correlated with IC35 with r=0.60)

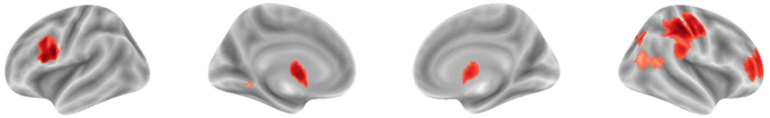

IC 29

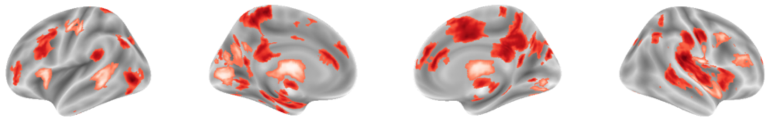

IC 30
(Most correlated with IC26 with r=0.76)

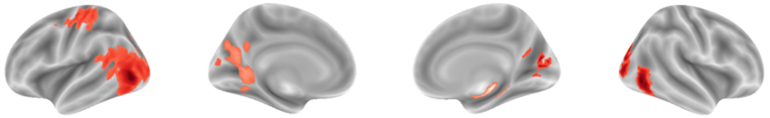

IC 31
(Most correlated with IC29 with r=0.56 and IC86 with r = 0.32)

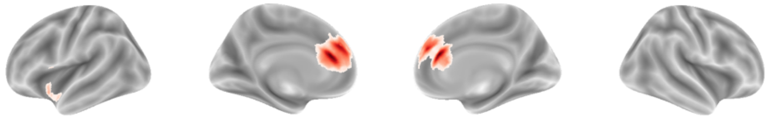

IC 32
(Most correlated with IC62 with r=0.17 and IC85 with r = 0.43)

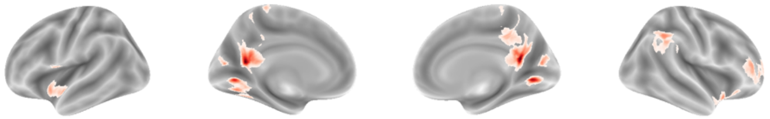

IC 33

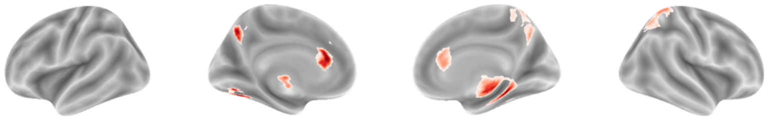

IC 34
(Most correlated with IC54 with r=0.65 and IC64 with r = 0.38)

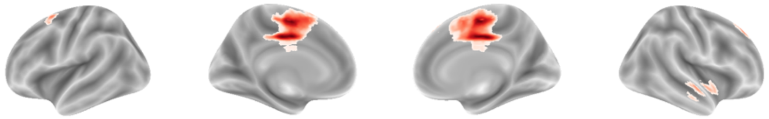

IC 35

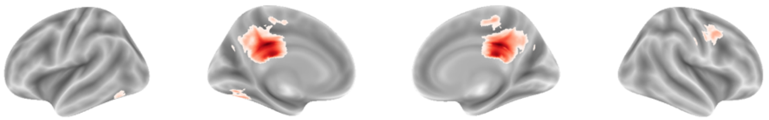

IC 36
(Most correlated with IC51 with r=0.69 and IC68 with r = 0.65)

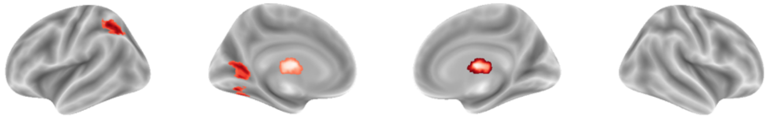

IC 37

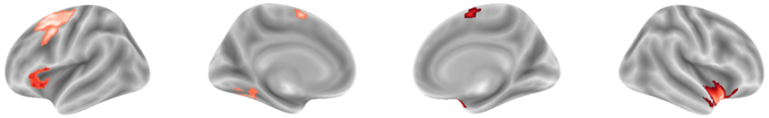

IC 38
(Most correlated with IC50 with r=0.73 and IC78 with r = 0.63)

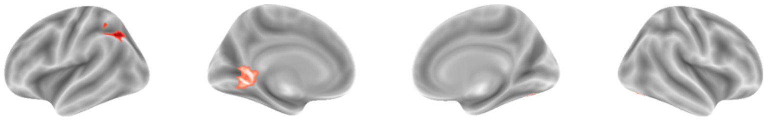

IC 39

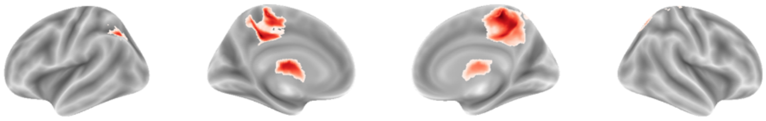

IC 40
(Most correlated with IC43 with r=0.79 and IC96 with r = 0.09)

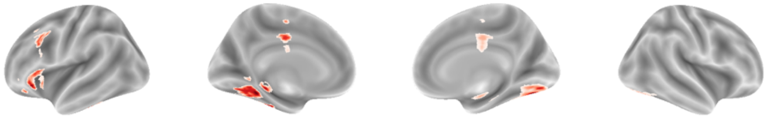

IC 41
(Most correlated with IC67 with r=0.17 and IC72 with r = 0.15)

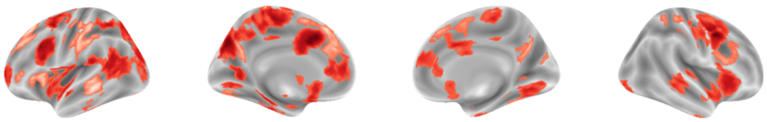

IC 42
(Most correlated with IC59 with r=0.48)

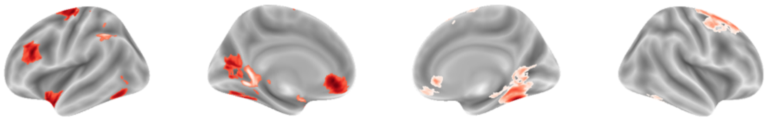

IC 43

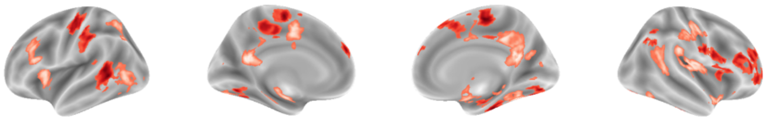

IC 44
(Most correlated with IC80 with r=0.24)

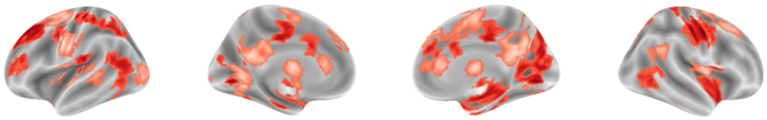

IC 45

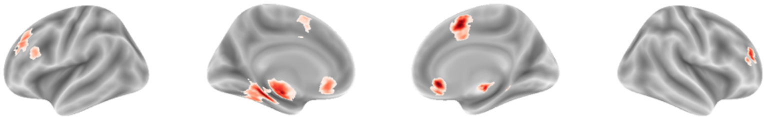

IC 46

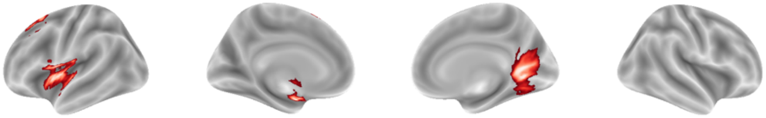

IC 47
(Most correlated with IC60 with r=0.70 and IC87 with r = 0.31)

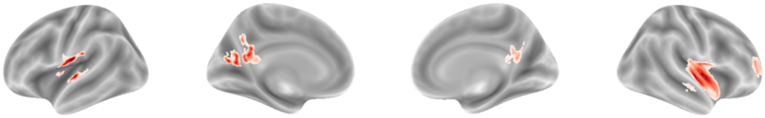

IC 48
(Most correlated with IC76 with r=0.36)

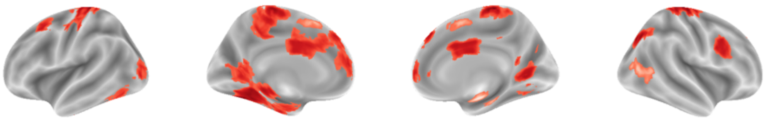

IC 49

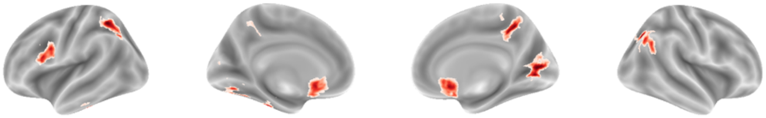

IC 50
(Most correlated with IC22 with r=0.16 and IC63 with r = 0.43)

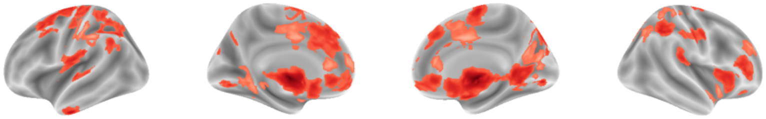

**Figure S2. 100-dimensional spatial maps.** *Spatial maps generated from independent component analysis with 100 components. Maps are thresholded to only include positive values and cluster sizes larger than 50 voxels, and any maps with no surviving clusters after thresholding are excluded from display.*

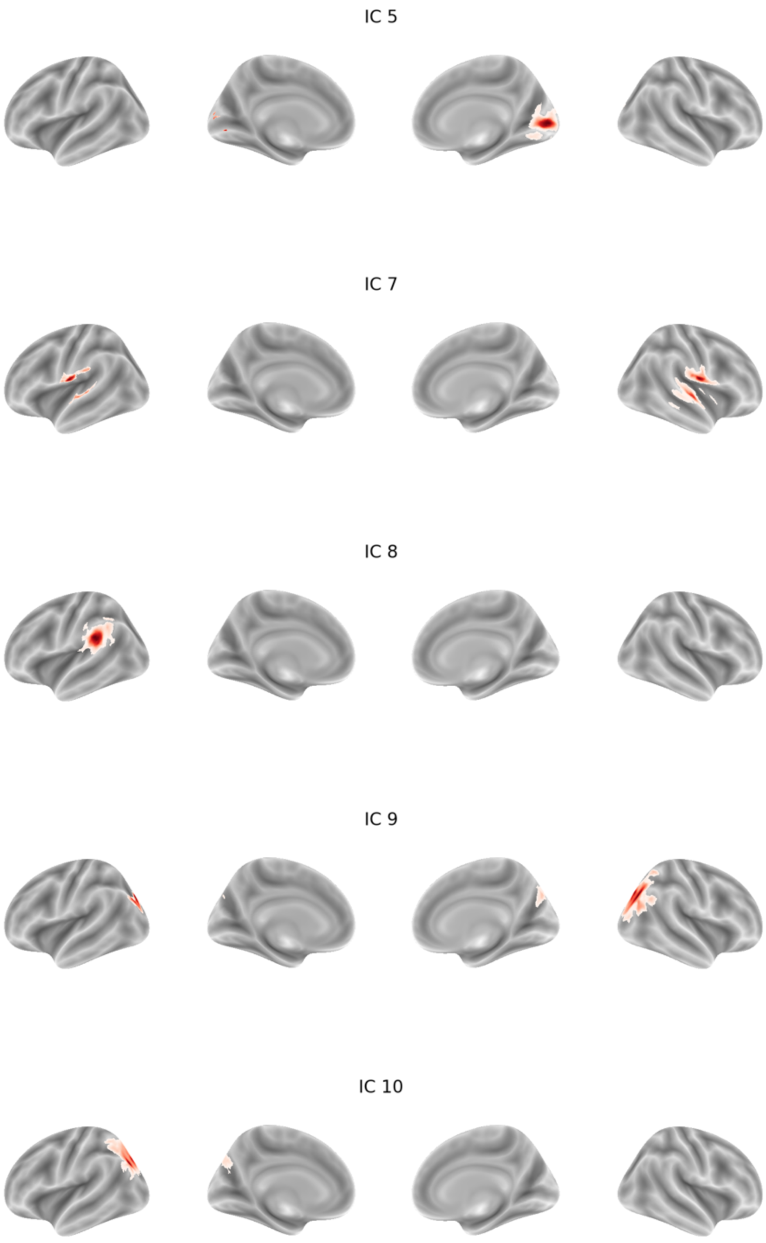

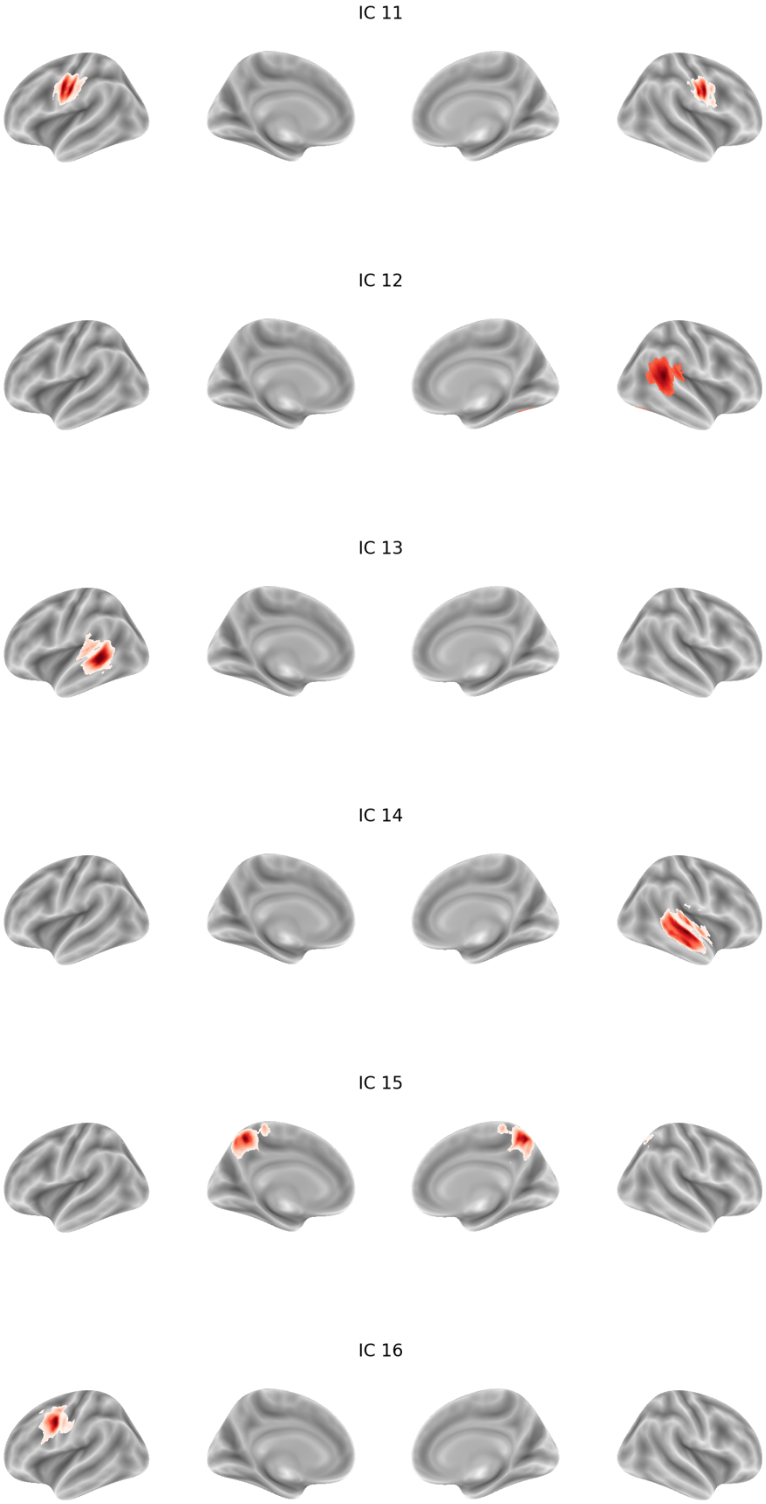
IC 11
IC 12
IC 13
IC 14
IC 15
IC 16

IC 17

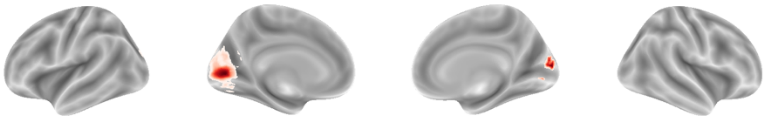

IC 18

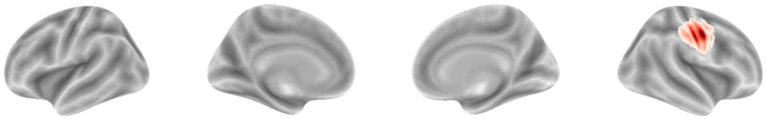

IC 19

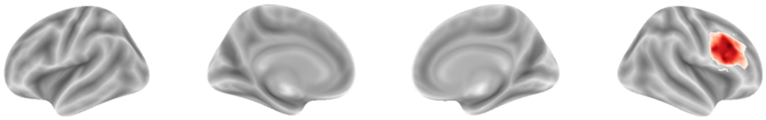

IC 20

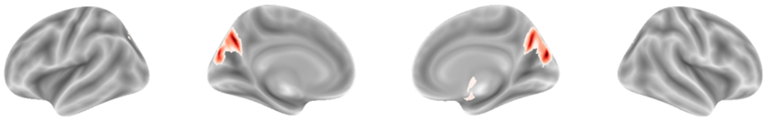

IC 21

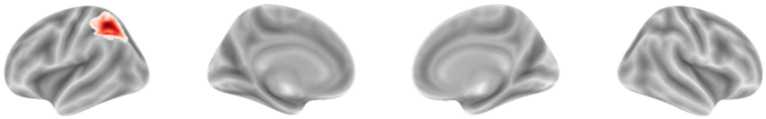

IC 22

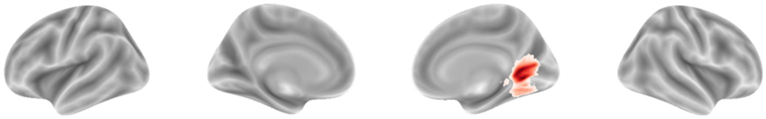

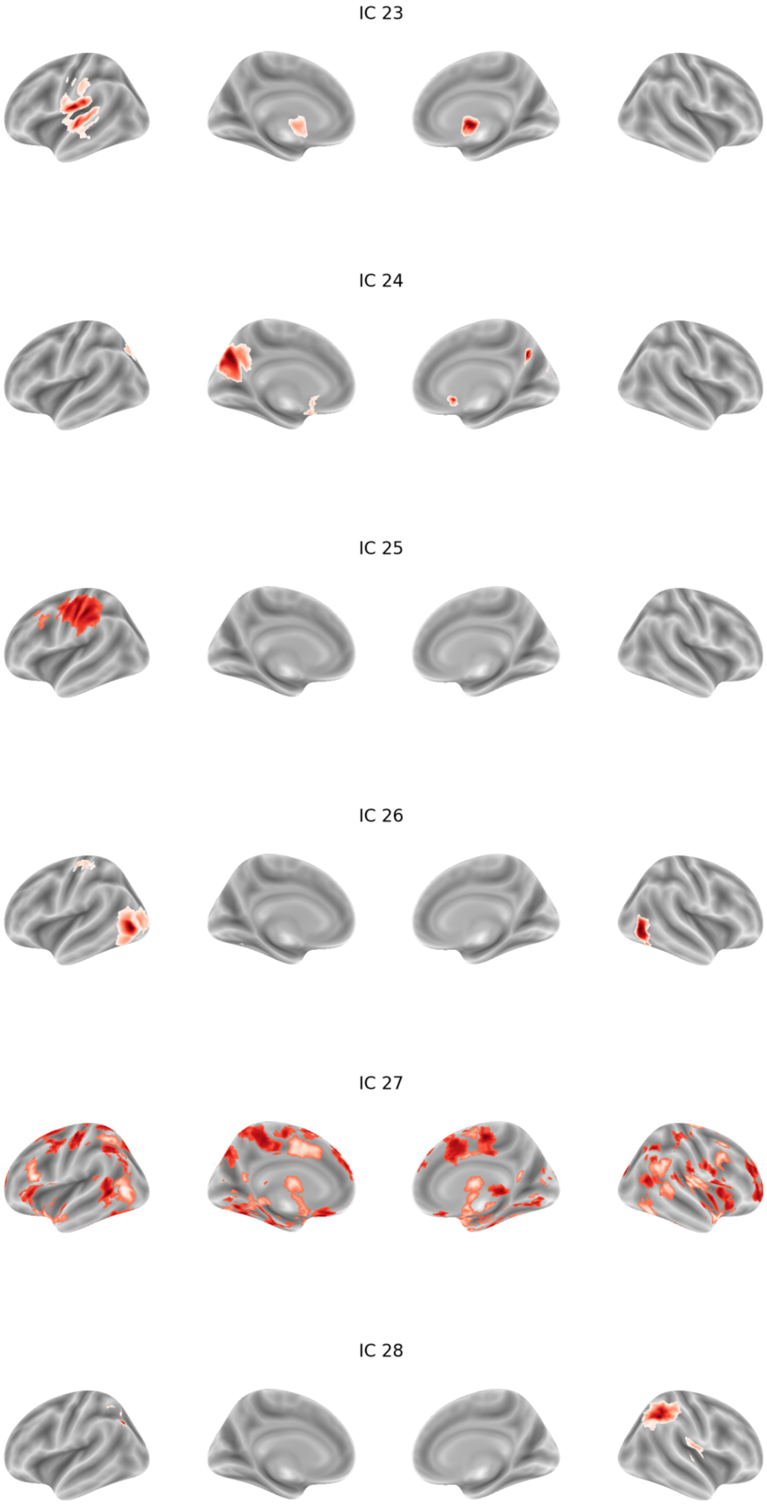
IC 23
IC 24
IC 25
IC 26
IC 27
IC 28

IC 29

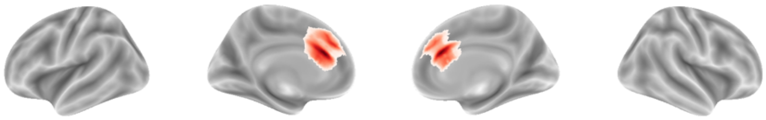

IC 30

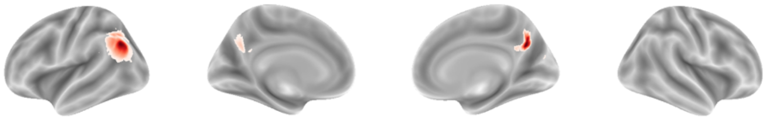

IC 31

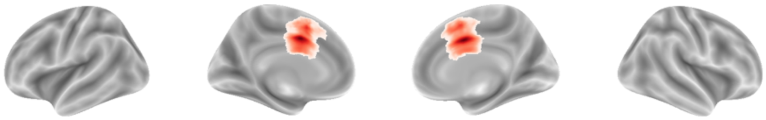

IC 32

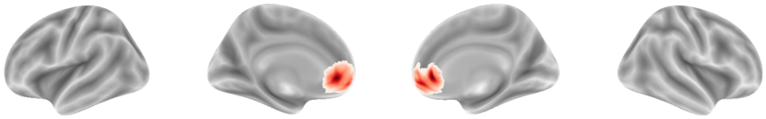

IC 33

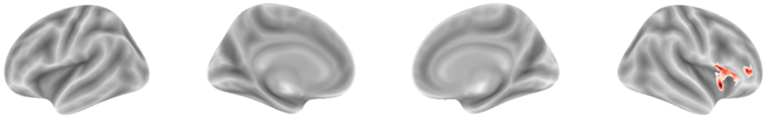

IC 34

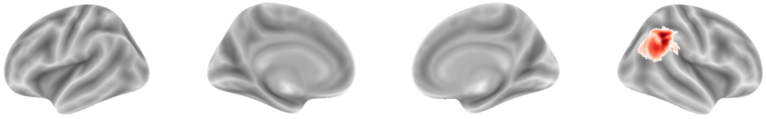

IC 35

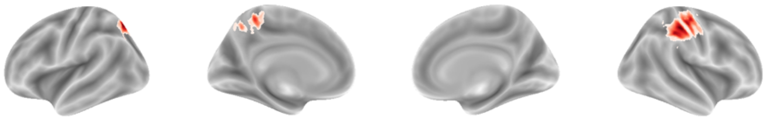

IC 36

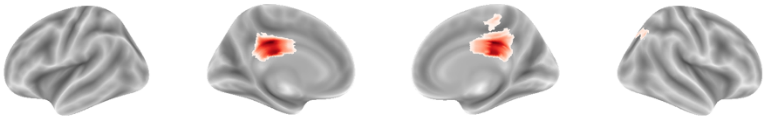

IC 37

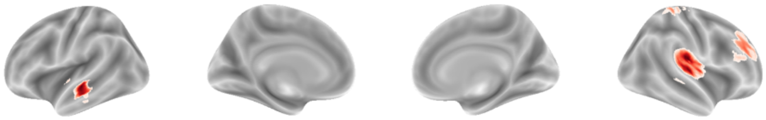

IC 38

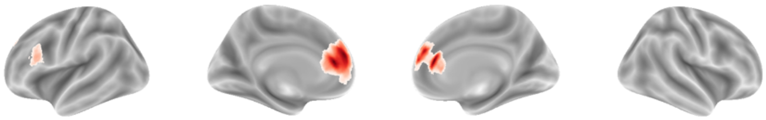

IC 39

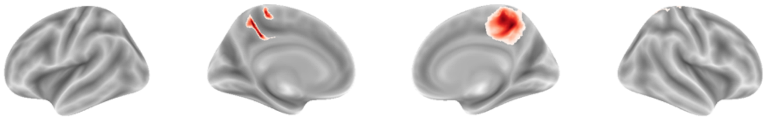

IC 40

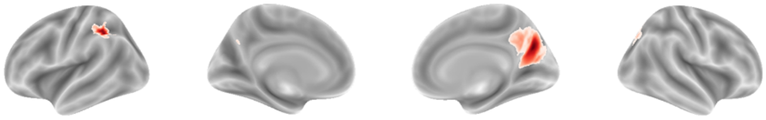

IC 41

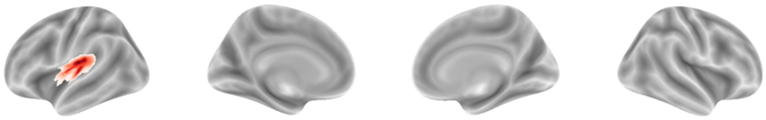

IC 42

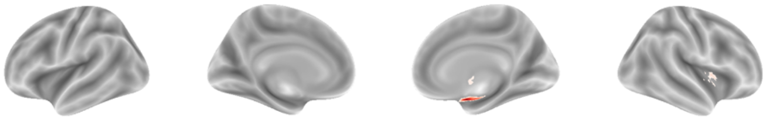

IC 43

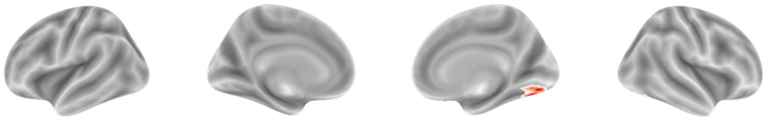

IC 44

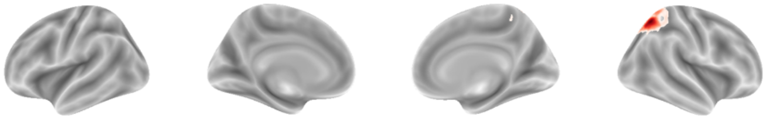

IC 45

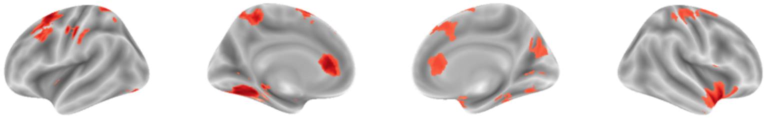

IC 46

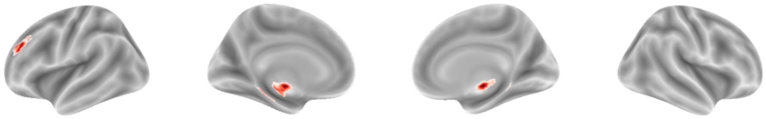

IC 47

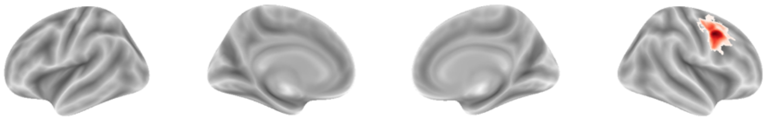

IC 48

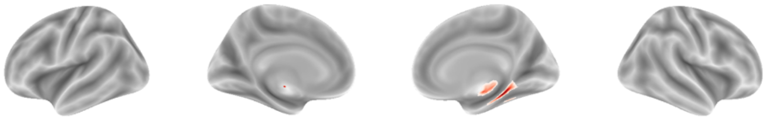

IC 49

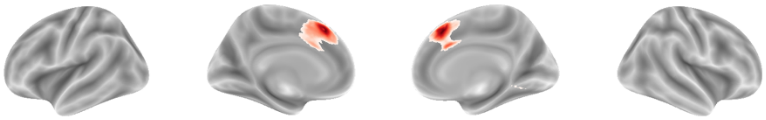

IC 50

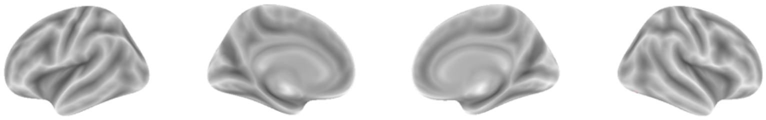

IC 51

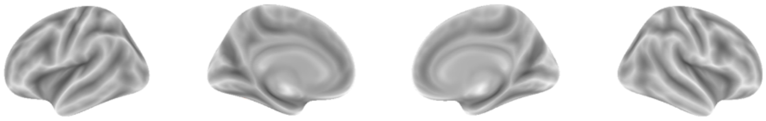

IC 52

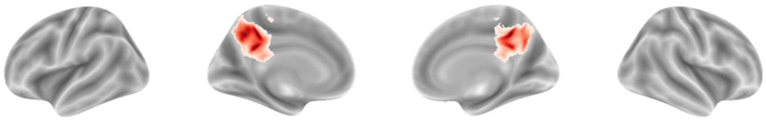

IC 53

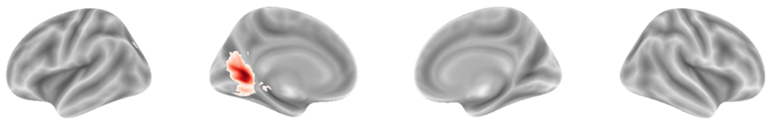

IC 54

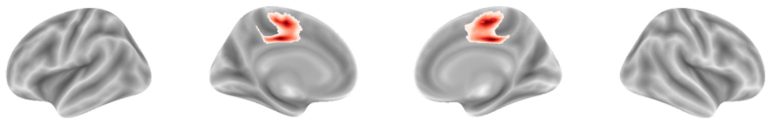

IC 55

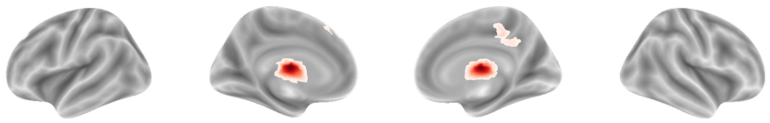

IC 56

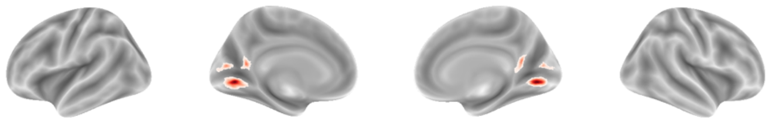

IC 57

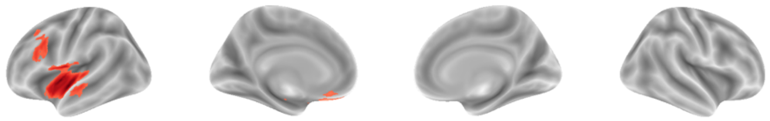

IC 58

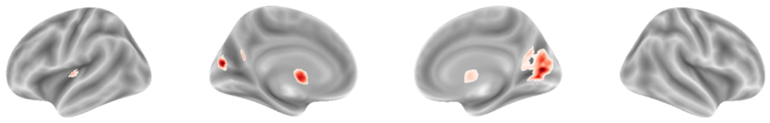

IC 59

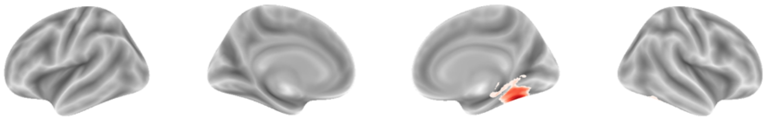

IC 60

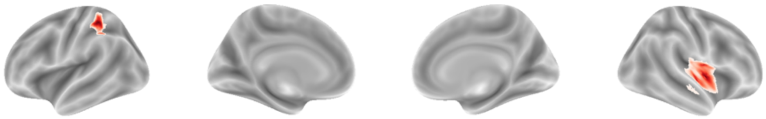

IC 61

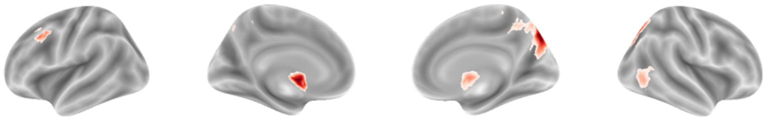

IC 62

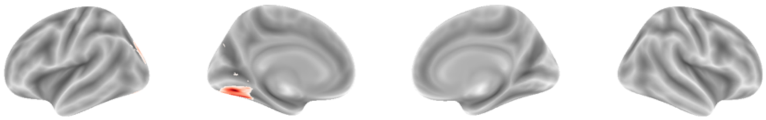

IC 63

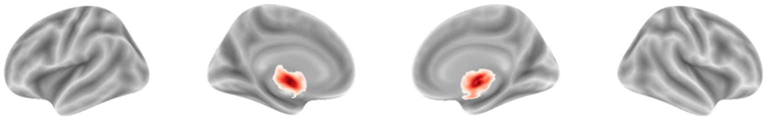

IC 64

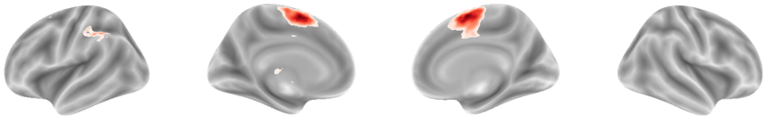

IC 65

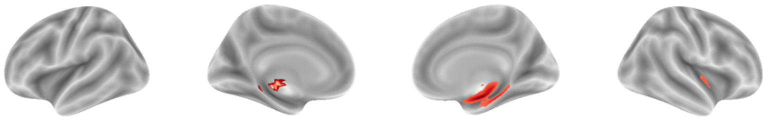

IC 66

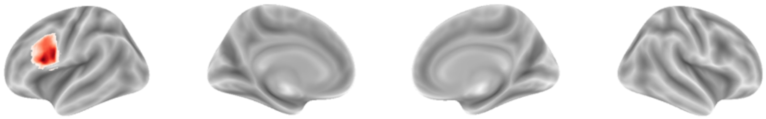

IC 67

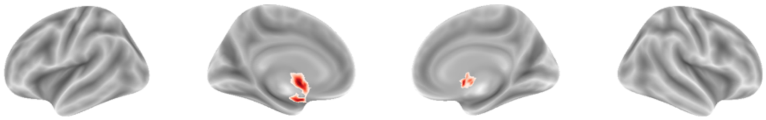

IC 68

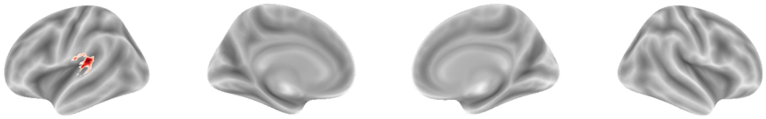

IC 69

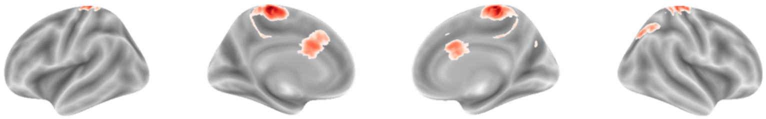

IC 70

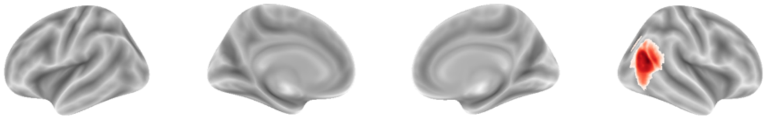

IC 71

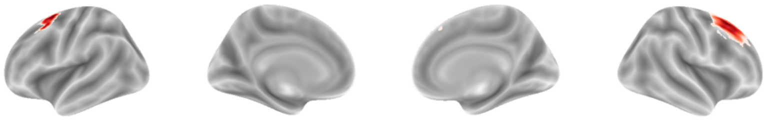

IC 72

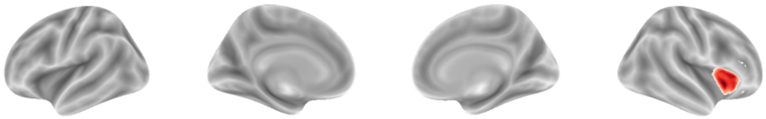

IC 73

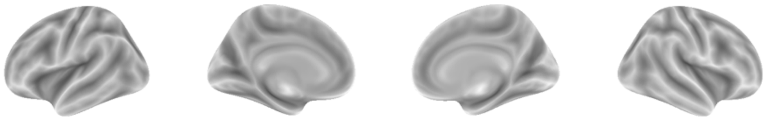

IC 74

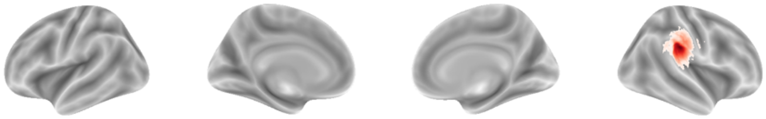

IC 75

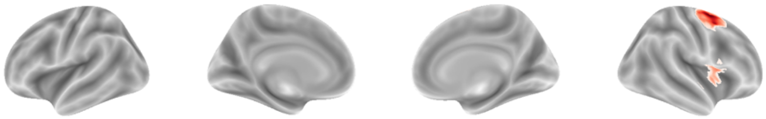

IC 76

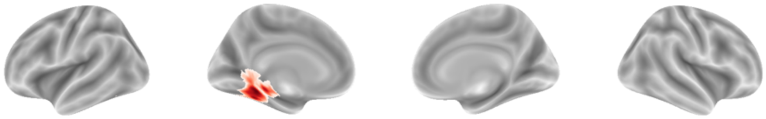

IC 77

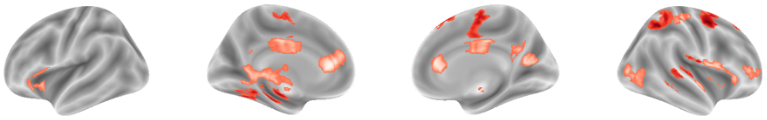

IC 78

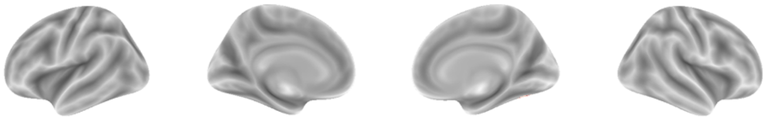

IC 79

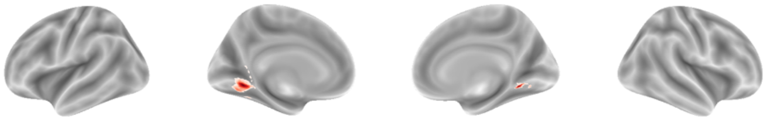

IC 80

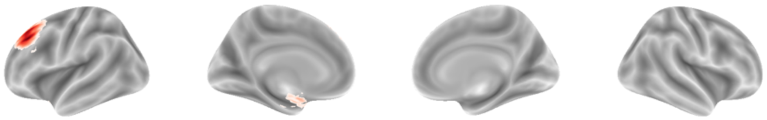

IC 81

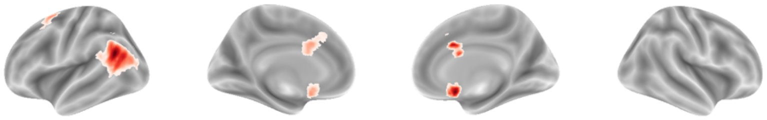

IC 82

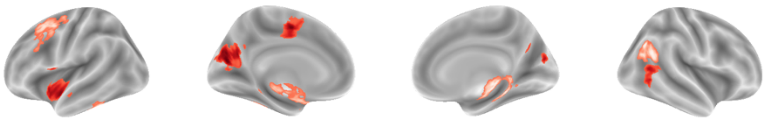

IC 83

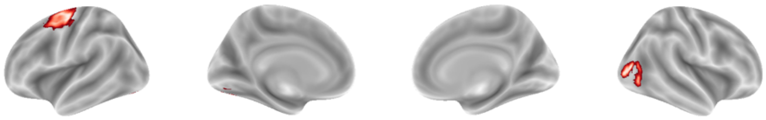

IC 84

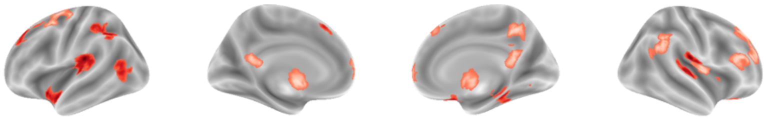

IC 85

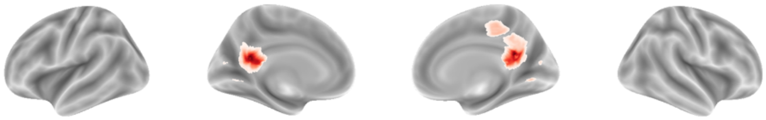

IC 86

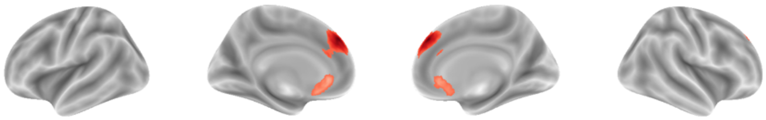

IC 87

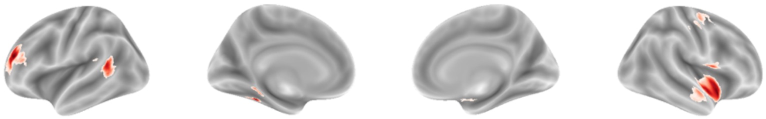

IC 88

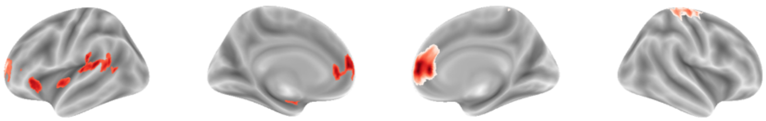

IC 89

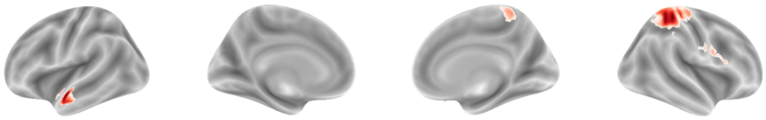

IC 90

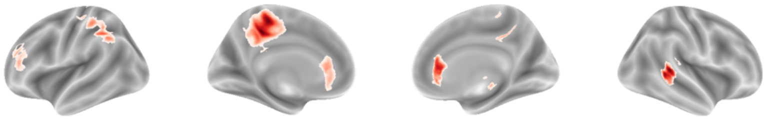

IC 91

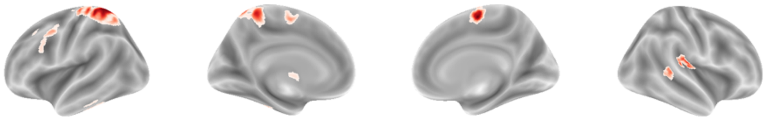

IC 92

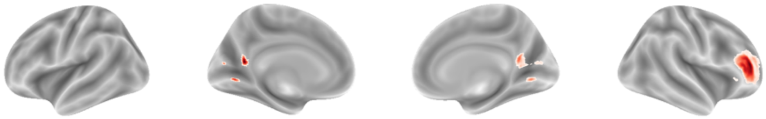

IC 93

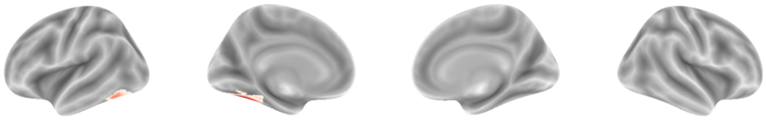

IC 94

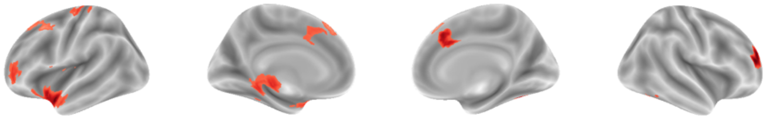

IC 95

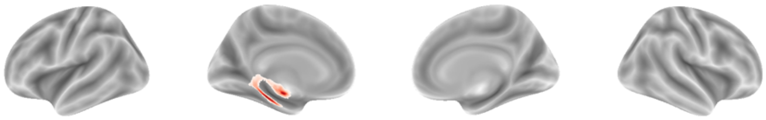

IC 96

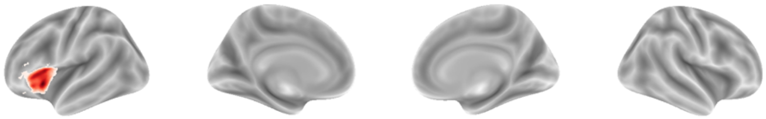

IC 97

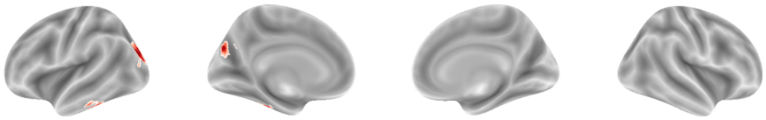

IC 98

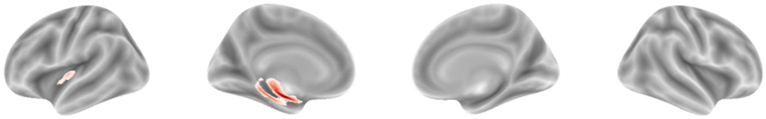

**Figure S3. 150-dimensional spatial maps.** *Spatial maps generated from independent component analysis with 150 components. Maps are thresholded to only include positive values and cluster sizes larger than 50 voxels, and any maps with no surviving clusters after thresholding are excluded from display. A red border highlights ICs with spatial maps that are most correlated with those of the 49 significant ICs identified from the 100-dimensional analysis in the Mantel test. The corresponding IC number and correlation value are shown.*

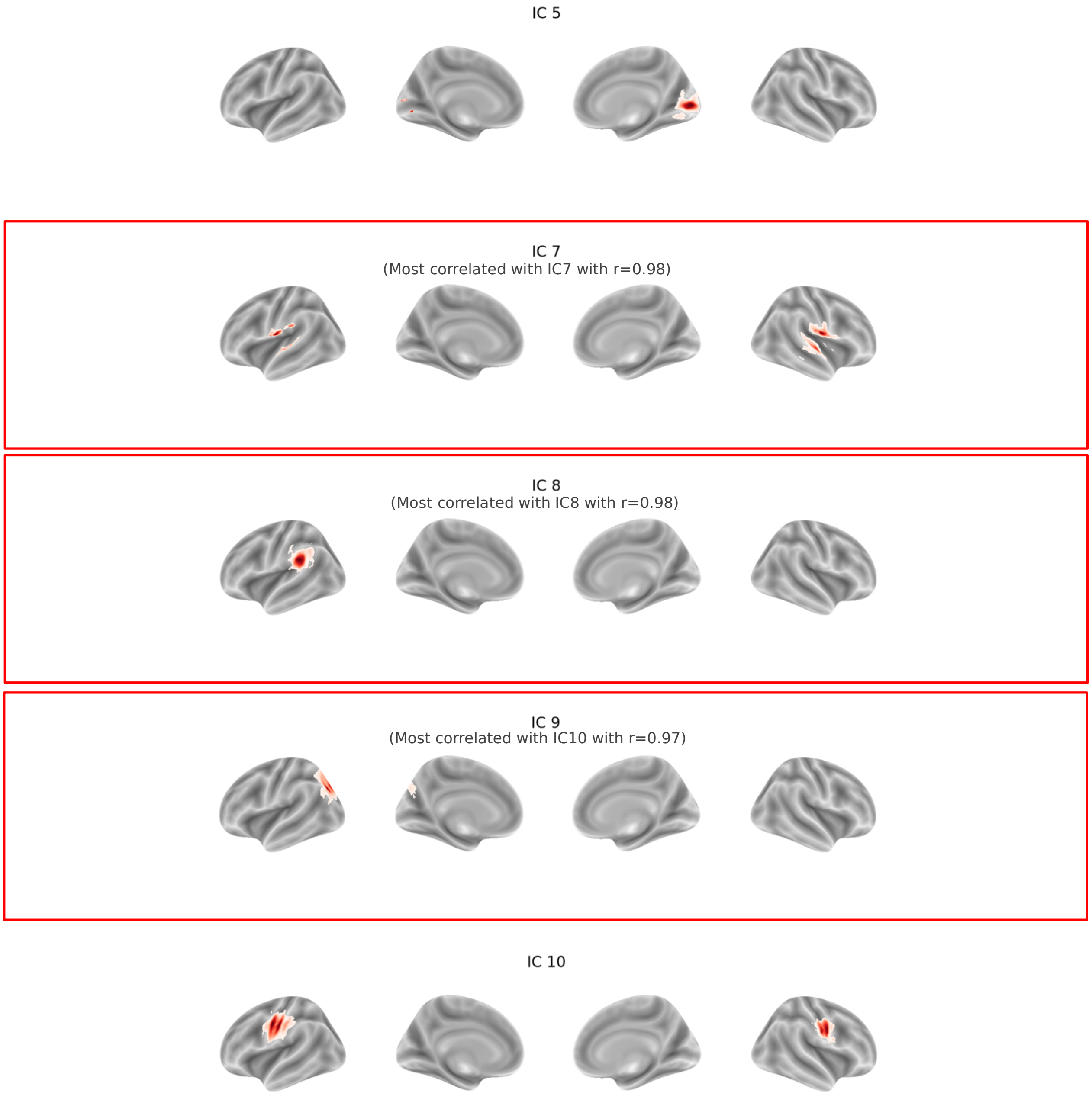

IC 11
(Most correlated with IC12 with r=0.95)

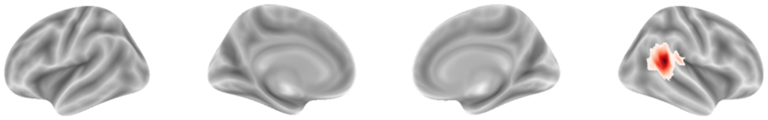

IC 12
(Most correlated with IC18 with r=0.96)

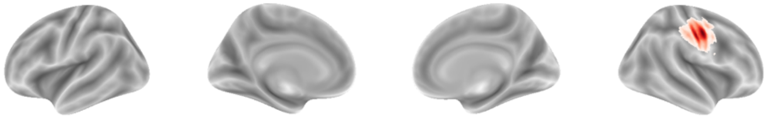

IC 13
(Most correlated with IC16 with r=0.97)

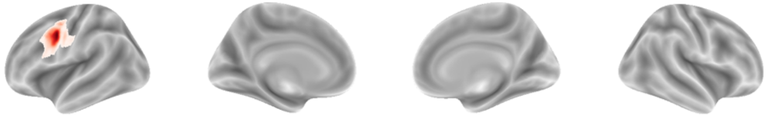

IC 14

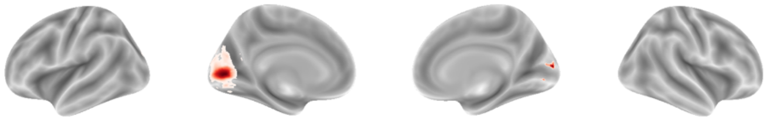

IC 15
(Most correlated with IC15 with r=0.95)

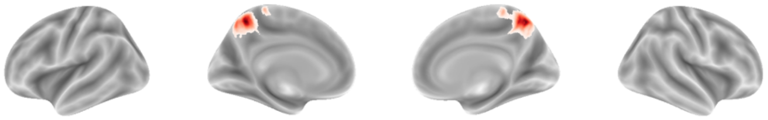

IC 16

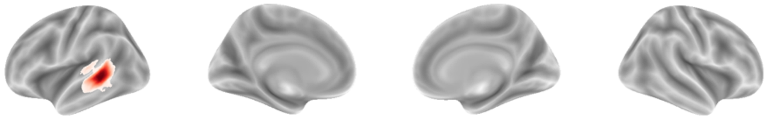

IC 17
(Most correlated with IC9 with r=0.79)

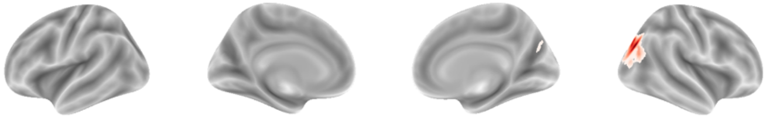

IC 18

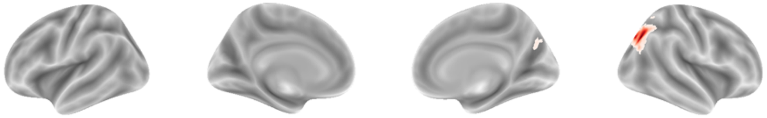

IC 19

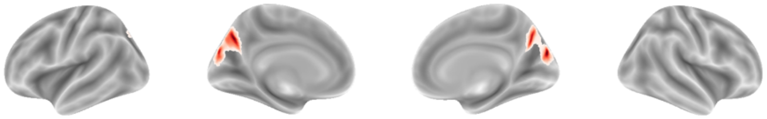

IC 21
(Most correlated with IC21 with r=0.87)

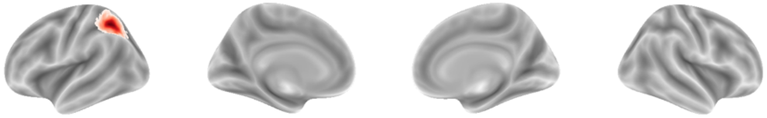

IC 22

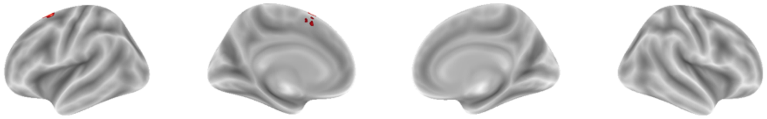

IC 23
(Most correlated with IC19 with r=0.90)

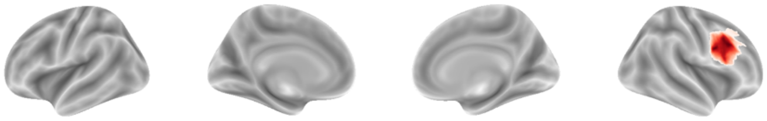

IC 24
(Most correlated with IC22 with r=0.94)

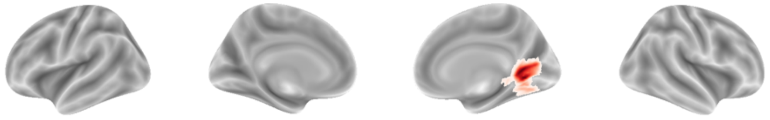

IC 25

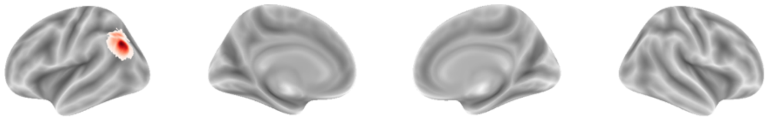

IC 26
(Most correlated with IC32 with r=0.92)

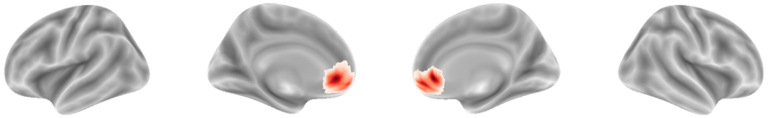

IC 27
(Most correlated with IC44 with r=0.87)

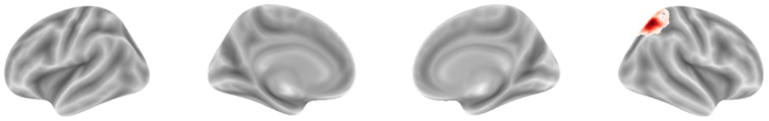

IC 28
(Most correlated with IC26 with r=0.85)

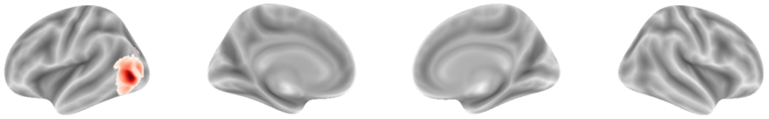

IC 29

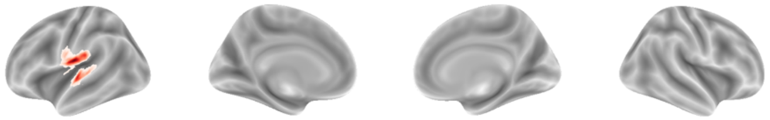

IC 30
(Most correlated with IC28 with r=0.88)

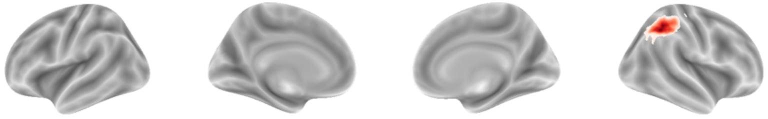

IC 31

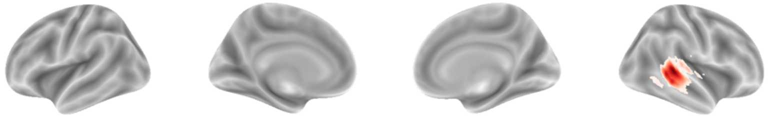

IC 32
(Most correlated with IC14 with r=0.70)

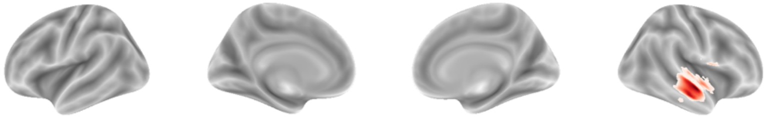

IC 33

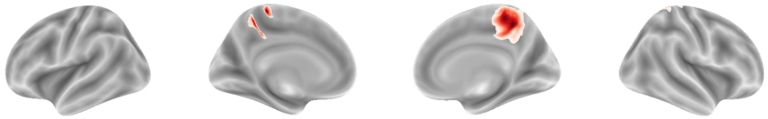

IC 34
(Most correlated with IC50 with r=0.96)

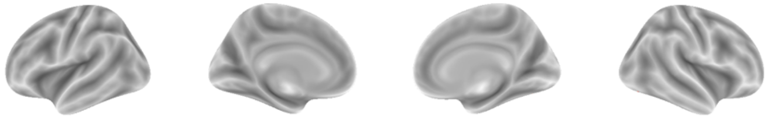

IC 35
(Most correlated with IC35 with r=0.86)

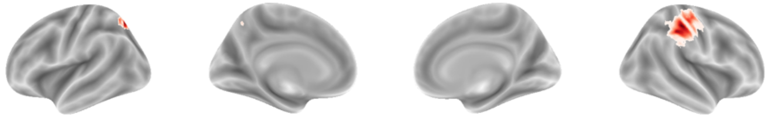

IC 36
(Most correlated with IC55 with r=0.91)

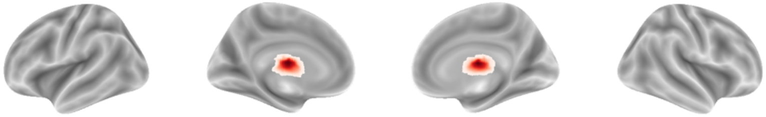

IC 37

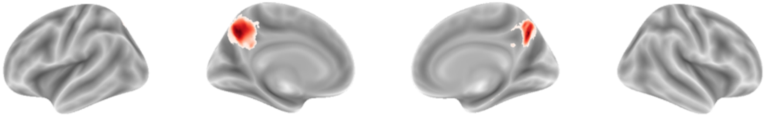

IC 38

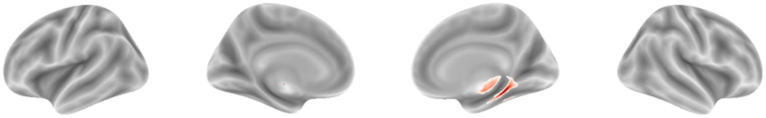

IC 39

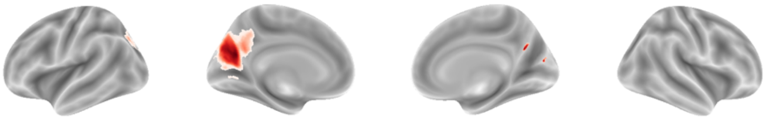

IC 40

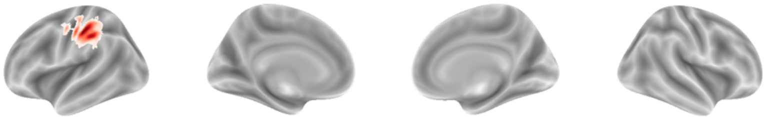

IC 41
(Most correlated with IC58 with r=0.86)

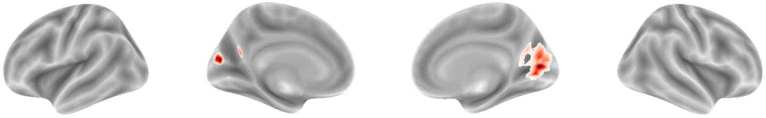

IC 42
(Most correlated with IC59 with r=0.92)

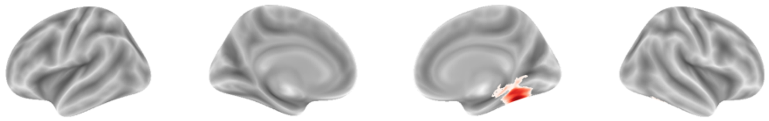

IC 43
(Most correlated with IC43 with r=0.92)

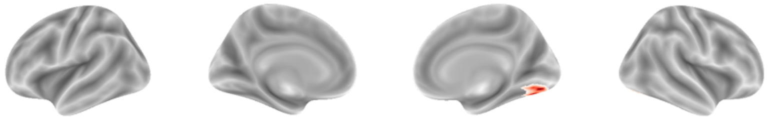

IC 44

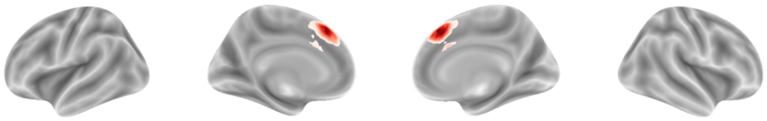

IC 45

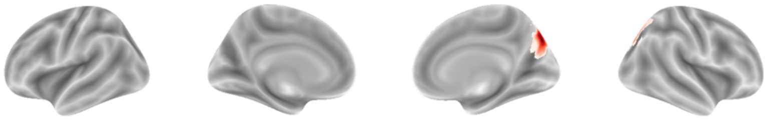

IC 46

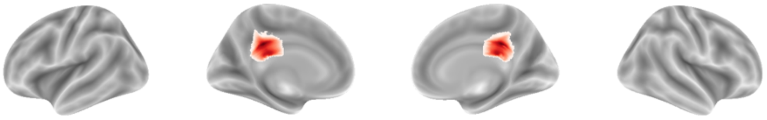

IC 47
(Most correlated with IC47 with r=0.92)

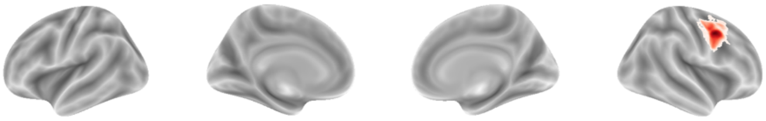

IC 48

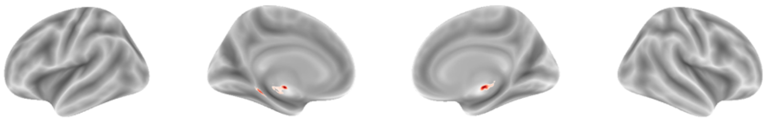

IC 49
(Most correlated with IC31 with r=0.88)

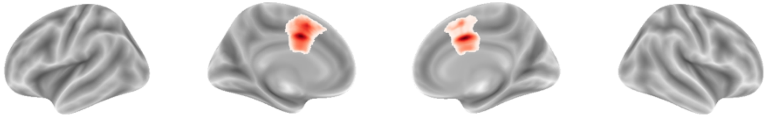

IC 50
(Most correlated with IC29 with r=0.84)

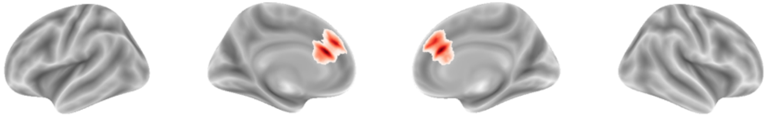

IC 51

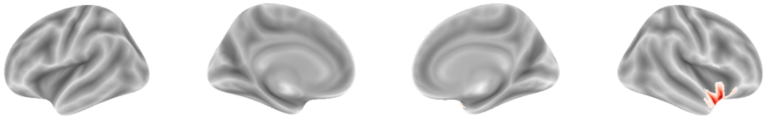

IC 52

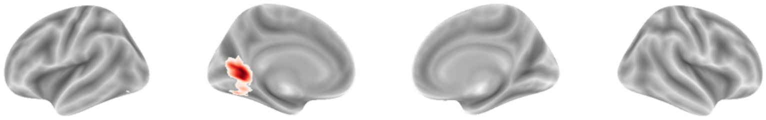

IC 53
(Most correlated with IC51 with r=0.84)

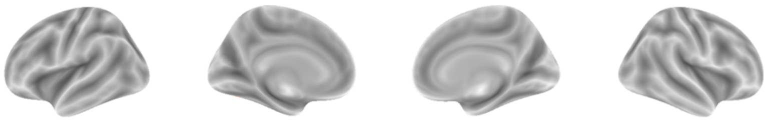

IC 54
(Most correlated with IC62 with r=0.77)

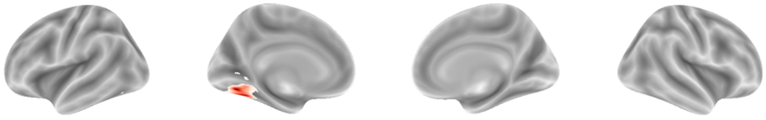

IC 55
(Most correlated with IC54 with r=0.91)

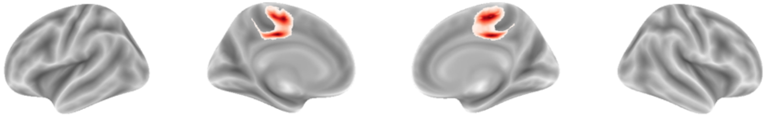

IC 56

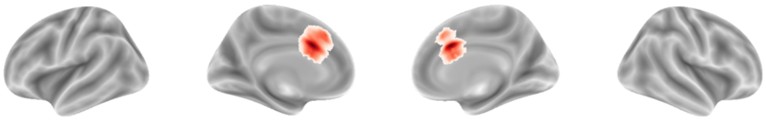

IC 57

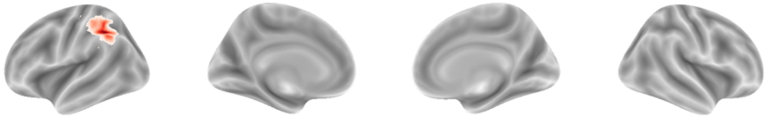

IC 58
(Most correlated with IC78 with r=0.97)

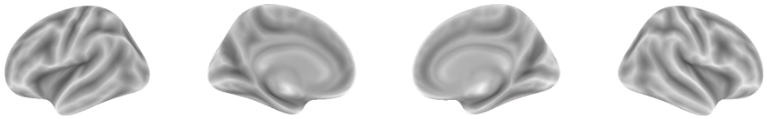

IC 59
(Most correlated with IC70 with r=0.59)

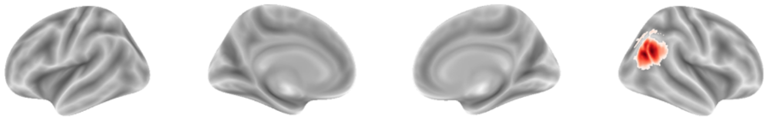

IC 60
(Most correlated with IC42 with r=0.89)

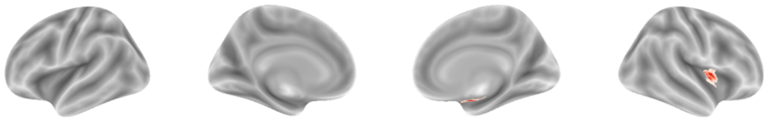

IC 61

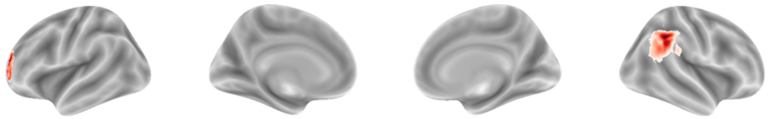

IC 62

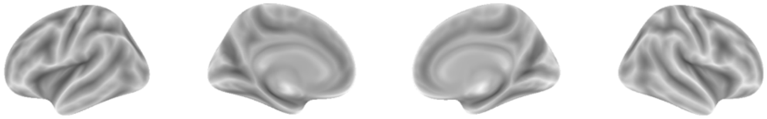

IC 63
(Most correlated with IC66 with r=0.88)

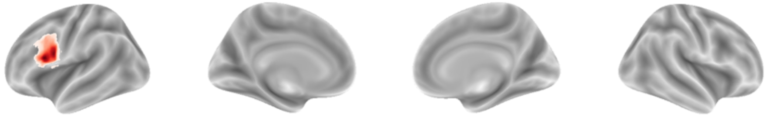

IC 64

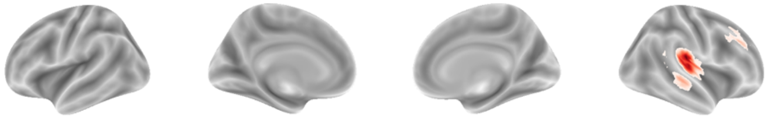

IC 65
(Most correlated with IC68 with r=0.88)

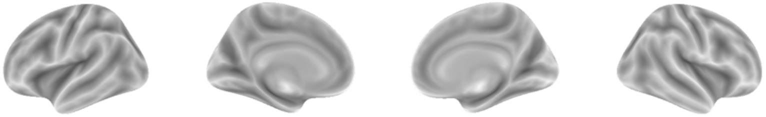

IC 66

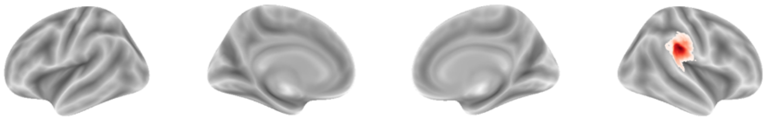

IC 67
(Most correlated with IC40 with r=0.78)

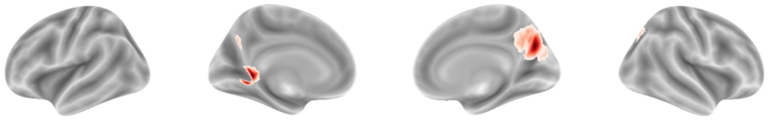

IC 68
(Most correlated with IC60 with r=0.88)

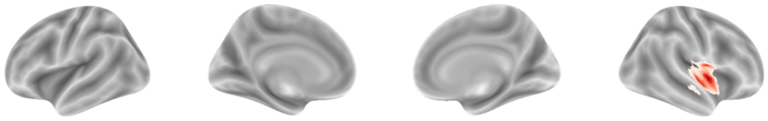

IC 69

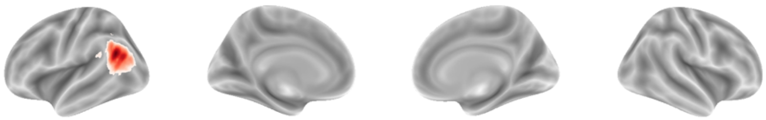

IC 70

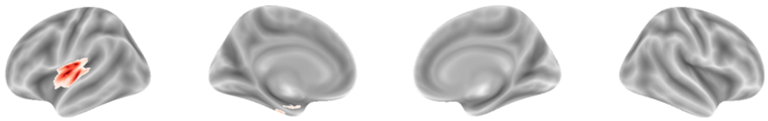

IC 71

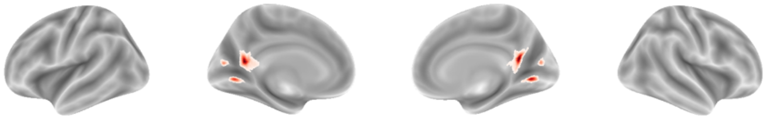

IC 72

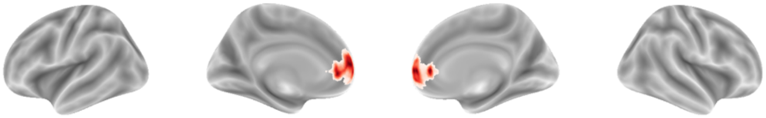

IC 73

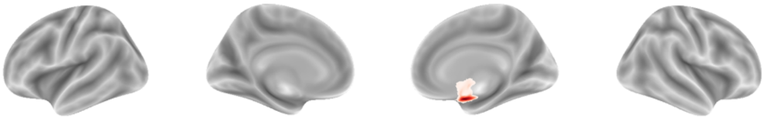

IC 74

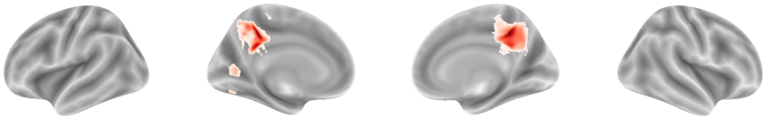

IC 75

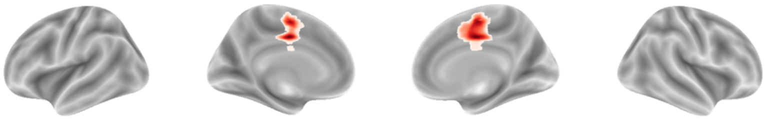

IC 76
(Most correlated with IC63 with r=0.79)

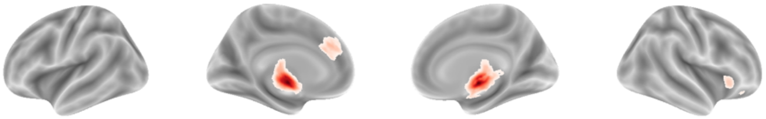

IC 77
(Most correlated with IC57 with r=0.84)

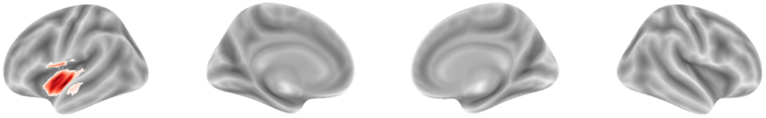

IC 78

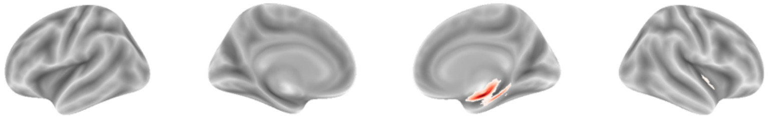

IC 79

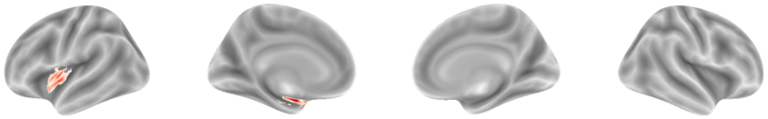

IC 80
(Most correlated with IC86 with r=0.78)

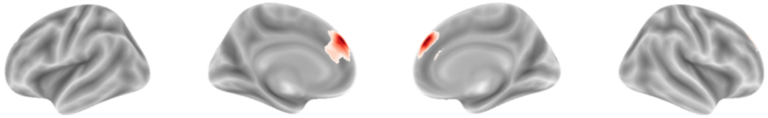

IC 81

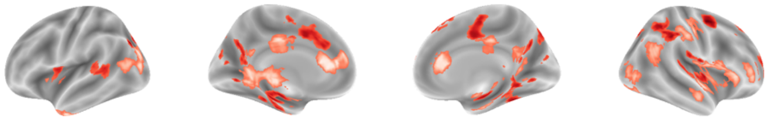

IC 82

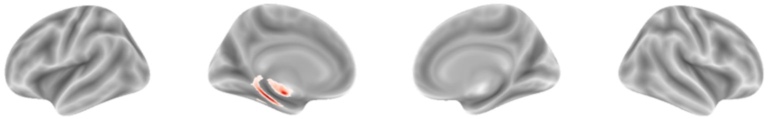

IC 83

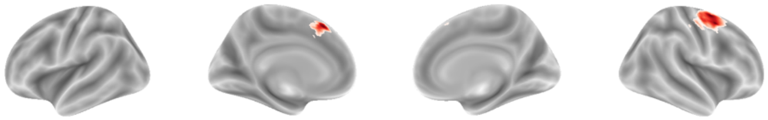

IC 84
(Most correlated with IC67 with r=0.82)

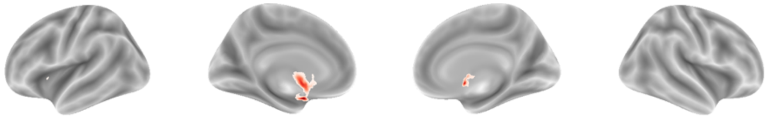

IC 85

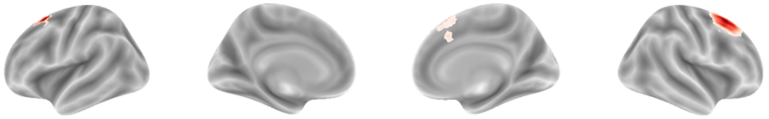

IC 86

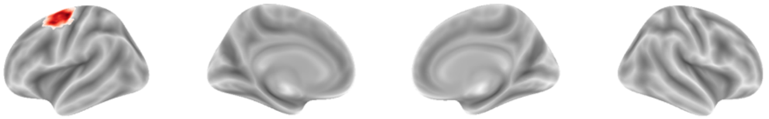

IC 87

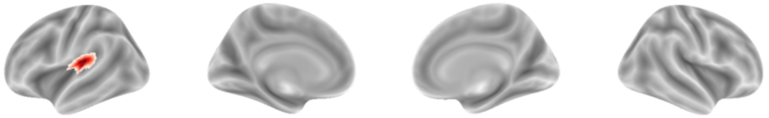

IC 88

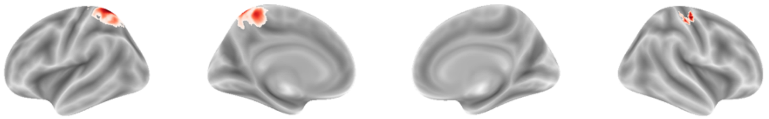

IC 89

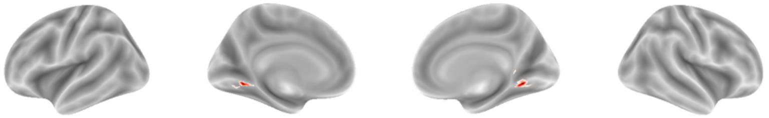

IC 90

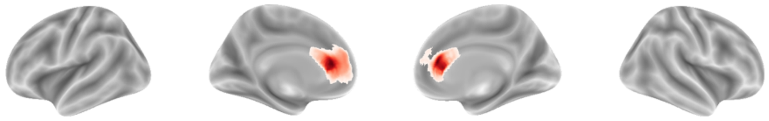

IC 91
(Most correlated with IC85 with r=0.87)

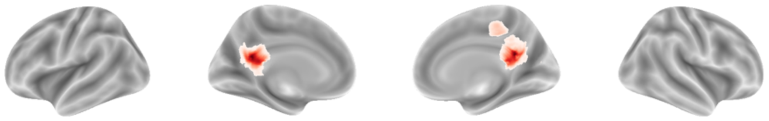

IC 92
(Most correlated with IC72 with r=0.83)

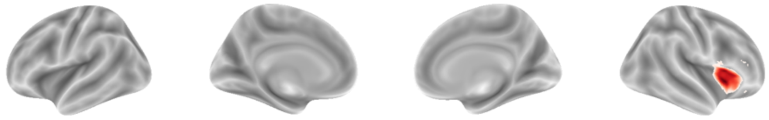

IC 93
(Most correlated with IC76 with r=0.81)

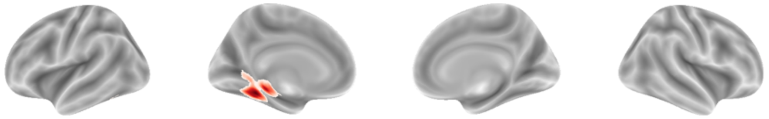

IC 94
(Most correlated with IC80 with r=0.85)

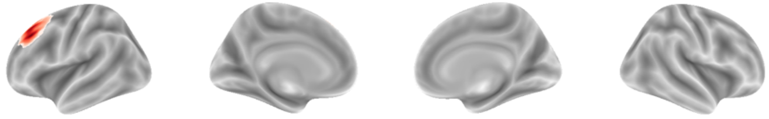

IC 95

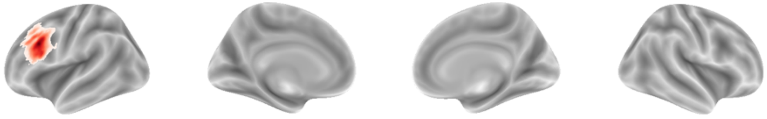

IC 96

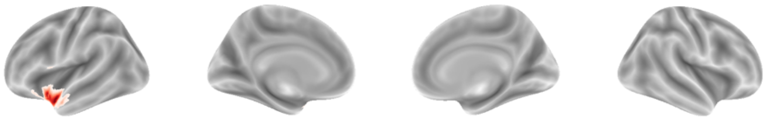

IC 97

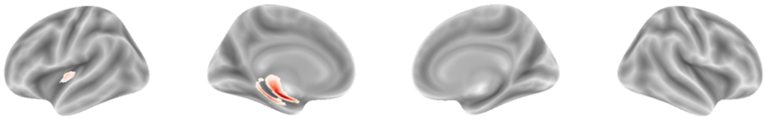

IC 98

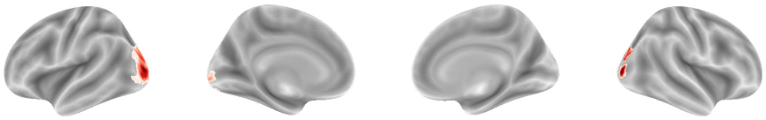

IC 99

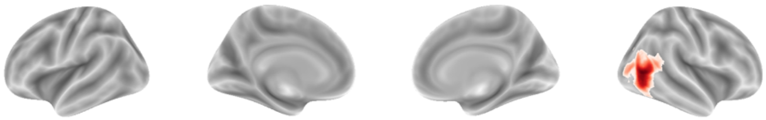

IC 100

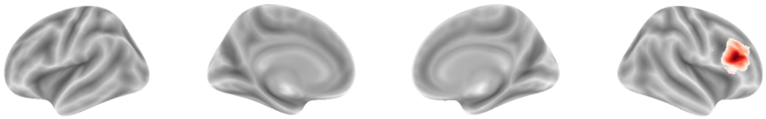

IC 101

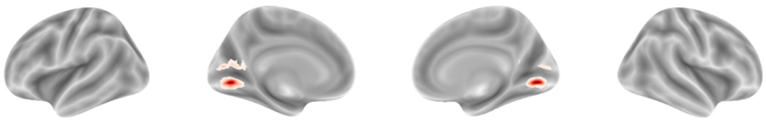

IC 102

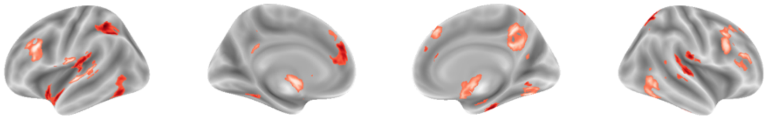

IC 103

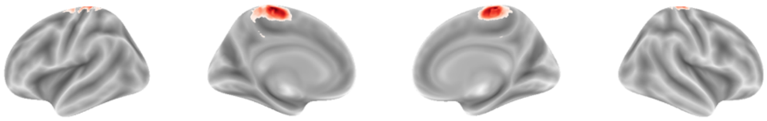

IC 104

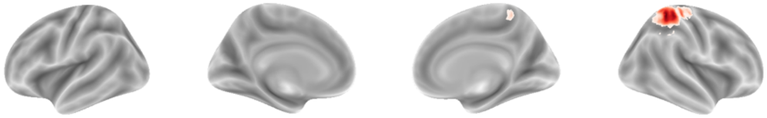

IC 105

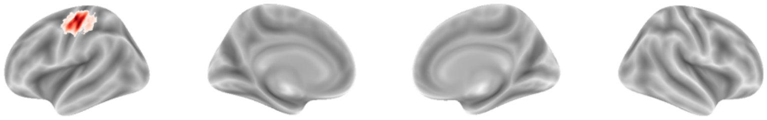

IC 106

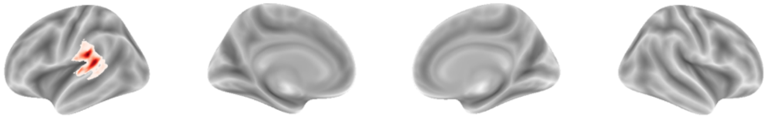

IC 107
(Most correlated with IC64 with r=0.64)

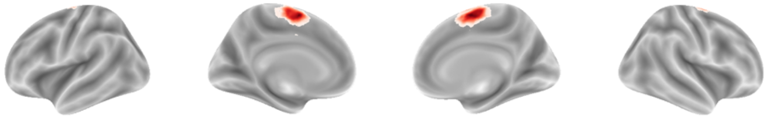

IC 108
(Most correlated with IC96 with r=0.70)

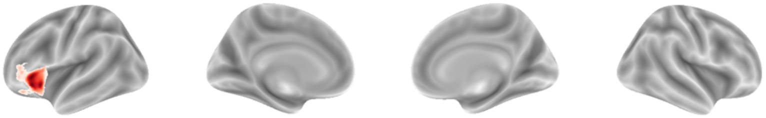

IC 109
(Most correlated with IC90 with r=0.68)

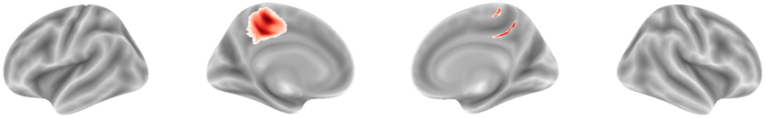

IC 110

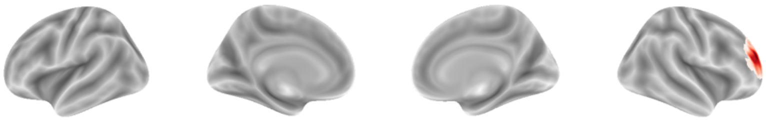

IC 111

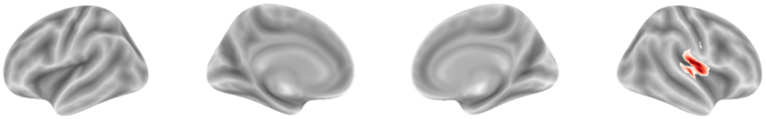

IC 112
(Most correlated with IC88 with r=0.47)

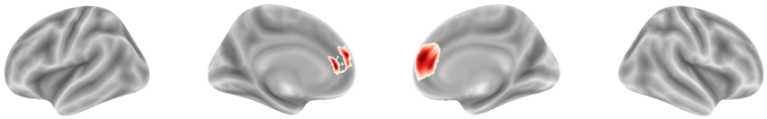

IC 113

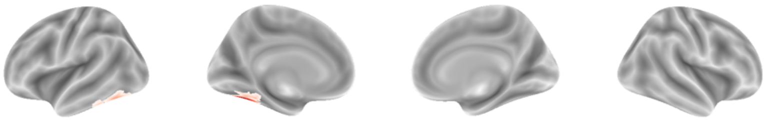

IC 114
(Most correlated with IC99 with r=0.48)

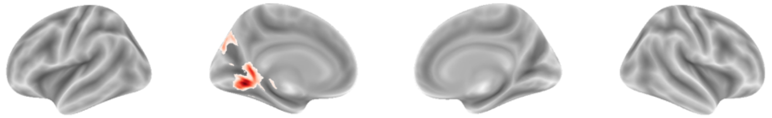

IC 115

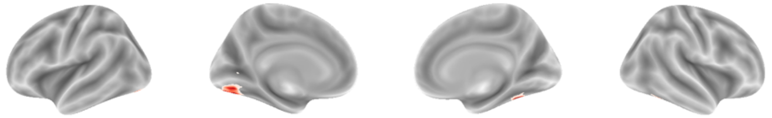

IC 116

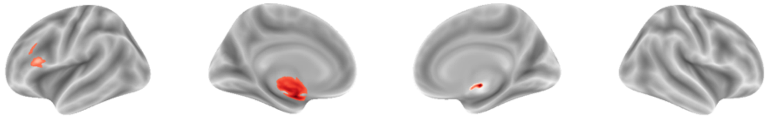

IC 117

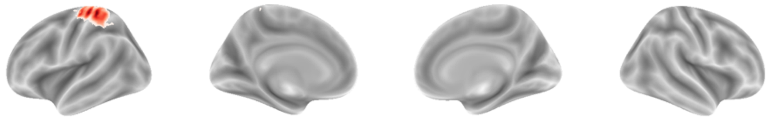

IC 118

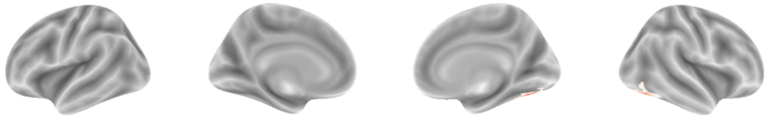

IC 119

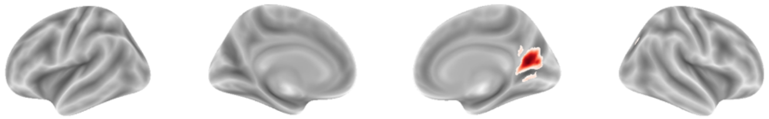

IC 120

(Most correlated with IC87 with r=0.75)

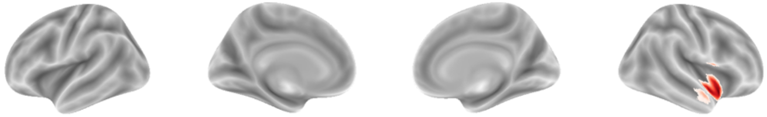

IC 121

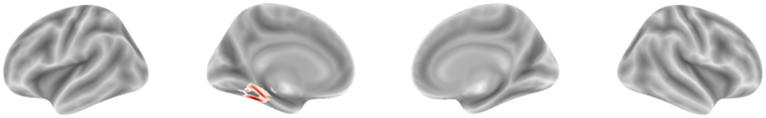

IC 122

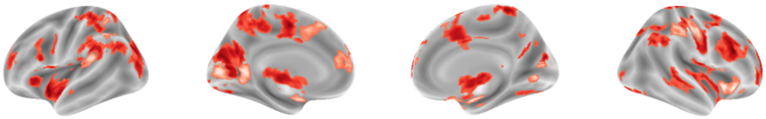

IC 123

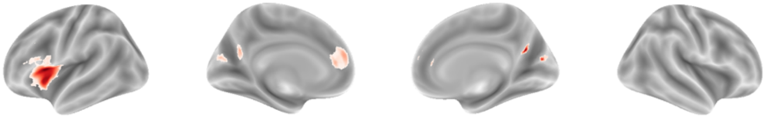

IC 124

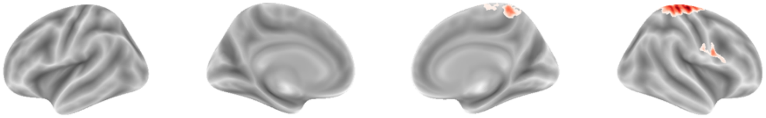

IC 125

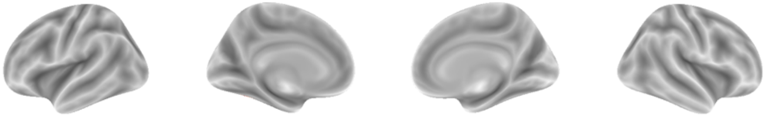

IC 126

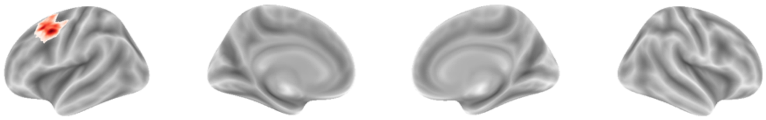

IC 127

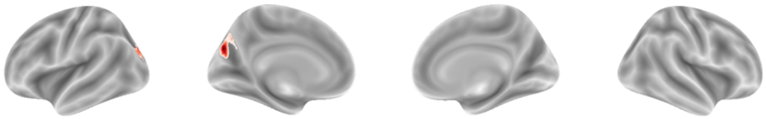

IC 128

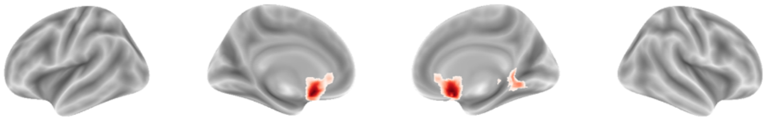

IC 129

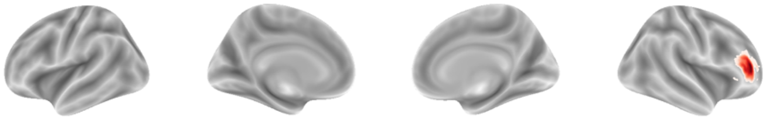

IC 130

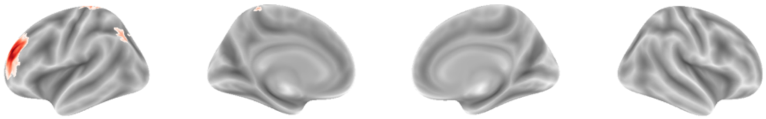

IC 131

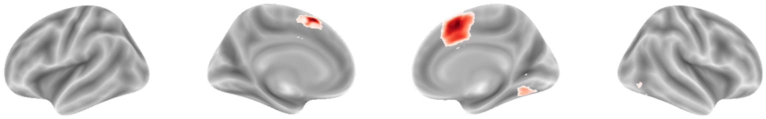

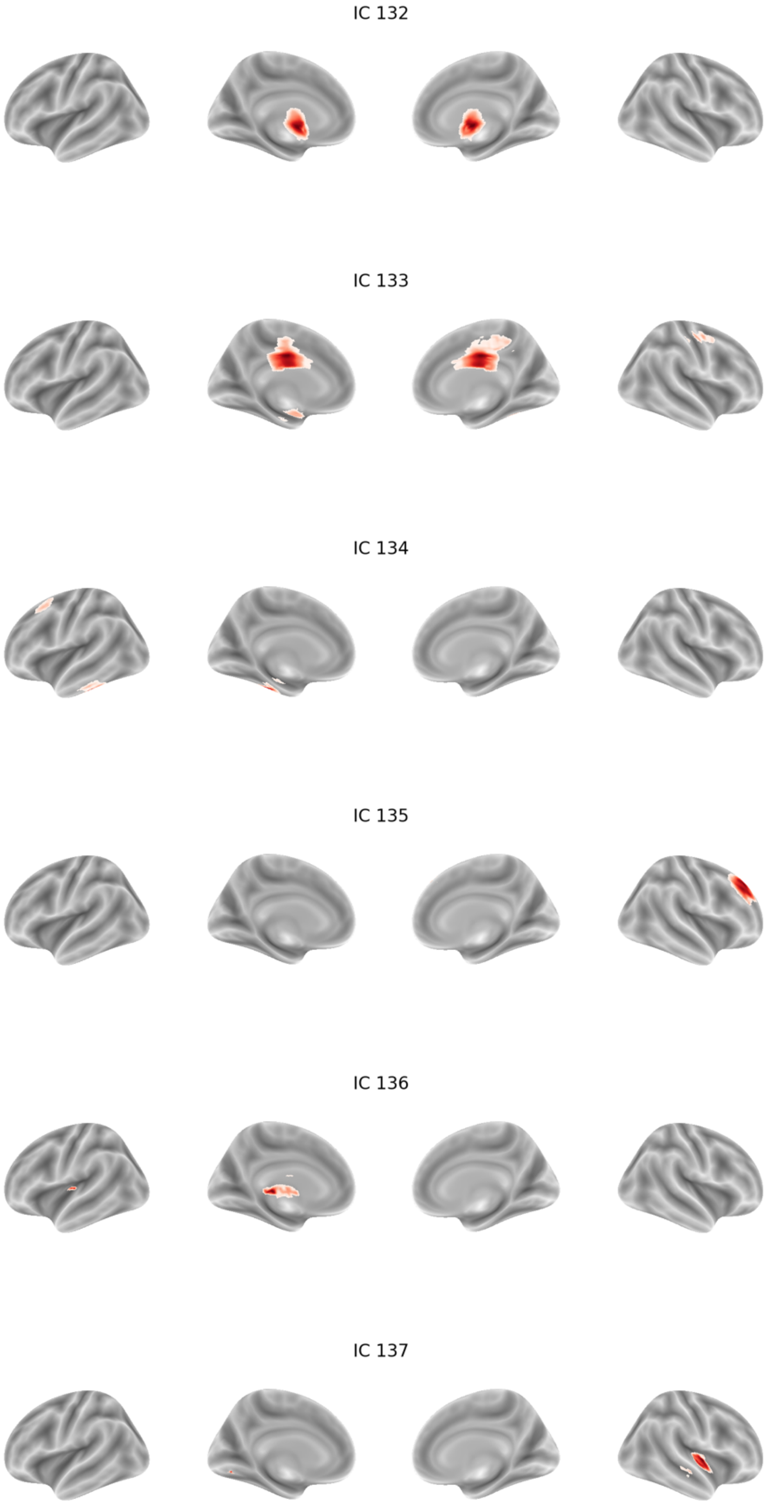
IC 132
IC 133
IC 134
IC 135
IC 136
IC 137

IC 138

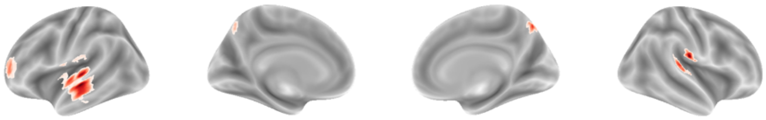

IC 139

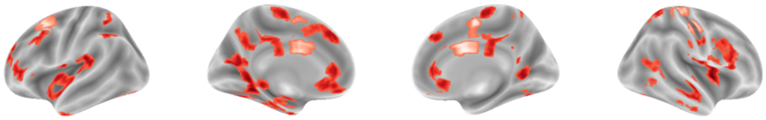

IC 140

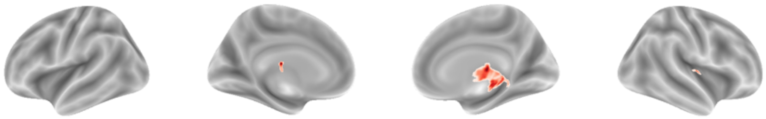

IC 141

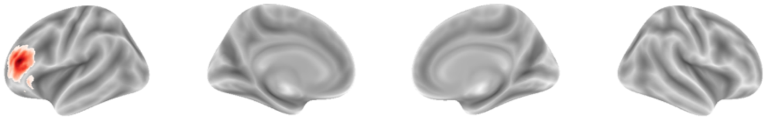

IC 142

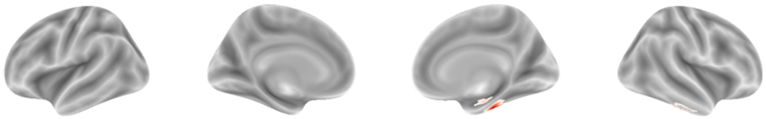

IC 143

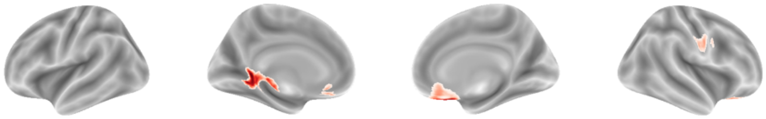

IC 144

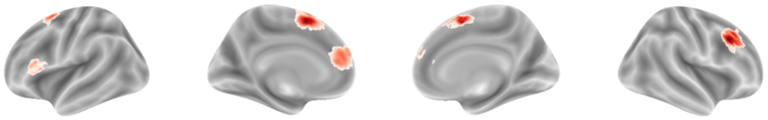

IC 145

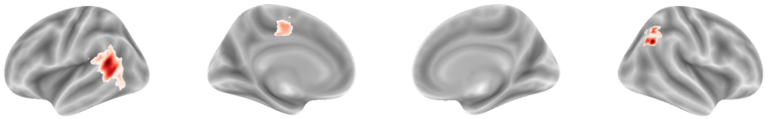

IC 146

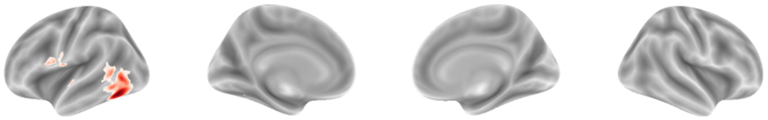

IC 147

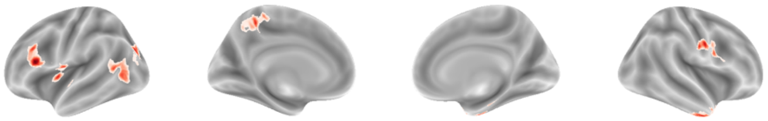

IC 148

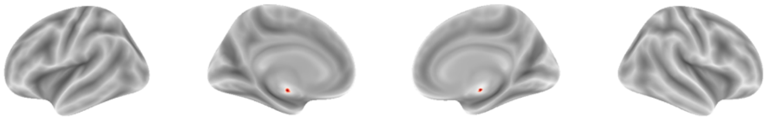

IC 149

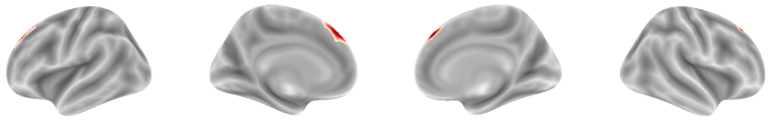

IC 150

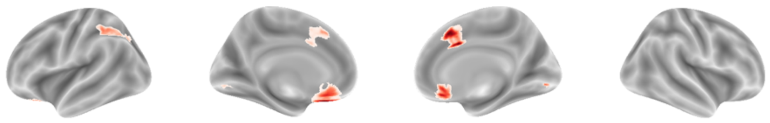

**Figure S4. 50-dimensional dendrogram.** *Dendrogram generated using hierarchical clustering of the IC time courses with 50 dimensions. A red border highlights ICs that show the highest spatial correlation with the 49 significant ICs identified from the 100-dimensional analysis in the Mantel test. The corresponding IC number from the 100-dimensional analysis is labeled below each highlighted IC.*

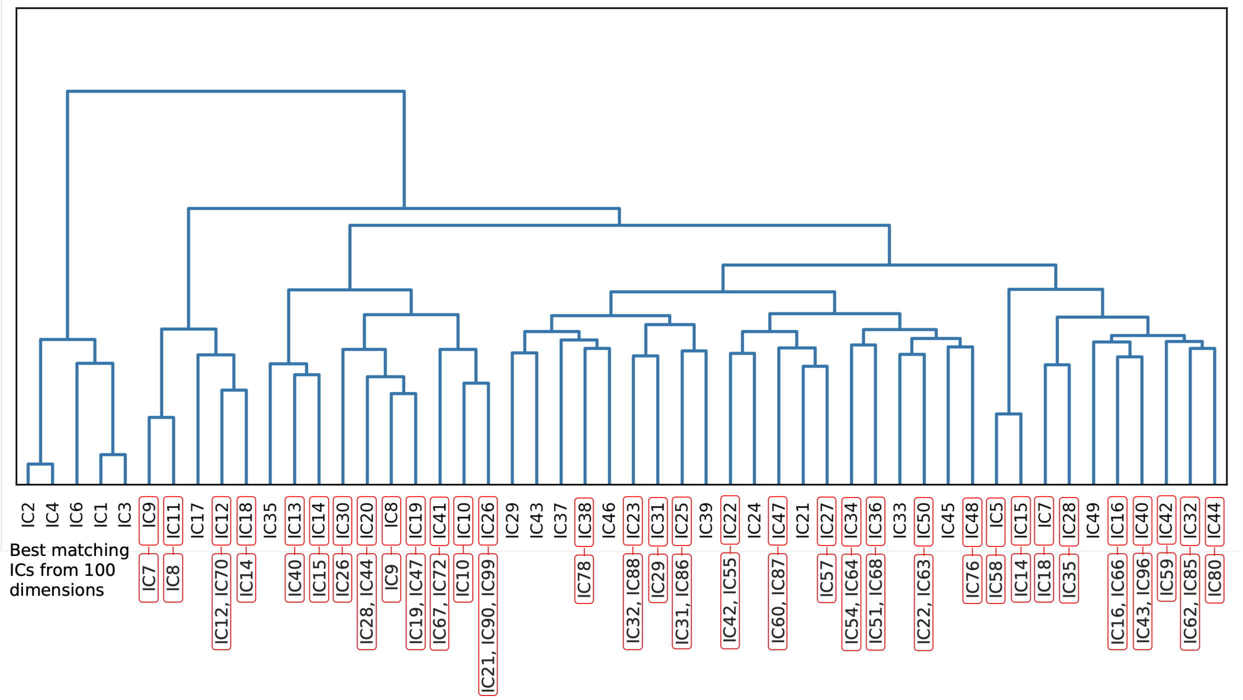

**Figure S5**. **100-dimensional dendrogram.** *Dendrogram generated using hierarchical clustering of the IC time courses with 100 dimensions.*

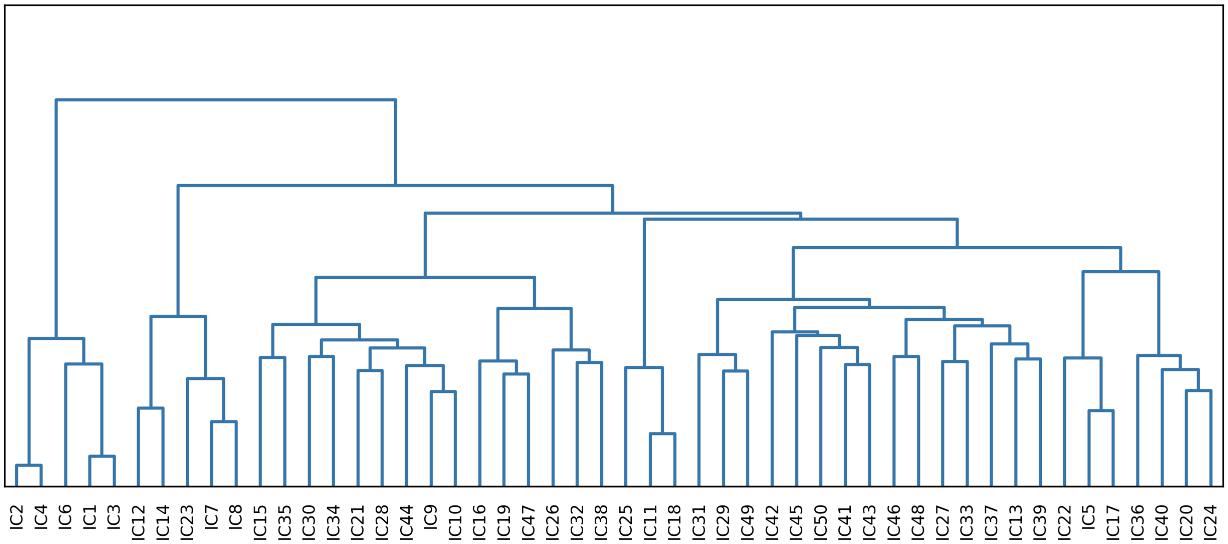


**Figure S6. 150-dimensional dendrogram.** *Dendrogram generated using hierarchical clustering of the IC time courses with 150 dimensions. A red border highlights ICs that show the highest spatial correlation with the 49 significant ICs identified from the 100-dimensional analysis in the Mantel test. The corresponding IC number from the 100-dimensional analysis is labeled below each highlighted IC.*

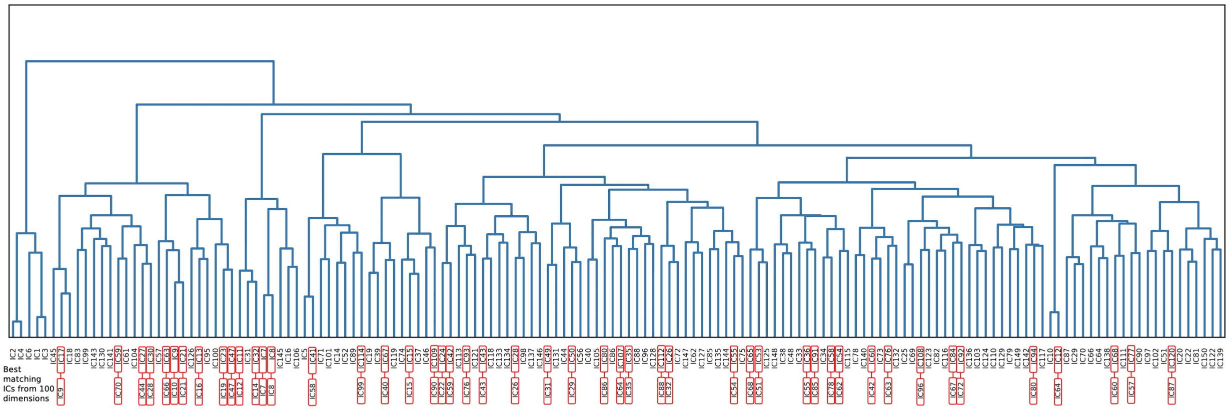


**Table S1. Network Information.** *Each row is the peak z-score voxel from statistically significant false discovery rate (FDR) corrected wellbeing-modulated independent component (IC) networks, as determined by a Mantel test (all FDR-corrected ps < 0.01). Rows are colour coded by co-activated networks or clades (see Fig. 1 and 3-5). The number of voxels is for the whole IC network (which might consist of multiple other regions). 'Decoding' provides the top 10 term-based meta-analysis associations at both the peak Montreal Neurological Institute (MNI) coordinate and whole clade for only cognitive/behavioural related terms (excluding, e.g., anatomical terms). Other abbreviations: A/P = anterior/posterior; I/S = inferior/superior; L/R = left/right; un/corr = un/corrected.*

| | | | | Peak MNI Coordinates | | | Decoding (N=10) | | | | | | |
|---|---|---|---|---|---|---|---|---|---|---|---|---|---|
| **Clade** | **IC** | **Region** | **Voxels** | **x (L/R)** | **y (A/P)** | **z (I/S)** | **Coordinate (z > 0)** | **Clade** | **ρ** | **Peak time** | **Z thresh** | **Uncorr p** | **FDR corr p** |
| 1 | 29 | Perigenual Anterior Cingulate Cortex | 241 | -2 | 28 | 29 | Pain, painful, gain, noxious, monetary, mood, nociceptive | Pain, painful, conflict, referential, self referential, ptsd, rehearsal, coordination, motor performance, default | 0.2793 | 5451 | 0.4 | 0.0003 | 0.0019 |
| 1 | 31 | Anterior Midcingulate Cortex | 100 | 2 | 3 | 44 | Pain, finger, movement, autonomic, movements, sensations, execution, tapping, index finger, target detection | | 0.2920 | 612 | 0.6 | 0.0004 | 0.0021 |
| 1 | 32 | Ventromedial Prefrontal Cortex | 90 | -2 | 45 | -3 | Default mode, default, mood, reward, valence, smoking, money, gambling, monetary, value | | 0.3107 | 2719 | 0.6 | 0.0022 | 0.0054 |
| 1 | 43 | Right Cerebellum | 184 | 23 | -75 | -24 | Interpersonal | | 0.2991 | 2241 | 0.3 | 0.0024 | 0.0056 |
| 1 | 50 | Right Cerebellum | 86 | 26 | -74 | -45 | Verbal working | | 0.3921 | 3411 | 0.6 | 0.0001 | 0.0009 |
| 1 | 51 | Left Cerebellum | 74 | -35 | -68 | -49 | Cognitive functions, suppressed, working memory, working | | 0.2675 | 2788 | 0.7 | 0.0004 | 0.0021 |
| 1 | 54 | Posterior Middle Cingulate Cortex | 309 | -2 | -26 | 49 | Foot, confidence, coordination | | 0.2557 | 4519 | 0.3 | 0.0010 | 0.0036 |
| 1 | 60 | Right Posterior Insula | 143 | 44 | -11 | 4 | Pain, auditory, music, discriminative, painful, musical, nociceptive, listening, audiovisual, noxious | | 0.3845 | 2445 | 0.5 | 0.0001 | 0.0009 |
| 1 | 64 | Dorsal Medial Frontal Cortex | 270 | 2 | -5 | 63 | Movements, movement, motor imagery, hand, sensorimotor, motor task, execution, finger, finger movements, imagery | | 0.2334 | 2598 | 0.4 | 0.0032 | 0.0066 |
| 1 | 68 | Left Cerebellum | 273 | -20 | -77 | -46 | Stimulation, pain, electrical, sensory, navigation, sensorimotor | | 0.2934 | 4314 | 0.3 | 0.0009 | 0.0036 |
| 1 | 78 | Right Cerebellum | 460 | 35 | -64 | -48 | Loop | | 0.2326 | 2782 | 0.2 | 0.0018 | 0.0046 |
| 1 | 80 | Left Superior Frontal Sulcus | 320 | -26 | 27 | 46 | Default, default network, autobiographical, default mode, real world, conceptual, episodic, autobiographical memory, memory, episodic memory | | 0.2167 | 2479 | 0.3 | 0.0044 | 0.0082 |
| 1 | 86 | Dorsal Medial Frontal Cortex | 146 | 2 | 43 | 43 | Inference, mind tom, arousal, emotional, social, tom | | 0.2574 | 5259 | 0.5 | 0.0001 | 0.0009 |
| 1 | 87 | Right Anterior Insula | 827 | 44 | -1 | -9 | Pain, intensity, aversive, taste, painful, articulatory, multisensory | | 0.2148 | 263 | 0.2 | 0.0027 | 0.0060 |
| 1 | 88 | Perigenual Anterior Cingulate Cortex | 61 | 8 | 47 | 14 | Default, beliefs, default mode | | 0.3378 | 1000 | 0.7 | 0.0001 | 0.0009 |
| 2 | 42 | Right Putamen | 1374 | 23 | -8 | 1 | Finger movements, finger tapping, movements, finger, movement, tapping, | Gain, | 0.2700 | 3852 | 0.1 | 0.0012 | 0.0036 |

| | | | | | | | | | | | | | |
|---|---|---|---|---|---|---|---|---|---|---|---|---|---|
| | | | | | | | sensorimotor, handed, sexual, execution | pain, reward, anticipation, painful, stop, monetary reward, monetary, sexual, mood | | | | | |
| 2 | 55 | Thalamus | 654 | 2 | -12 | 14 | Monetary reward, monetary, anticipation, retrieved, video clips, self reported | | 0.3812 | 391 | 0.1 | 0.0011 | 0.0036 |
| 2 | 57 | Left Posterior Insula | 346 | -44 | -7 | -6 | Experiences, pain, unfamiliar, acoustic | | 0.2478 | 51 | 0.3 | 0.0001 | 0.0009 |
| 2 | 63 | Ventral Tegmental Area | 274 | -2 | -17 | -6 | Reward, motivational, rewards, reinforcement learning, behaviours, reinforcement, motivation, addiction, choices, monetary | | 0.3229 | 1527 | 0.3 | 0.0003 | 0.0019 |
| 2 | 67 | Left Pallidum | 97 | -17 | -1 | -2 | Movement, movements, execution, coordination, tapping | | 0.2518 | 2198 | 0.6 | 0.0056 | 0.0098 |
| 2 | 72 | Right Anterior Insula | 1061 | 38 | 23 | 3 | Task, gain, calculation, mood, pain, choose, german, demands, tasks, orthographic | | 0.3142 | 4697 | 0.2 | 0.0004 | 0.0021 |
| 2 | 96 | Left Anterior Insula | 359 | -35 | 23 | 3 | Gain, phonological, language, orthographic, competing, words, heard, demands, pain | | 0.2432 | 1966 | 0.3 | 0.0025 | 0.0057 |
| 3 | 7 | Right Heschl's gyrus | 96 | 59 | -11 | 11 | Auditory, pitch, sounds, listening, touch, listened, noise, tactile, production, sound | Auditory, listening, speech, sounds, acoustic, sound, pitch, listened, speech perception, audiovisual | 0.3709 | 5111 | 0.3 | 0.0014 | 0.0039 |
| 3 | 8 | Left Supramarginal Gyrus | 418 | -50 | -46 | 25 | Mind tom, tom, phonological, social interaction, reading, speech production, language, listening, linguistic, theory mind | | 0.5245 | 1203 | 0.1 | 0.0001 | 0.0009 |
| 3 | 12 | Right Superior Temporal Gyrus | 176 | 53 | -46 | 16 | Theory mind, tom, mental states, mind tom, mind, motion, theory, watching, empathy, empathic | | 0.3690 | 2502 | 0.4 | 0.0012 | 0.0036 |
| 3 | 14 | Right Superior Temporal Gyrus | 281 | 53 | -23 | -3 | Listening, speech, auditory, language, sentences, comprehension, voice, linguistic, sentence, speech perception | | 0.3508 | 1223 | 0.5 | 0.0011 | 0.0036 |
| 4 | 18 | Right Precentral Gyrus | 459 | 47 | -16 | 40 | Sensorimotor, finger, speech production, vocal, production | Speech production, production, oral, sensorimotor, vocal, naming, articulatory, movement, movements, finger | 0.3753 | 9 | 0.2 | 0.0030 | 0.0063 |
| 5 | 15 | Left Precuneus | 241 | 2 | -63 | 54 | Spatial, calculation, judgement task, | Locations, | 0.3413 | 836 | 0.3 | 0.0020 | 0.0050 |

| | | | | | | | | | | | | | |
|---|---|---|---|---|---|---|---|---|---|---|---|---|---|
| | | | | | | | attention network, navigation | navigation, orienting, spatial attention, default, default mode, spatial, virtual, visual attention, location | | | | | |
| 5 | 40 | Left Precuneus | 619 | 14 | -65 | 31 | Retrieval, episodic, episodic memory, memory, recognition memory, memory retrieval, amnestic, items, autobiographical | | 0.3578 | 1641 | 0.2 | 0.0011 | 0.0036 |
| 6 | 22 | Right Lingual Gyrus | 475 | 20 | -60 | 2 | Implicit | Place, navigation, objects, visual, episodic, encoding, visual stream, episodic memory, object, autobiographical | 0.2430 | 3965 | 0.2 | 0.0029 | 0.0063 |
| 6 | 58 | Right Calcarine Sulcus | 55 | 14 | -77 | 14 | Sighted, visual, eye, eye fields, frontal eye, ageing, saccades, lexical | | 0.3463 | 1572 | 0.6 | 0.0007 | 0.0031 |
| 6 | 59 | Right Cerebellum | 179 | 32 | -53 | -26 | Motor performance, deficient, sensorimotor, motor imagery, execution, movement, movements, hand | | 0.2318 | 5230 | 0.5 | 0.0035 | 0.0068 |
| 6 | 62 | Left Cerebellum | 313 | -23 | -69 | -20 | Movements, movement, production, sensorimotor, finger tapping, pitch, tapping, rhythm, musicians, execution | | 0.2367 | 2653 | 0.3 | 0.0034 | 0.0068 |
| 6 | 76 | Left Parahippocampal Gyrus | 229 | -29 | -41 | -11 | Episodic, navigation, episodic memory, autobiographical, memory, retrieval, encoding, objects, place, construction | | 0.3027 | 1975 | 0.5 | 0.0035 | 0.0068 |
| 6 | 85 | Posterior Cingulate Cortex | 1279 | -2 | -52 | 15 | Default, default mode, autobiographical, episodic, memory retrieval, semantic memory, recollection, retrieval, default network, dmn | | 0.2723 | 3192 | 0.1 | 0.0040 | 0.0076 |
| 6 | 99 | Left Lingual Gyrus | 287 | -11 | -54 | -5 | Picture, navigation, words, semantic, sensorimotor | | 0.2383 | 1838 | 0.3 | 0.0014 | 0.0039 |
| 7 | 9 | Right Intraparietal Sulcus | 314 | 35 | -74 | 30 | Spatial, object, eye fields, virtual, objects, navigation, tasks, frontal eye, visuospatial, rotation | Tasks, working memory, working, task, visuospatial, attention, calculation, visual, spatial, attentional | 0.3618 | 2219 | 0.3 | 0.0053 | 0.0095 |
| 7 | 10 | Left Intraparietal Sulcus | 55 | -32 | -77 | 33 | Retrieval, memory, encoded, navigation, retrieved, episodic, object, episodic memory, ageing, visual | | 0.3372 | 3422 | 0.7 | 0.0001 | 0.0009 |
| 7 | 16 | Left Premotor Cortex | 270 | -47 | -3 | 41 | Working memory, working, speech production, saccades, phonological, production, eye, sensorimotor, rehearsal, language | | 0.5137 | 4027 | 0.2 | 0.0001 | 0.0009 |
| 7 | 19 | Right Dorsolateral Prefrontal Cortex | 127 | 44 | 12 | 32 | Mood, task, interference, demands, cognitive control, tasks, | | 0.3320 | 4498 | 0.7 | 0.0012 | 0.0036 |

| | | | | | | | | | | | | | |
|---|---|---|---|---|---|---|---|---|---|---|---|---|---|
| | | | | | | | stroop, response time, stroop task, ptsd | | | | | | |
| 7 | 21 | Left Intraparietal Sulcus | 281 | -35 | -56 | 48 | Memory, working memory, working, retrieval, recognition memory, word, calculation, memory retrieval, words, retrieved | | 0.2815 | 3645 | 0.4 | 0.0023 | 0.0055 |
| 7 | 26 | Bilateral Middle Temporal Visual Motion Area | 307 | -41 | -76 | -3 | Visual, objects, visual motion, motion, face recognition, object, face, negative neutral, orthographical, stream | | 0.3630 | 2096 | 0.2 | 0.0015 | 0.0041 |
| 7 | 28 | Right Intraparietal Sulcus | 622 | 38 | -53 | 45 | Tasks, working memory, working, task, memory, load, attentional, calculation, maintenance, demand | | 0.2677 | 1987 | 0.2 | 0.0048 | 0.0088 |
| 7 | 35 | Right Intraparietal Sulcus | 594 | 44 | -32 | 49 | Finger, tactile, planning, movements, hand, spatial, finger movements, tasks, sensorimotor, sequential | | 0.3522 | 879 | 0.3 | 0.0001 | 0.0009 |
| 7 | 44 | Right Intraparietal Sulcus | 153 | 23 | -60 | 54 | Eye movements, eye, visual, eye fields, attention network, spatial, movements, objects, attention, visually | | 0.2708 | 2384 | 0.4 | 0.0008 | 0.0034 |
| 7 | 47 | Right Frontal Eye Field | 55 | 41 | 2 | 47 | Eye fields, frontal eye, eye, spatial attention, eye movements, action, eye field, orienting, saccades, tasks | | 0.2371 | 481 | 0.7 | 0.0005 | 0.0023 |
| 7 | 66 | Left Dorsolateral Prefrontal Gyrus | 501 | -50 | 10 | 25 | Phonological, language, semantic, orthographic, words, languages, word, linguistic, reading, tasks | | 0.3355 | 801 | 0.2 | 0.0002 | 0.0015 |
| 7 | 70 | Middle Temporal Visual Motion Area | 235 | 44 | -64 | 11 | Motion, video clips, video, action observation, hands, actions, body, eye, eye movements, visual | | 0.2994 | 3546 | 0.5 | 0.0002 | 0.0015 |
| 7 | 90 | Midcingulate Cortex | 1008 | -11 | -41 | 48 | Self referential, referential, video clips | | 0.2271 | 519 | 0.2 | 0.0005 | 0.0023 |
